# Evaluating beta-tubulin variants as predictors of benzimidazole resistance across *Caenorhabditis* nematodes

**DOI:** 10.1101/2025.03.13.643047

**Authors:** Amanda O. Shaver, Ryan McKeown, Joyce M. Reyes Otero, J.B. Collins, Daniel W. Hogan, James S. Fraser, Stephen M. Dreyer, Erik J. Ragsdale, Erik C. Andersen

## Abstract

Benzimidazoles, a widely used class of anthelmintic drugs, target beta-tubulin, disrupt microtubule formation, and delay nematode development. In parasitic nematodes, mutations in beta-tubulin genes are predicted to inhibit benzimidazole binding and are associated with resistance. In the free-living nematode *Caenorhabditis elegans*, loss-of-function mutations in the beta-tubulin gene *ben-1* cause benzimidazole resistance. Although several beta-tubulin mutations serve as established markers of resistance, the prediction of the effects of novel variants in different nematode species remains challenging. Here, we identified novel beta-tubulin variants predicted to confer benzimidazole resistance across wild strains in three *Caenorhabditis* species: *C. elegans*, *Caenorhabditis briggsae*, and *Caenorhabditis tropicalis*. The three *Caenorhabditis* species are experimentally tractable, have characterized beta-tubulin gene complements, and defined natural niches, which allowed us to identify variants in beta-tubulin genes and test which variants are associated with resistance. We hypothesized that, if these species experienced similar selective pressures, they would evolve resistance to benzimidazoles by mutations in a beta-tubulin gene (*tbb-1*, *tbb-2*, *mec-7*, *tbb-4*, and *ben-1*). In the three *Caenorhabditis* species, we tested all strains harboring variants in the five conserved beta-tubulin genes for benzimidazole resistance. In *C. elegans*, we found that a heterogeneous set of variants in *ben-1* were associated with resistance. By contrast, only two variants in *C. briggsae ben-1* (W21stop and Q134H) were associated with resistance. *C. tropicalis* was distinct from the other two species, where no strains with variants in any beta-tubulin gene were resistant. We generated deletions of *ben-1* in *C. briggsae* and *C. tropicalis* and confirmed that loss of *ben-1* confers resistance in both species. Our findings reveal species-specific patterns of beta-tubulin-mediated benzimidazole resistance and emphasize that prediction of variants in beta-tubulin genes alone is not sufficient to predict resistance, especially across diverse nematode species.

**AUTHOR SUMMARY:** Mutations in beta-tubulin genes have been associated with benzimidazole resistance across nematode species, yet predicting novel resistance variants remains challenging. Using wild strains from three *Caenorhabditis* species, we identified strains with variants in beta-tubulin genes and tested each strain for benzimidazole resistance. In *C. elegans,* a diverse set of loss-of-function variants in *ben-1* were associated with resistance. Whereas in *C. briggsae*, only two *ben-1* alleles were associated with resistance, suggesting selection acts differently in this species despite a similar niche as *C. elegans*. *C. tropicalis* had no strains with beta-tubulin variants that were resistant. Our results highlight species-specific patterns of benzimidazole resistance.

## INTRODUCTION

Global control of parasitic nematode infections relies on the efficacy of a small arsenal of anthelmintic drugs, including benzimidazoles (BZs) [1]. BZs are a widely used class of anthelmintic drugs that inhibit the polymerization of microtubules [2–4] and delay nematode development [5]. Although BZs are essential to human and veterinary health, resistance is prominent in parasitic nematode populations [6,7]. In clade V nematodes (*e.g.*, *Ancylostoma caninum*, *Ancylostoma duodenale*, *Caenorhabditis elegans*, *Haemonchus contortus*, *Necator americanus*, *Teladorsagia circumcincta*, and *Trichostrongylus colubriformis*), BZ resistance is associated with mutations in beta-tubulin genes [8–16].

Among parasitic nematodes, BZ resistance has been best characterized in the small ruminant parasite *H. contortus* [17]. The *H. contortus* genome contains four genes that encode beta-tubulins (*tbb-isotype-1*, *tbb-isotype-2*, *tbb-isotype-3*, and *tbb-isotype-4*), where each encoded protein has a phenylalanine at position 200, which is thought to confer binding to BZs [18]. In field populations of *H. contortus*, BZ resistance has been associated historically with three canonical missense variants (F167Y, E198A, and F200Y) in *tbb-isotype-1* [19–22]. Recently, additional novel missense variants in *tbb-isotype-1* have been associated with *H. contortus* BZ resistance (*e.g.*, E198I, E198K, E198T, and E198stop) [23] These newly described missense variants in *tbb-isotype-1* are hypothesized to disrupt BZ binding, without causing loss-of-function (LoF). Given the high expression of *tbb-isotype-1*, it is likely an essential beta-tubulin in *H. contortus [18]*. By contrast, *tbb-isotype-2* LoF alleles have been identified in highly resistant *H. contortus* populations [24], indicating that it is not an essential gene and does not confer resistance on its own because variants are not observed in resistant *H. contortus* populations [18]. The other two beta-tubulin genes are expressed at low levels and have not been associated with BZ resistance [18]. The beta-tubulin genes and alleles involved in BZ resistance suggest that specific variants that prevent or reduce BZ binding can be tolerated, whereas complete loss of essential beta-tubulin genes cannot.

Unlike *H. contortus*, the free-living model nematode species *C. elegans* has six beta-tubulin genes (*tbb-1*, *tbb-2*, *mec-7*, *tbb-4*, *ben-1*, and *tbb-6*) [25]. LoF mutations in *ben-1*, an ortholog of *H. contortus tbb-isotype-1* and *tbb-isotype-2* [18], confer BZ resistance [26]. Additionally, a heterogeneous set of LoF variants in *ben-1* identified in *C. elegans* wild strains cause natural BZ resistance in this species [14]. The ability to maintain wild-type growth despite the loss of *ben-1* is likely explained by functional redundancy among beta-tubulin genes [27]. Previous studies have demonstrated that *Cel-tbb-1* and *Cel-tbb-2* act redundantly for viability [28–30] and for movement, body morphology, and growth [31]. In *C. elegans*, *tbb-1* and *tbb-2* are the most broadly and highly expressed beta-tubulin genes, whereas *ben-1* expression is restricted largely to neurons, specifically cholinergic and glutamatergic neurons [25,32]. Consistent with this pattern, cell-specific rescue experiments demonstrated that *ben-1* acts in cholinergic and GABAergic neurons to confer BZ susceptibility, suggesting that these neurons are key sites of BZ action [33]. Of the six *C. elegans* beta-tubulins, only MEC-7, TBB-4, and BEN-1 contain a phenylalanine at position 200, the residue associated with BZ binding in *H. contortus*, whereas TBB-1 and TBB-2 contain tyrosine at this position and are predicted not to bind BZs. Together, these findings illustrate how loss of a *C. elegans* beta-tubulin that can bind BZs causes resistance because other beta-tubulin genes are functionally redundant for viability and do not bind BZs.

The recurrent association of beta-tubulin variants with BZ resistance in clade V parasitic and free-living nematodes suggests the hypothesis that resistance might be predictable across species. However, the reliability of beta-tubulin variants as predictors of BZ resistance across nematode species is unknown. This hypothesis is difficult to test directly in parasitic nematode species because of their host-dependent life cycles, poorly annotated reference genomes, and limited molecular and genetic tools [34,35]. By contrast, the availability of high-quality genomic data for hundreds of wild strains [36] and the laboratory tractability of the free-living *Caenorhabditis* nematode species, *C. elegans*, *Caenorhabditis briggsae*, and *Caenorhabditis tropicalis*, provide an opportunity to test predictions of beta-tubulin mediated BZ resistance in the *Caenorhabditis* genus. Establishing and interrogating patterns of repeated evolution in these three free-living species would provide a mechanistic framework to anticipate resistance-associated variants in beta-tubulin genes across other clade V nematode species, including parasitic nematodes where direct experimental validation of resistance mechanisms remains a challenge.

Using the global natural diversity of *C. elegans*, *C. briggsae*, and *C. tropicalis*, we assessed variation in the beta-tubulin genes *tbb-1*, *tbb-2*, *mec-7*, *tbb-4*, and *ben-1* and identified high-impact variants (*i.e.*, single nucleotide variants (SNVs), small insertions or deletions (INDELs), and structural variants (SVs) predicted to disrupt beta-tubulin function and confer BZ resistance. Because *ben-1* is the primary driver of BZ resistance in *C. elegans*, we used an established high-throughput larval development assay (HTLDA) to expose strains with novel *ben-1* variants to the highly used BZ, albendazole (ABZ). In *C. elegans*, strains harboring seven of nine novel *Cel-ben-1* variants were resistant to ABZ, adding more alleles hypothesized to confer resistance in this species. By contrast, strains harboring only two of the eight unique *Cbr-ben-1* variants (W21stop and Q134H) were resistant to ABZ. *C. tropicalis* was distinct from the other two species, where no strains with variants in *ben-1* were resistant to ABZ. To validate the roles of *Cbr-ben-1* and *Ctr-ben-1* in BZ resistance, we generated deletion alleles and confirmed that loss of *ben-1* confers resistance in both *C. briggsae* and *C. tropicalis*. Fecundity assays showed that a loss of *Cbr-ben-1* did not affect *C. briggsae* fitness, whereas a deletion of *Ctr-ben-1* significantly reduced fecundity in *C. tropicalis*. Because *ben-1* variants could not always predict ABZ resistance, we tested whether strains harboring high-impact variants in the other four conserved beta-tubulin genes were resistant to ABZ. Strains with variants in *tbb-1*, *tbb-2*, *mec-7*, and *tbb-4* were not resistant to ABZ in any of the three *Caenorhabditis* species, indicating that *ben-1* is the primary beta-tubulin that confers resistance. The species-specific patterns of beta-tubulin-mediated BZ resistance might reflect species-specific selection pressures, such as exposure to natural BZs, and can shape the evolution of beta-tubulin-mediated resistance. The identification of alleles associated with BZ resistance in experimentally tractable species establishes a framework to predict beta-tubulin variants associated with resistance in other clade V nematodes.

## RESULTS

### Strains with variants or expression differences in *ben-1* were predicted to be ABZ resistant

To predict ABZ resistant strains in the three *Caenorhabditis* species, we identified high-impact variants (SNVs, INDELs, or SVs) in *ben-1* (see *Materials and Methods*). In a set of 611 *C. elegans* wild strains [36], we identified 65 strains with 33 unique high-impact variants in *ben-1* (**S1 Table**). Of the 33 variants, 24 were previously phenotyped and 20 were associated with ABZ resistance (**S1A Fig.**) [14,37]. In 641 *C. briggsae* wild strains, 22 strains with eight unique high-impact variants in *ben-1* were identified (**S2 Table**). In a set of 518 *C. tropicalis* wild strains, only two strains with unique high-impact variants in *ben-1* were identified (**S3 Table**).

Next, because low *ben-1* expression was previously correlated with ABZ resistance in *C. elegans* wild strains [38], we evaluated whether *ben-1* expression levels were predictive of ABZ responses in *C. elegans.* We assessed the relationship between the expression of *ben-1* [38] and ABZ responses in 180 *C. elegans* wild strains [14,37] (*p*-value = 5.16e-16, *r*^2^= 0.344). We hypothesized that low *ben-1* expression could contribute to ABZ resistance in *C. briggsae* and *C. tropicalis*, but expression data from wild strains in these species have yet to be collected.

Altogether, we identified nine novel variants in *Cel-ben-1* (E3stop, Y50C, P80S, VDN113N, Q131L, frameshift 319, frameshift 368, stop445S, and a duplication) (**S1 Table**), one strain with low *Cel-ben-1* expression, eight variants in *Cbr-ben-1* (W21stop, V91I, Q94K, D128E, Q134H, S218L, M299V, and R359H) (**S2 Table**), and two variants in *Ctr-ben-1* (P80T and R121Q) (**S3 Table**) to test for ABZ resistance. In addition to these selected strains, we also selected *C. briggsae* and *C. tropicalis* strains that lacked high-impact variants in beta-tubulin genes but were closely related to strains with variants (**S2 Fig., S3 Fig., and S4 Table**). The strains with no high-impact variants in any beta-tubulin gene might control for differences in wild strain genetic backgrounds.

### High-throughput larval development assays (HTLDAs) reveal species-specific associations between *ben-1* variation and ABZ resistance

Nematodes grow longer as they progress through development, and BZs slow this progression [14–16,27,37]. A longer animal length (*i.e.*, larger animals) corresponds to increased ABZ resistance, and a shorter animal length (*i.e.*, smaller animals) corresponds to increased ABZ sensitivity. To evaluate the effects of ABZ on animal length (a proxy for development), we used image-based HTLDAs to expose all *C. elegans*, *C. briggsae*, and *C. tropicalis* wild strains with novel high-impact variants in *ben-1* or low *ben-1* expression (*i.e.*, predicted resistant strains) to control (DMSO) (**S4 Fig., S5 Fig., and S6 Fig.**) and drug (ABZ) conditions (**S7 Fig., S8 Fig., and S9 Fig.**) (see *Materials and Methods*). The assay included 48 replicates per strain with five to 30 animals per replicate in DMSO or ABZ conditions. The reported nematode length of each strain is the difference between animal lengths in DMSO and ABZ.

To define the role of *ben-1* in BZ response and classify wild strains as BZ resistant, we first obtained or created *ben-1* deletions in the reference strains for each of the three *Caenorhabditis* species. In *C. elegans*, the strain with a *ben-1* LoF variant in the N2 strain background (ECA882) has been used as an ABZ-resistant control previously [14,27,37]. For *C. briggsae* and *C. tropicalis*, we used CRISPR-Cas9 genome editing to generate two independent *ben-1* deletion alleles per species (see *Materials and Methods*). In *C. briggsae*, deletions of *Cbr-ben-1* were created in the AF16 reference strain background (ECA3953 and ECA3954) (**S10 Fig. and S5 Table**). In *C. tropicalis*, deletions of *Ctr-ben-1* were created in the NIC58 reference strain background (ECA4247 and ECA4248) (**S11 Fig. and S5 Table**). All deletions of *ben-1* in each of the three *Caenorhabditis* species conferred high levels of ABZ resistance (**S12 Fig. and S13 Fig.**). Wild strains were then classified as resistant using a threshold where the median animal lengths after ABZ exposure were no more than two SD below the median animal length of the species-specific *ben-1* deletion strain.

In *C. elegans*, seven of the nine strains with *ben-1* variants showed minimal developmental delays after exposure to ABZ, a phenotype similar to loss of *ben-1*, and were classified as resistant (**Fig. 1**). The two remaining strains with *ben-1* variants (P80S, stop445S) and the strain with low *ben-1* expression were not strongly resistant to ABZ. The P80S variant might partially alter *ben-1* function, causing a moderate resistance phenotype. In stop445S, the normal stop codon was replaced with a serine. This variant likely does not affect *ben-1* function because position 445 is at the end of the BEN-1 protein, and only four amino acid residues (NRKL) are added beyond the wild-type stop codon. Finally, the strain with low *ben-1* expression and no high-impact variants in *ben-1* (JU1581) was sensitive to ABZ, indicating that the selected threshold of *ben-1* expression (≥3.75 TPM) still retained strains with adequate *ben-1* function. Overall, natural allelic variation in *Cel-ben-1* is associated with BZ resistance, recapitulating previous findings that BZ resistance is associated with a diverse set of LoF variants in *Cel-ben-1* [14,27,31,37].

**Fig 1.**
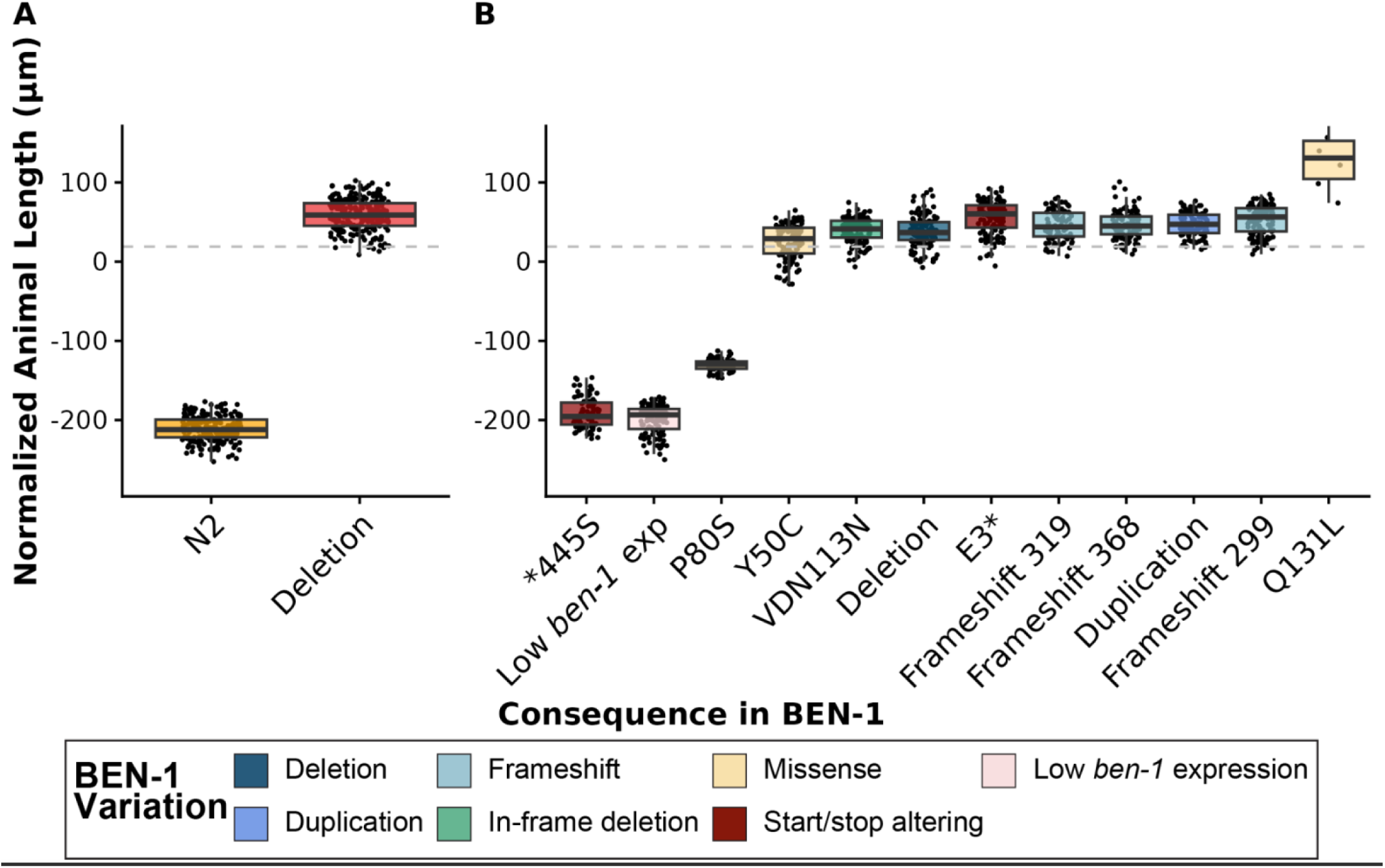
High-throughput larval development assays in the presence of albendazole for *C. elegans* strains with unique high-impact variants in BEN-1. The regressed median animal length values for populations of nematodes grown in 30 μM albendazole (ABZ) are shown on the y-axis. Each point represents the normalized median animal length value of a well containing approximately 5-30 animals. Strains are sorted by their relative resistance to ABZ based on median animal length. Data are shown as Tukey box plots with the median as a solid horizontal line, and the top and bottom of the box representing the 75th and 25th quartiles, respectively. The top whisker is extended to the maximum point that is within the 1.5 interquartile range from the 75th quartile. The bottom whisker is extended to the minimum point that is within the 1.5 interquartile range from the 25th quartile. The gray dashed line marks the *C. elegans* resistance threshold, defined as two standard deviations below the mean of the *ben-1* deletion strain in the N2 reference strain background. Results for **(A)** the N2 reference strain (orange) and a strain with a *ben-1* deletion in the N2 background (red), and **(B)** all wild *C. elegans* strains with unique high-impact variants in *ben-1* are sorted by their relative resistance to ABZ based on median animal length. Wild *C. elegans* strains are colored by beta-tubulin variant status.

In *C. briggsae*, we performed HTLDAs on the AF16 reference strain (ABZ sensitive), two *ben-1* deletion strains (ECA3953 and ECA3954) (ABZ resistant), 11 strains with eight unique variants in *ben-1*, and 13 strains genetically related to strains with *ben-1* variants in DMSO (control) (**S5 Fig.**) and ABZ conditions (**Fig. 2, S8 Fig., and S12 Fig.**). Of the 11 strains with *Cbr-ben-1* variants, only two strains with unique variants in BEN-1 (Q134H and W21stop) were ABZ resistant (**Fig. 2**). The Q134H amino acid change has been associated with ABZ resistance in *A. caninum* and validated in *C. elegans* [11]. An early stop gain at position 21 is predicted to cause the premature termination of protein synthesis and LoF. Of all the assayed strains with variants in BEN-1, six (V91I, Q94K, D128E, S218L, M299V, and R359H) were not resistant to ABZ. Additionally, eleven *C. briggsae* strains were exposed to a maximum concentration of 120 μM ABZ, where only two strains with the variants Q134H and W21stop were resistant at these higher concentrations (**S14 Fig.**). Overall, because only one of the seven missense variants was associated with ABZ resistance, we could not reliably predict *C. briggsae* ABZ resistance based on the presence of a missense variant alone. To evaluate the potential fitness consequences of *ben-1* LoF alleles, fecundity assays were performed between the AF16 reference strain and the two strains with a loss of *ben-1* in the AF16 background (ECA3953 and ECA3954) (**S15A Fig. and S6 Table**). We found no significant differences in fecundity between the three strains, indicating that a loss of *ben-1* does not affect fitness in this species. These results indicate that, although a loss of *Cbr-ben-1* confers ABZ resistance with no fitness detriment, ABZ resistance associated with *Cbr-ben-1* variants is uncommon.

**Fig 2.**
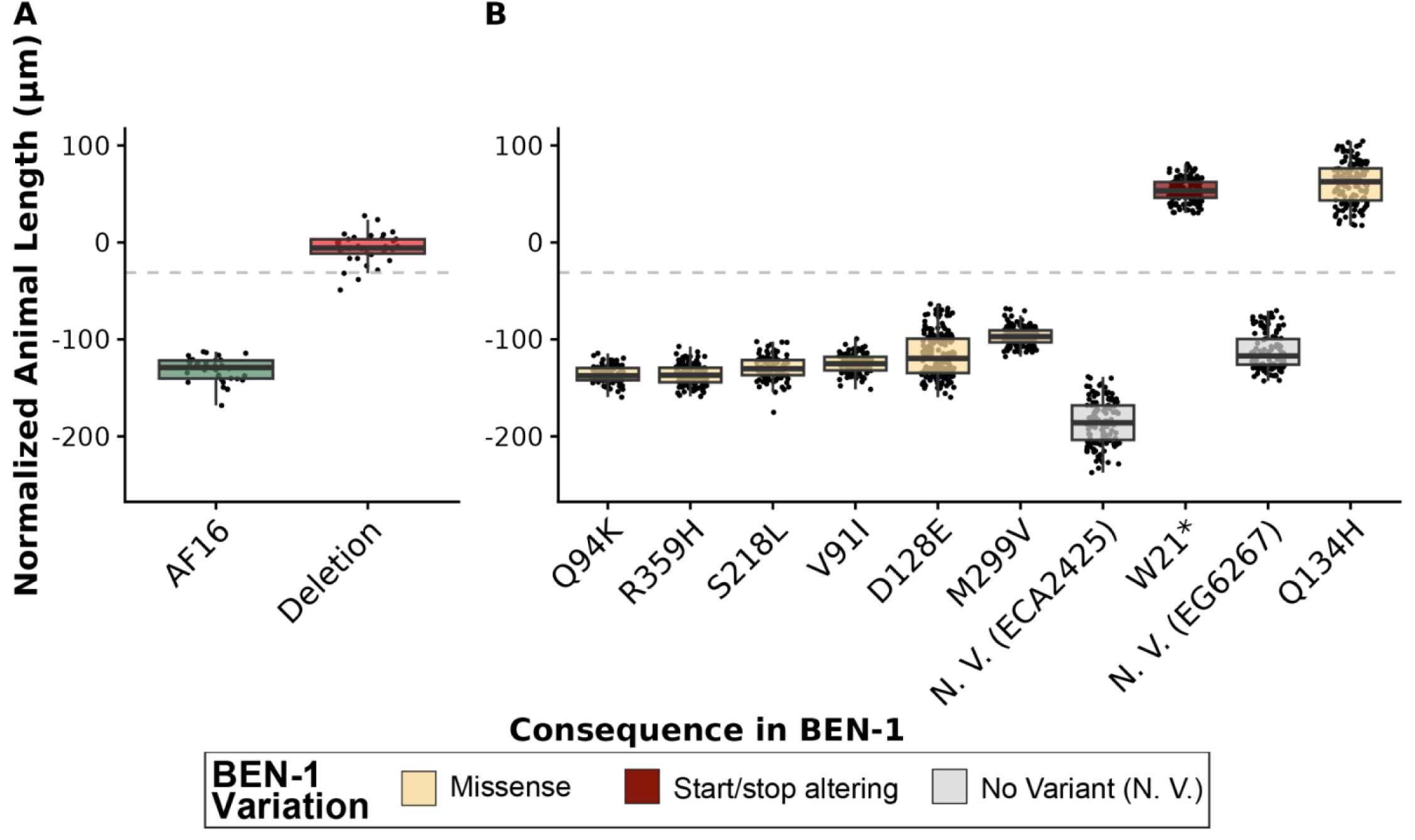
High-throughput larval development assays in the presence of albendazole for *C. briggsae* strains with unique high-impact variants in BEN-1. The regressed median animal length values for populations of nematodes grown in 30 μM albendazole (ABZ) are shown on the y-axis. Each point represents the normalized median animal length value of a well containing approximately 5-30 animals. Data are shown as Tukey box plots with the median as a solid horizontal line, and the top and bottom of the box representing the 75th and 25th quartiles, respectively. The top whisker is extended to the maximum point that is within the 1.5 interquartile range from the 75th quartile. The bottom whisker is extended to the minimum point that is within the 1.5 interquartile range from the 25th quartile. The gray dashed line marks the *C. briggsae* resistance threshold, defined as two standard deviations below the mean of the *ben-1* deletion strain in the AF16 reference strain background. Results for **(A)** the AF16 reference strain (green) and a strain with a *ben-1* deletion in the AF16 background (red), and **(B)** all wild *C. briggsae* strains with unique high-impact variants in *ben-1* are sorted by their relative resistance to ABZ based on median animal length. No variant (N. V.) strains (gray) paired with strains that have a high-impact variant in *ben-1* that pass the resistance threshold are shown alongside each corresponding strain with a high-impact variant in *ben-1*. Wild *C. briggsae* strains are colored by beta-tubulin variant status.

In *C. tropicalis*, we performed HTLDAs on the reference strain NIC58 (ABZ sensitive), two independent deletions of *Ctr-ben-1* (ECA4247 and ECA4248) (ABZ resistant), two *C. tropicalis* strains with variants in *ben-1* (P80T and R121Q), and two strains genetically related to *ben-1* variant strains in DMSO (**S6 Fig.**) and ABZ conditions (**Fig. 3 and S9 Fig.**). We found that all *C. tropicalis* wild strains displayed ABZ sensitivity, indicating that P80T and R121Q are not associated with ABZ resistance (**Fig. 3**). Additionally, all tested wild strains were exposed to a maximum concentration of 120 μM ABZ, and none of the strains displayed resistance at these higher concentrations (**S16 Fig.**). Finally, to evaluate the potential fitness consequences caused by a loss of *ben-1*, fecundity assays were performed between the NIC58 reference strain and the two strains with a loss of *ben-1* in the NIC58 background (ECA4247 and ECA4248) (**S15B Fig. and S7 Table**). We found that a deletion of *ben-1* caused a significant reduction in fecundity in *C. tropicalis*. The fitness defect caused by a loss of *ben-1* could explain the absence of naturally occurring *ben-1* variants in wild *C. tropicalis* strains, suggesting that *C. tropicalis* is unlikely to acquire natural *ben-1* variants that confer ABZ resistance without strong selection. If ABZ resistance is present in *C. tropicalis* wild strains, its genetic basis remains unknown and involves factors distinct from *ben-1*.

**Fig 3.**
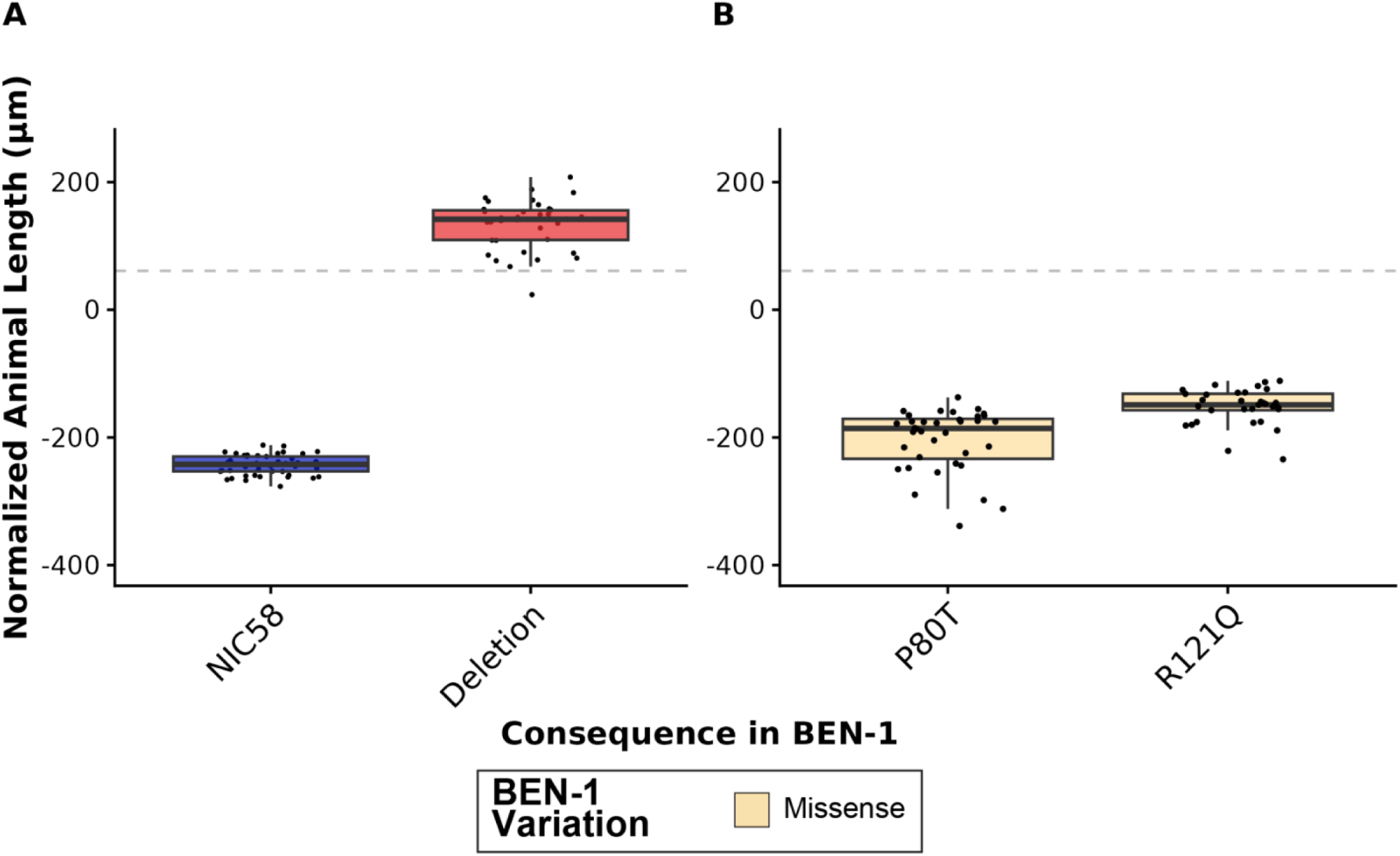
High-throughput larval development assays in the presence of albendazole for *C. tropicalis* strains with unique high-impact variants in BEN-1. The regressed median animal length values for populations of nematodes grown in 30 μM albendazole (ABZ) are shown on the y-axis. Each point represents the normalized median animal length value of a well containing approximately 5-30 animals. Strains are sorted by their relative resistance to ABZ based on median animal length. Data are shown as Tukey box plots with the median as a solid horizontal line, and the top and bottom of the box representing the 75th and 25th quartiles, respectively. The top whisker is extended to the maximum point that is within the 1.5 interquartile range from the 75th quartile. The bottom whisker is extended to the minimum point that is within the 1.5 interquartile range from the 25th quartile. The gray dashed line marks the *C. tropicalis* resistance threshold, defined as two standard deviations below the mean of the *ben-1* deletion strain in the NIC58 reference strain background. Results for **(A)** the NIC58 reference strain (blue) and a strain with a *ben-1* deletion in the NIC58 background (red), and **(B)** all wild *C. tropicalis* strains with unique high-impact variants in *ben-1* are sorted by their relative resistance to ABZ based on median animal length. Wild *C. tropicalis* strains are colored by beta-tubulin variant status.

### The predicted molecular consequence and severity of *ben-1* variants do not predict BZ resistance

To evaluate whether the predicted molecular consequence and severity of *ben-1* variants were predictive of BZ resistance, we characterized the identities and predicted functional impacts of each amino acid altering variant. We distinguished between alleles that disrupt multiple amino acids (*i.e.*, frameshifts, stop/start altering variants, and SVs) and alleles that affect a single amino acid (*i.e.*, missense alleles). Multi-site alleles were excluded from further analysis because they are predicted to severely disrupt or terminate BEN-1 function, whereas missense alleles might preserve an intact protein and allow interpretation of residue-level effects on BZ interaction.

In *C. elegans*, seven unique missense substitutions in BEN-1 were identified, of which five (Y50C, Q131L, S145F, A185P, M275I) were associated with BZ resistance (**S17A Fig.**) [14]. In *C. briggsae*, seven unique missense substitutions in *Cbr-*BEN-1 were identified, but only Q134H was associated with BZ resistance (**S17B Fig.**). In *C. tropicalis*, both missense substitutions (P80T and R121Q) were not associated with BZ resistance (**S17C Fig.**). To evaluate if missense variants with larger predicted functional impacts were associated with BZ resistance, we quantified the severity of each substitution using BLOSUM62 and Grantham scores [39,40]. BLOSUM scores measure the evolutionary likelihood of observing a specific missense substitution, and Grantham scores measure the physicochemical severity of a substitution. Because *C. elegans* had the most missense variants associated with BZ resistance, we performed a regression analysis of each strain’s response to ABZ by the BLOSUM or Grantham scores of the strain’s beta-tubulin variant. For *C. elegans*, neither BLOSUM or Grantham scores showed a significant correlation with ABZ response (**S18 Fig.**). By contrast, too few missense variants were associated with BZ resistance among *C. briggsae* and *C. tropicalis* strains to meaningfully interpret BLOSUM or Grantham scores for these species (**S8 Table**). The limited number of missense substitutions among wild strains and the even smaller subset associated with BZ resistance constrained statistical power to evaluate structure-function relationships between naturally occurring beta-tubulin variants and BZ response.

Next, we mapped missense variants onto an AlphaFold-predicted BEN-1 protein structure that showed that the five *C. elegans* missense substitutions associated with resistance are located within the core, whereas the two substitutions not associated with resistance (P80S and D404N) are on the protein surface (**S17A Fig.**). By contrast, all *C. briggsae* (except Q134H) and all *C. tropicalis* missense substitutions are on the protein surface and are not associated with resistance (**S17B Fig.**). For structural comparison, we modeled *H. contortus tbb-isotype-1* beta-tubulin bound to ABZ, and highlighted the canonical missense variants at positions 167, 198, and 200 (**S17D Fig.**), all of which are located within the core of the structure, near the predicted BZ-binding pocket and are associated with resistance (**S17D Fig.**) [18–22,41]. These results suggest that BZ resistance is associated with missense substitutions occurring within the protein core likely affecting the theoretical BZ-binding pocket.

### Natural variants in *tbb-1*, *tbb-2*, *mec-7*, and *tbb-4* are not associated with ABZ resistance across the three free-living *Caenorhabditis* species

Although it has been established that *ben-1* is the primary gene involved in ABZ resistance in *C. elegans* [14,27,37], we know less about the role each beta-tubulin gene plays in *C. briggsae* and *C. tropicalis* ABZ resistance. Therefore, we identified variants in the other four conserved beta-tubulin genes across the three *Caenorhabditis* species. We found no variants in *Cel-tbb-1* or *Cel-tbb-2* in any *C. elegans* wild strains (CaeNDR Release ID: *C. elegans* - 20231213) (**Fig. 4**). However, we did identify strains with one missense variant (A9T) in *Cel-mec-7*, a splice donor variant in *Cel-mec-7*, or one missense variant in *Cel-tbb-4* (Q8H) (**S1 Table**). The strain with a splice donor variant in *Cel-mec-7* also carried a high-impact variant (*445S) in *Cel-ben-1*. We also assessed the relationship between the expression of *Cel-tbb-1*, *Cel-tbb-2*, *Cel-mec-7*, or *Cel-tbb-4* [38] and ABZ responses in 180 wild strains (**S19 Fig.**) [14,37]. In *C. briggsae*, we identified strains with missense variants in *Cbr-tbb-1* (T35A, V64I, A275T, and L377I), *Cbr-tbb-2* (E441A), *Cbr-tbb-4* (A271V), and *Cbr-mec-7* (T136I, N165S, and S338C) (CaeNDR Release ID: *C. briggsae* - 20240129) (**S2 Table**). In *C. tropicalis*, we identified no high-impact variants in *Ctr-tbb-1*, one missense variant in *Ctr-tbb-2* (N89S), two missense variants in *Ctr-mec-7* (P80F, D433E), and one missense variant in *Ctr-tbb-4* (A78V) (CaeNDR Release ID: *C. tropicalis* - 20231201) (**S3 Table**). Because expression data from *C. briggsae* and *C. tropicalis* wild strains have yet to be collected, we could not test correlations of beta-tubulin gene expression with resistance. We performed HTLDAs under control (DMSO) (**S20 Fig., S21 Fig., and S22 Fig.**) and ABZ conditions (**S23 Fig., S24 Fig., and S25 Fig.**) on strains in the three *Caenorhabditis* species carrying variants in the conserved beta-tubulin genes *tbb-1*, *tbb-2*, *mec-7*, and *tbb-4*, along with strains genetically related to strains with variants in those beta-tubulin genes. No variants outside of *ben-1* conferred ABZ resistance in any species, indicating that natural missense variants in *tbb-1*, *tbb-2*, *mec-7*, and *tbb-4* are not associated with BZ resistance in the three selfing *Caenorhabditis* species.

**Fig 4.**
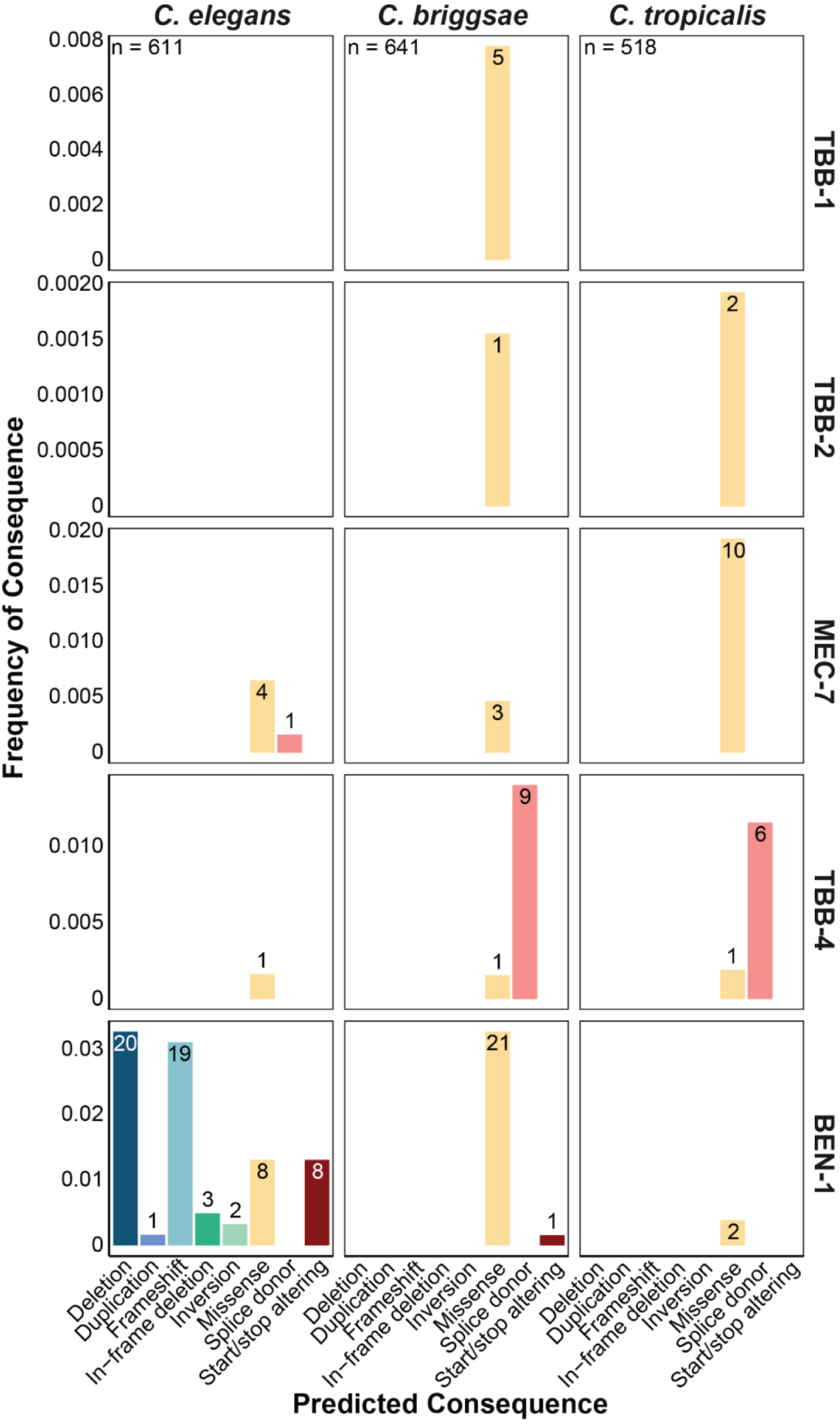
The frequency of predicted high-impact consequences in the five conserved beta-tubulin genes present in natural populations of *C. elegans*, *C. briggsae*, and *C. tropicalis*. The frequency of single nucleotide variants (SNVs) and structural variants (SVs) present in natural populations of *C. elegans* (n = 611), *C. briggsae* (n = 641), and *C. tropicalis* (n = 518) (y-axis) are shown by their predicted consequence in each beta-tubulin gene (x-axis). The total number of isotype reference strains with a given predicted consequence are displayed on top of each bar plot.

### Differences in tissue- and cell-specific expression patterns of beta-tubulins might influence BZ susceptibility

Tissue-specific differences in beta-tubulin expression among the three *Caenorhabditis* species could influence BZ resistance. To compare beta-tubulin expression patterns, we analyzed two whole-animal single-cell transcriptomic datasets that quantify gene expression patterns across *C. elegans*, *C. briggsae*, and *C. tropicalis* [42,43]. A comparison of embryonic gene expression levels in *C. elegans* and *C. briggsae* revealed that beta-tubulin gene expression during embryogenesis is highly conserved (**S9 Table and S10 Table**). On average, the five beta-tubulin gene distances, which quantify expression conservation across homologous cell-types, were small (mean beta-tubulin JSDgene = 0.29), which indicated conserved embryonic expression patterns between *C. elegans* and *C. briggsae* (JSDgene < 0.45) (**S9 Table**) [43]. Expression breadth across cell-types (*i.e.*, cell-type specificity) was also conserved. The broad expression of *tbb-1* and *tbb-2*, and the cell-specific expression of *ben-1*, *tbb-4*, and *mec-7* are conserved between *C. elegans* and *C. briggsae* during embryogenesis (**S10 Table**). Among beta-tubulin genes, *tbb-2* displayed the greatest divergence in embryonic gene expression breadth, the largest difference across species in cell-type specificity (**S10 Table**). Overall, embryonic beta-tubulin expression patterns are conserved between *C. elegans* and *C. briggsae*, although *tbb-2* expression breadth is an exception (**S10 Table**). Next, because neuronal *ben-1* expression restores BZ susceptibility in *C. elegans* [33]we focused on the divergence of beta-tubulin expression across homologous neuronal cell classes at the L2 larval stage [43]. For each gene, we quantified neuron-class-specific expression divergence across the three *Caenorhabditis* species using the proportion of neuronal classes where gene expression differs among species (**S11 Table**) [43]. At least one species expressed *ben-1* in 98 of 118 neuronal cell classes. However, all three *Caenorhabditis* species expressed *ben-1* in only 35 of these classes (Jaccard distance = 0.64) (**S11 Table**). By contrast, *tbb-1* and *tbb-2* had highly conserved expression across the three *Caenorhabditis* species (98 of 118 neuronal cell classes expressed *tbb-1* or *tbb-2* in all three species) (Jaccard distance = 0) (**S11 Table**) [43]. The broad and conserved neuronal expression of *tbb-1* and *tbb-2* contrasts with *ben-1*, which showed narrower and species-specific expression patterns. Next, because *ben-1* expression in cholinergic neurons restores BZ susceptibility in *C. elegans [33]*, we compared cholinergic neuronal expression and identified several cell classes with species-specific patterns. Generally, beta-tubulin cell-specificity is conserved across the three *Caenorhabditis* species (**S9 Table**, **S10 Table**, **S11 Table**). However, examples of expression divergence among *C. elegans*, *C. briggsae*, and *C. tropicalis* were identified for *ben-1* and *tbb-2.* The divergence in *ben-1* expression across cholinergic cell classes indicates that the neuronal sites where *ben-1* function causes BZ susceptibility could differ among species. Future work should test this hypothesis using transgenic strains that express *ben-1* in different tissues and neuronal cells.

### The most diverse high-impact variants are found in *Cel-ben-1*

To better understand the evolution of predicted BZ resistance alleles in the three *Caenorhabditis* species, we assessed the population-wide frequencies of each beta-tubulin variant. First, to determine the prevalence of high-impact variants in the five conserved beta-tubulin genes (*tbb-1*, *tbb-2*, *mec-7*, *tbb-4*, and *ben-1*) across *Caenorhabditis* species, we quantified the frequency of each variant (deletion, duplication, frameshift, in-frame deletion, inversion, missense, splice donor, and start/stop altering) in each species. With global sampling of the three *Caenorhabditis* species (*C. elegans*: 611 strains, *C. briggsae*: 641 strains, *C. tropicalis*: 518 strains), we found that *C. elegans* variants predicted to cause deleterious functional effects were present in 1% of strains. By contrast, variants predicted to cause deleterious functional effects in *C. briggsae* and *C. tropicalis* were rare (< 0.05% of all strains in either species) (**Fig. 4**). The most variation in beta-tubulin genes was identified in *Cel-ben-1*, where deletions, frameshifts, in-frame deletions, inversions, missense, stop/start altering variants, and a duplication were found. Next, we found one missense variant in *Cel*-*tbb-4* and several missense variants and a splice donor in *Cel*-*mec-7*. Overall, *C. elegans* has acquired the most diverse set of high-impact variants in and predicted functional effects on *ben-1*. In *C. briggsae*, we found 21 missense amino acid substitutions and a single start/stop altering consequence in *Cbr-ben-1*. In both *Cbr-tbb-1* and *Cbr-tbb-2*, we identified rare missense consequences. Additionally, we found nine strains with splice variants in *Cbr-tbb-4* and one strain with a missense variant. In *C. tropicalis*, we found missense consequences in all beta-tubulin genes, except *tbb-1*. Additionally, we found six strains with splice variants in *Ctr-tbb-4*. These findings highlight that *Cel-ben-1* has the most diverse set of variants, reinforcing its role in BZ resistance.

Next, we examined the geographic distribution of strains carrying high-impact variants in *tbb-1*, *tbb-2*, *mec-7*, *tbb-4*, and *ben-1* to determine if beta-tubulin variants were associated with natural sampling location. In *C. elegans*, variants in *ben-1* were found globally with no discernible geographic pattern but were concentrated in clades that have experienced recent selective sweeps [44,45] (**Fig. 5**). *Cel*-*ben-1* variants in swept clades suggest that these mutations arose as relatively recent evolutionary events in response to BZ-like compounds in the natural niche. By contrast, in *C. briggsae*, variants in *ben-1* were distributed throughout the species tree and found on more ancestral branches (*i.e.*, earlier diverged lineages in the species) (**Fig. 5C**). For *C. tropicalis*, the limited number of variants in *ben-1* precludes any definitive conclusions regarding their evolutionary patterns (**Fig. 5D**). Because few variants are found in *tbb-1* (**S26 Fig.**), *tbb-2* (**S27 Fig.**), *mec-7* (**S28 Fig.**), and *tbb-4* (**S29 Fig.**), we cannot identify the evolutionary patterns of BZ resistance in these genes for any of the three species.

**Fig 5.**
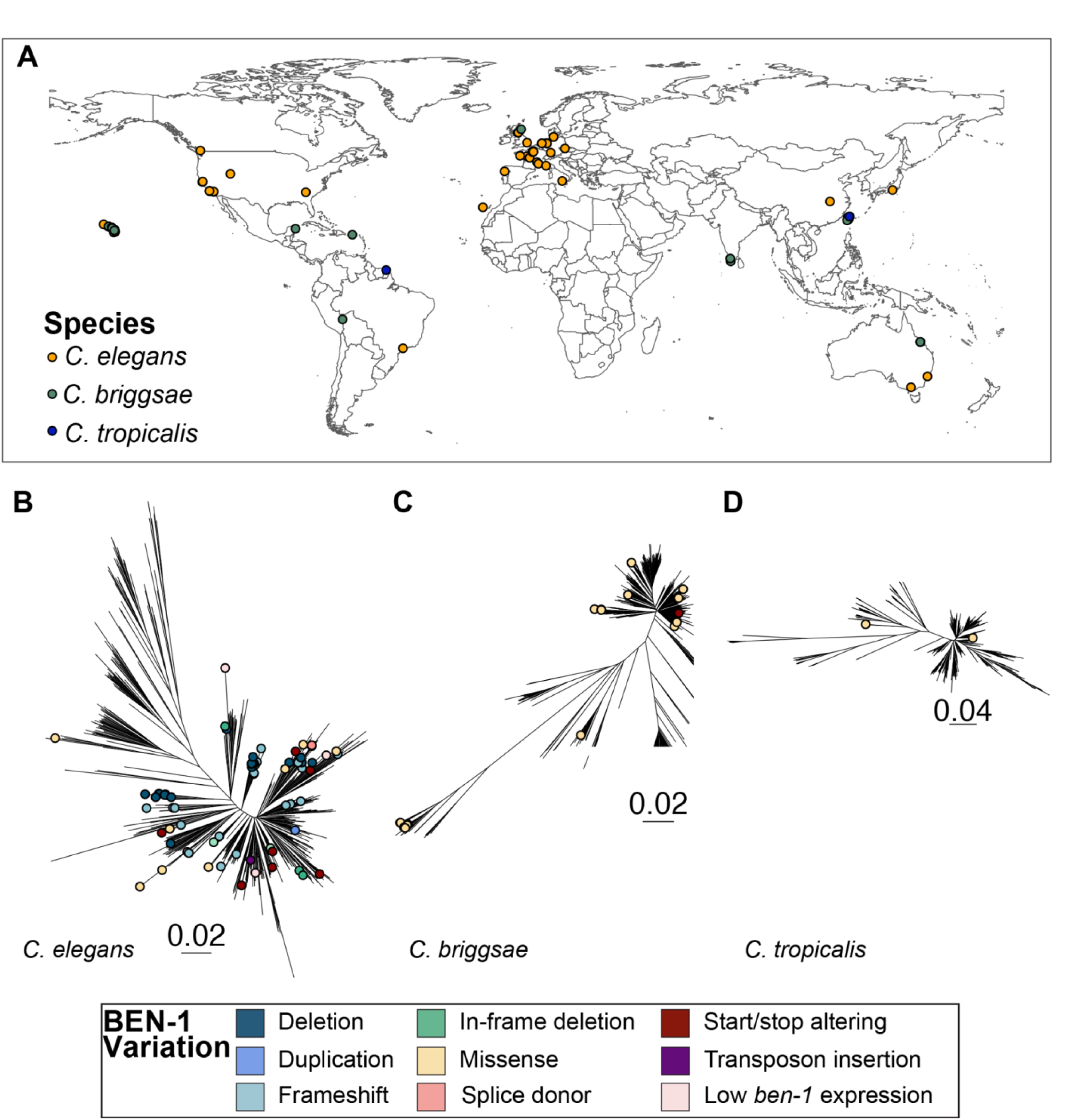
The global distribution of *Caenorhabditis* strains that contain predicted high-impact variation in BEN-1. **(A)** Each point corresponds to the sampling location of an individual *C. elegans* (orange), *C. briggsae* (green), or *C. tropicalis* (blue) isotype reference strain with a predicted high-impact consequence in BEN-1. A genome-wide phylogeny of **(B)** 611 *C. elegans*, **(C)** 641 *C. briggsae*, and **(D)** 518 *C. tropicalis* isotype reference strains, where each point denotes an isotype reference strain with a predicted high-impact consequence in BEN-1 is shown.

Finally, because substrates harbor distinct microbial communities that can influence the evolution of BZ resistance alleles, we determined if strains carrying a high-impact variant in a beta-tubulin gene were associated with specific substrates. We performed substrate enrichment analysis to assess correlations between 12 substrates and all strains in the three *Caenorhabditis* species (**S12 Table**). However, no significant enrichment was observed between a high-impact variant in a beta-tubulin gene and any given substrate (Fisher’s Exact Test, *p*=1) (**S30 Fig.**). Because no geographic or substrate enrichment was observed, evolutionary pressures driving beta-tubulin variation are likely not strongly tied to substrate.

## DISCUSSION

### Beta-tubulin-mediated BZ resistance varies across natural populations of *Caenorhabditis* nematodes

This study provides new insights into beta-tubulin-mediated ABZ resistance across three *Caenorhabditis* species. Each *Caenorhabditis* species harbored a unique set of predicted high-impact beta-tubulin alleles, but only variants in *ben-1* conferred resistance. With additional wild strains since our first study, we identified additional *Cel-ben-1* LoF alleles associated with BZ resistance, which confirmed that a diverse collection of predicted LoF variants in *ben-1* are associated with ABZ resistance in *C. elegans* [14,16]. In *C. briggsae*, strains harboring only two of the eight unique *Cbr-ben-1* variants were resistant to ABZ. One *Cbr-ben-1* variant (W21stop) causes early protein termination, and the other (Q134H) alters a residue within the *Cbr-*BEN-1 protein that likely affects ABZ binding and is associated with resistance in *A. caninum [11]*. A CRISPR-Cas9-generated deletion of *Cbr-ben-1* conferred resistance similar to that displayed by wild strains with these alleles and did not impact fitness. To date, resistance has yet to be identified in *C. tropicalis* wild strains. However, a CRISPR-Cas9-generated deletion of *Ctr-ben-1* conferred resistance and caused a significant reduction in fitness (fecundity), which likely explains why just two high-impact variants were identified among wild strains and neither caused loss of *ben-1* function. Overall, our results highlight the complexity of BZ resistance. Accurate prediction of BZ resistance across nematode species requires a clear understanding of the contributions of both beta-tubulin dependent and beta-tubulin independent mechanisms.

### Nematode survival under BZ exposure depends on beta-tubulin dosage and drug-binding ability

BZs bind to beta-tubulins and inhibit the polymerization of microtubules [2–4]. Therefore, despite the presence of beta-tubulin independent resistance mechanisms [46,47], beta-tubulins have a large impact on BZ efficacy as an anthelmintic treatment. Two factors determine how beta-tubulin impacts susceptibility to BZs: (1) the beta-tubulin’s ability to bind to BZs and (2) the dosage of the beta-tubulin protein, which can be modified by tissue-specific expression. Nematode species have distinct beta-tubulin gene complements with divergence in number of genes, BZ binding affinities, expression levels and cell types, and redundancy. Ultimately, this divergence in beta-tubulin complement shapes the type(s) of resistance-associated alleles that arise in each nematode species. In species where the primary BZ-binding beta-tubulin is redundant with other isoforms, as in selfing *Caenorhabditis* species, loss of a single copy can confer resistance without disrupting essential microtubule functions. In species without redundancy, such as *H. contortus*, beta-tubulin variants that alter the BZ binding affinity are the only available beta-tubulin dependent route to BZ resistance.

In *C. elegans*, *ben-1* is the primary target of BZs but is redundant with *tbb-1* and *tbb-2* [28–30]. LoF alleles in *ben-1* reduce the total amount of BZ-binding protein while preserving essential microtubule functions, which permits nematode development during BZ exposure. A similar pattern likely occurs in *C. briggsae*, where rare high-impact variants in *ben-1* confer resistance, and other beta-tubulin genes maintain essential microtubule functions. By contrast, *H. contortus* relies on the essential beta-tubulin *tbb-isotype-1* and cannot tolerate loss without severe fitness consequences [18]. The differences between the *Caenorhabditis* species and *H. contortus* demonstrate that the prediction of beta-tubulin dependent resistance requires the identification of the total number and expression levels of beta-tubulin genes and determination of which encoded beta-tubulins can bind to BZ. Altogether, BZ resistance depends not on the copy number of beta-tubulins alone but on the availability of functional beta-tubulins capable of interacting with BZs.

### How can we accurately predict BZ resistance across nematode species?

To accurately predict BZ resistance across nematode species, we need (1) tractable nematode models with beta-tubulin gene complements that resemble that of parasite species, (2) improved parasitic nematode genomes that enable the comprehensive identification of beta-tubulin genes, (3) functional tests that define how parasite beta-tubulin alleles contribute to BZ response, and (4) to define the contribution of non-beta-tubulin genes to overall BZ response. To date, no experimentally tractable model nematode species exist with beta-tubulin genes that have both limited redundancy and BZ binding properties similar to those of parasitic nematodes. *Pristionchus pacificus*, a free-living clade V nematode has three beta-tubulin genes, comprising two *ben-1* orthologs (*Ppa-ben-1.1* and *Ppa-ben-1.2*) and an ortholog of *Cel-mec-7* (*Ppa-mec-7*) [48]. All three beta-tubulin genes in *Pristionchus pacificus* are predicted to have high ABZ binding affinity (**S31 Fig.**). Using CRISPR-Cas9 genome editing, we found that homozygous LoF alleles for either *Ppa-ben-1.1* or *Ppa-ben-1.2* could not be recovered in *P. pacificus*, suggesting that both genes are essential **(S13 Table**). The fact that *P. pacificus* has fewer beta-tubulin genes than *C. elegans* likely contributes to the lack of redundancy among beta-tubulins. The number, essentiality, and BZ binding affinity of the *P. pacificus* beta-tubulins more closely resembles that of *H. contortus*, which positions *P. pacificus* as a valuable free-living model to study essential beta-tubulin function, beta-tubulin dosage effects, and BZ resistance relevant to parasitic nematode species.

Second, to understand the role beta-tubulins play in BZ resistance across parasitic nematode species, we must significantly improve genomes and gene models. Most parasitic nematode genomes remain incomplete or are poorly annotated, which obscures beta-tubulin copy number and gene identity. Recent efforts have produced higher quality genomes for some parasitic nematodes [49–51], but we must still define beta-tubulin copy number across diverse nematode species. For example, improved reference genomes and gene models for *C. briggsae* [52,53] and *C. tropicalis* [54] enabled accurate beta-tubulin gene identification for these species. Until genome assemblies, technologies, and analytical techniques improve, we will be unable to accurately predict the number of beta-tubulin genes in a given species and their respective BZ binding affinities.

Third, functional validation of parasitic resistance alleles presents additional challenges. Although tools such as RNAi can work in parasitic nematodes [55], delivery challenges and difficulty isolating edited individuals limit its use. An alternative strategy is to introduce predicted parasite resistance alleles into *C. elegans* to test effects on fitness and BZ response, as shown previously [14–16]. Finally, some parasitic nematodes, such as ascarid species, exhibit BZ resistance without known beta-tubulin resistance alleles [56–58]. This pattern suggests that resistance arises independently of detected beta-tubulin sequence changes or reflects incomplete identification of beta-tubulin genes caused by poor genome assemblies. To date, the underlying cause of BZ resistance in ascarid species remains unclear. Reduced functional beta-tubulin dosage, altered BZ binding, or both could contribute to BZ resistance in ascarids [59]. Improved genome assemblies and annotation are required to identify all beta-tubulin genes and drug-binding sites to define the mechanisms of BZ resistance in ascarid species. In the future, the introduction of newly identified ascarid alleles into free-living nematode models can directly test resistance and fitness, and clarify the mechanisms that drive BZ resistance in clade III nematodes. Additionally, we must define how redundancy, essentiality, beta-tubulin dosage, and drug-binding properties shape the evolution of resistance to improve the prediction of BZ resistance across the huge diversity of parasitic nematode species.

## MATERIALS AND METHODS

### Identification of beta-tubulin loci

Amino acid sequences for all six *C. elegans* beta-tubulin proteins were obtained from WormBase (WS283) [60] and used as queries in a BLASTp search (Version 2.12.0) [61] against protein sequence databases constructed using gene models for *C. briggsae* [53] and *C. tropicalis* [62]. To construct the protein sequence database, we extracted gene model transcript features from the gene feature file with *gffread* (Version 0.9.11) [63] and processed them using the *makeblastdb* function from BLAST (Version 2.12.0). From the BLASTp search, we identified *C. briggsae* and *C. tropicalis* protein sequences with the highest percent identity (PID) to each *C. elegans* beta-tubulin protein. Only protein sequences with the highest PID in both searches were considered orthologs (**S14 Table**). For some *C. elegans* beta-tubulin orthologs, multiple *C. briggsae* or *C. tropicalis* gene models contained multiple splice isoforms. All gene models for all beta-tubulin transcripts were manually inspected, and isoforms that were not fully supported by short-read RNA sequencing data were removed.

### Single nucleotide variant (SNV) and indel calling and annotation

To identify single nucleotide variants (SNVs) or indels (insertions and deletions) in the beta-tubulin genes across the selfing *Caenorhabditis* species, we used the Variant Annotation Tool from the *Caenorhabditis* Natural Diversity Resource (CaeNDR) (Release IDs: *C. elegans* - 20231213, *C. briggsae* - 20240129, *C. tropicalis* - 20231201) [36]. The identified SNVs and indels included small insertions and deletions, frameshifts, altered stop and start codons, nonsynonymous changes, and splice variants (**S1 Table, S2 Table, and S3 Table**).

### Structural variant (SV) calling and annotation

Structural variant (SV) calling was performed using *DELLY* (Version 0.8.3), a SV caller optimized to detect large insertions, deletions, and other complex structural variants such as inversions, translocations, and duplications in paired-end short-read alignments [64] and shown to perform well on *C. elegans* short-read sequence data [65]. SVs that overlapped with beta-tubulin genes were extracted using *bcftools* (Version 1.10.1) [66]. Insertions, deletions, inversions, and duplications that passed the *DELLY* (Version 0.8.3) default quality threshold (greater than three supporting read pairs with a median MAPQ > 20), filtered to high-quality genotypes (genotype quality > 15), and had at least one alternative allele were retained. For complex variants (inversions and duplications), the identification of at least one split-read pair was required (variants flagged as a precise SV by *DELLY*). To validate SVs that passed quality filtering, each SV was manually inspected for breakpoints in the raw-read alignments (*Wally*, Version 0.5.8) and for impacts on the beta-tubulin coding sequence (CaeNDR Genome Browser) [36] (**S15 Table**). We retained SVs where raw read alignments suggested that the SV impacted the beta-tubulin coding sequence. We compared the *Cel-ben-1* SVs called by *DELLY* to those SVs identified previously [14]. *DELLY* successfully recalled structural variants in several strains, including deletions in JU751, JU830, JU1395, JU2582, JU2587, JU2593, JU2829, and QX1233, as well as an inversion in MY518. However, *DELLY* did not detect a previously reported transposon insertion in strain JU3125. To assess if other SVs could have been missed by *DELLY*, we manually inspected the read alignments for all strains that had not been previously phenotyped to check if any other SVs were not detected by *DELLY*. We confirmed the presence of novel *Cel-ben-1* SVs in multiple strains, including putative deletions in ECA706 and NIC1832, and a duplication in NIC1107. Additionally, we identified a previously undetected putative deletion in JU4287. We also examined the amino acids at position 200 in TBB-1, TBB-2, MEC-7, TBB-4, and BEN-1 orthologs and found that all MEC-7, TBB-4, and BEN-1 orthologs contained phenylalanine, and that all TBB-1 and TBB-2 orthologs contained tyrosine at position 200 (**S31 Fig.**).

### Association of *Cel-ben-1* expression with ABZ response

Two previous assays measured developmental responses of wild *C. elegans* strains after ABZ exposure [14,67]. For 180 of these wild *C. elegans* strains, whole-animal expression levels (transcripts per million estimates [TPM]) were collected from untreated young-adult animals [38]. We identified strains with low *Cel-ben-1* expression by selecting strains with TPM values more than one standard deviation (SD) below the mean expression level across all 207 wild strains with expression data. Of the 180 wild *C. elegans* strains, 105 strains were measured for both ABZ response and gene expression. A linear model was built using the *lm* function in R to account for assay effects. Subsequently, the residuals of the linear model were used to normalize previous measures of ABZ response. We evaluated the linear fit between each strain’s expression of *ben-1* (**S1 Fig.**), *tbb-1*, *tbb-2*, *mec-7*, or *tbb-4* (**S19 Fig.**) and the developmental delay following ABZ exposure.

### Phylogenetic analysis

We characterized the relatedness of isotype reference strains (genetically unique strains) with beta-tubulin variants using species trees downloaded from CaeNDR and generated by the ‘post-gatk-nf’ pipeline (https://github.com/AndersenLab/post-gatk-nf) (Release IDs: *C. elegans* - 20231213, *C. briggsae* - 20240129, *C. tropicalis* - 20231201) [36]. Briefly, the trees were generated using high-quality SNVs in isotype reference strains retained in the hard-filtered variant call format (VCF) file. *vcf2phylip* [68] and the *bioconvert* [69] function *phylip2stockholm* were used to prepare inputs for *quicktree*, which was used to construct a tree using a neighbor-joining algorithm [70]. All versions of these software can be accessed from the ‘post-gatk’ docker container (https://hub.docker.com/r/andersenlab/tree), used by the ‘post-gatk-nf’ pipeline. We visualized the trees for each species using the *ggtree* function from the *ggtree* (v3.6.2) R package [71].

### Strain selection and maintenance

Eighteen *C. elegans* strains, 45 *C. briggsae* strains, and 15 *C. tropicalis* strains from the CaeNDR [36] were used in this study (**S1 Table, S2 Table, and S3 Table)**. Isolation details for each strain are included in CaeNDR. For each species, we selected strains that had variants (SNV or SV) with unique high-impact consequences in *tbb-1*, *tbb-2*, *mec-7*, *tbb-4*, or *ben-1* that had not been previously phenotyped. High-impact consequences included changes to amino acids, start and stop codon positions, or splice variants [36]. Strains with high-impact consequences in a beta-tubulin gene are herein referred to as “predicted resistant” strains. Strains that were closely related to predicted resistant strains with no high-impact consequences in beta-tubulin genes were also included for *C. briggsae* and *C. tropicalis* and herein classified as “predicted susceptible” strains (**S2 Fig., S3 Fig., and S4 Table**). The reference strains for all three species were included.

Before measuring ABZ responses, *C. elegans* and *C. briggsae* animals were maintained at 20°C and *C. tropicalis* animals were maintained at 25°C. All animals were maintained on 6 cm plates with modified nematode growth medium (NGMA), which contains 1% agar and 0.7% agarose to prevent animals from burrowing [72]. The NGMA plates were seeded with the *Escherichia coli* strain OP50 as a nematode food source. All strains were grown for three generations without starvation on NGMA plates before anthelmintic exposure to reduce the transgenerational effects of starvation stress [73].

### CRISPR-Cas9 genome editing

To validate the role that *ben-1* plays in ABZ resistance in the three *Caenorhabditis* species, we used CRISPR-Cas9 to create *ben-1* deletions. For *C. elegans*, we used a previously generated strain (ECA882) with a *ben-1* deletion in the N2 background [14,15,27,37]. The *ben-1* deletions were generated in the AF16 background for *C. briggsae* and the NIC58 background for *C. tropicalis*. Injections were performed by InVivo Biosystems (Eugene, OR), and deletions of *Cbr-ben-1* and *Ctr-ben-1* were confirmed using PCR (**S10 Fig. and S11 Fig.**). Briefly, two primer pairs were designed for the deletion alleles for each species, with each pair designed to bind to a region external or internal to both of the deletions. Confirmation of deletion was performed by performing two amplification reactions for each sample: (1) the use of both external primers, and (2) the use of an internal and external pair (**S5 Table**). The parental strain was used as a control in each PCR. Deletions were confirmed by a reduction in the size of the external-external amplicon in the edited strains compared to the unedited parental control strain. Homozygosity was confirmed by the loss of a band amplified from the external-internal primer pair. Edited strains underwent two generations of PCR confirmation for homozygosity. Two independent edits of each allele in each species were generated to control for any potential off-target effects caused by CRISPR-Cas9 genome editing (**S12 Fig., S13 Fig., and S5 Table**).

### Nematode food preparation for NGMA 6 cm plates

The OP50 *E. coli* strain was used as a nematode food source for NGMA plates. A frozen stock of OP50 was streaked onto a 10 cm Luria-Bertani (LB) agar plate and incubated overnight at 37°C. The following day, a single bacterial colony was transferred into each of two culture tubes that contained 5 mL of 1x LB. The starter cultures and two negative controls (1X LB without *E. coli*) were incubated for 18 hours at 37°C shaking at 210 rpm. The OD_600_ value of the starter cultures were measured using a spectrophotometer (BioRad, SmartSpec Plus) to calculate how much starter culture was needed to inoculate a 1 L culture at an OD_600_ value of 0.005. For each assay, one culture containing 1 L of pre-warmed 1X LB inoculated with the starter culture grew for approximately 4 - 4.5 hours at 37°C at 210 rpm to an OD_600_ value between 0.45 and 0.6. Cultures were transferred to 4°C to slow growth. OP50 was spotted on NGMA test plates (two per culture) and grown at 37°C overnight to assay for contamination.

### Nematode food preparation for high-throughput larval development assays (HTLDAs)

One batch of HB101 *E. coli* was used as a nematode food source for all HTLDAs in this study. A frozen stock of HB101 *E. coli* was streaked onto a 10 cm LB agar plate and incubated overnight at 37°C. The following day, a single bacterial colony was transferred into three culture tubes that contained 5 mL of 1x Horvitz Super Broth (HSB). The starter cultures and two negative controls (1X HSB without *E. coli*) were incubated for 18 hours at 37°C shaking at 180 rpm. The OD_600_ value of the starter cultures were measured using a spectrophotometer (BioRad, SmartSpec Plus) to calculate how much starter culture was needed to inoculate a 1 L culture at an OD_600_ value of 0.001. A total of four cultures each containing 1 L of pre-warmed 1X HSB inoculated with the starter culture grew for 15 hours at 37°C while shaking at 180 rpm. After 15 hours, flasks were removed from the incubator and transferred to 4°C to slow growth. The 1X HSB was removed from the cultures by performing three rounds of centrifugation, where the supernatant was removed, and the bacterial cells were pelleted. Bacterial cells were washed with K medium, resuspended in K medium, pooled, and transferred to a 2 L glass beaker. The OD_600_ value of the bacterial suspension was measured and diluted to a final concentration of OD_600_100 with K medium, aliquoted to 15 mL conicals, and stored at -80°C for use in the HTLDAs.

### ABZ dose-response assays for *C. briggsae* and *C. tropicalis*

Because ABZ response had been minimally characterized in *C. briggsae* [74] and has not yet been described in *C. tropicalis*, we first measured dose-response curves for both species after exposure to ABZ to assess developmental delay. Before performing HTLDAs, ABZ (Sigma-Aldrich, Catalog # A4673-10G) stock solutions were prepared in dimethyl sulfoxide (DMSO) (Fisher Scientific, Catalog # D1281), aliquoted, and stored at -20°C for use in the assays. For the dose-response assays, animals were exposed to ABZ at the following concentrations (μM): 0 (1% DMSO), 0.12, 0.23, 0.47, 0.94, 1.88, 3.75, 7.5, 15, 30, 60, and 120. Animals developed in the presence of ABZ as described in *HTLDAs to assess nematode development*.

Dose-response model estimation and statistics were performed as described previously [67,75]. Briefly, a four-parameter log-logistic dose-response curve was fit independently for a genetically diverse set of 11 *C. briggsae* strains (**S14 Fig.**) and seven *C. tropicalis* strains (**S16 Fig.**), where normalized median animal length was used as a metric for phenotypic response (see *HTLDA data collection and data cleaning*). For each strain-specific dose-response model, slope (*b*) and concentration (*e*) were estimated with strain as a covariate. We calculated EC_10_ as we have previously found EC_10_ response to be more heritable than half maximal effective concentration (EC_50_) estimates and were used in our analysis [67,75]. A dosage of 30 μM ABZ was closest to the EC_10_ for *C. briggsae* and *C. tropicalis*, consistent with ABZ concentrations used in past *C. elegans* assays [15,16,37] and in all HTLDAs in this study.

### HTLDAs to assess nematode development

Populations of each strain were amplified and bleach-synchronized in three independent assays. Independent bleach synchronizations controlled for variation in embryo survival and subsequent effects on developmental rates. After bleach synchronization, approximately 30 embryos were dispensed into each well of a 96-well microplate in 50 μL of K medium. Each strain had sixteen wells per condition (DMSO or ABZ) in each assay. Three independent assays yielded a total of forty-eight wells per condition per strain. Each 96-well microplate was prepared, labeled, and sealed using gas-permeable sealing films (Fisher Scientific, Catalog # 14-222-043). Plates were placed in humidity chambers to incubate for 24 hours at 20°C for *C. elegans* and *C. briggsae*, and 25°C for *C. tropicalis* while shaking at 170 rpm (INFORS HT Multitron shaker). After 24 hours, every plate was inspected to ensure that all embryos hatched and animals were developmentally arrested at the first larval (L1) stage so all strains started each assay at the same developmental stage. Next, food was prepared to feed the developmentally arrested L1 animals using the required number of OD_600_100 HB101 aliquots (see *Nematode food preparation for HTLDAs*). The HB101 aliquots were thawed at room temperature, combined into a single conical tube, and diluted to an OD_600_30 with K medium. To inhibit further bacterial growth and prevent contamination, 150 μL of kanamycin was added to the HB101. An aliquot of 100 μM ABZ stock solution was thawed at room temperature and added to an aliquot of OD_600_30 K medium at a 3% volume/volume ratio. Next, 25 μL of the food and ABZ mixture was transferred into the appropriate wells of the 96-well microplates to feed the arrested L1 animals at a final HB101 concentration of OD_600_10 and expose L1 animals to ABZ. Immediately afterward, the 96-well microplates were sealed using a new gas permeable sealing film, returned to the humidity chambers, and incubated for 48 hours at 20°C (*C. elegans* and *C. briggsae*) or 42 hours at 25°C (*C. tropicalis*) while shaking at 170 rpm. After 48 hours (*C. elegans* and *C. briggsae*) or 42 hours (*C. tropicalis*) of incubation and shaking in the presence of food and either DMSO or ABZ, the 96-well microplates were removed from the incubator and treated with 50 mM sodium azide in M9 for 10 minutes to paralyze and straighten nematodes. After 10 minutes, images of nematodes in the microplates were immediately captured using Molecular Devices ImageXpress Nano microscope (Molecular Devices, San Jose, CA) using a 2X objective. The ImageXpress Nano microscope acquires brightfield images using a 4.7 megapixel CMOS camera and stores images in a 16 - bit TIFF format. The images were used to quantify the development of nematodes in the presence of DMSO or ABZ as described below (see *High-throughputput imager assays [HTLDA] data collection and cleaning*). A full step-by-step protocol for the HTLDA has been deposited on <u>protocols.io</u> [76].

### HTLDA data collection and data cleaning

*CellProfiler* (Version 24.10.1) was used to characterize and quantify biological data from the image-based assays. Custom software packages designed to extract animal measurements from images collected on the Molecular Devices ImageXpress Nano microscope were previously described [77]. *CellProfiler* modules and *WormToolbox* were developed to extract morphological features of individual animals from images from the HTLDA [78]. Worm model estimations and custom *CellProfiler* pipelines were written using the *WormToolbox* in the GUI-based instance of *CellProfiler* [75]. Next, a Nextflow pipeline (Version 24) was written to run command-line instances of *CellProfiler* in parallel on the Rockfish High-Performance Computing Cluster (Johns Hopkins University). The *CellProfiler* workflow can be found at https://github.com/AndersenLab/cellprofiler-nf. The custom *CellProfiler* pipeline generates animal measurements by using four worm models: three worm models tailored to capture animals at the L4 larval stage, in the L2 and L3 larval stages, and the L1 larval stage, as well as a “multi-drug high dose” (MDHD) model, to capture animals with more abnormal body sizes caused by extreme anthelmintic responses. These measurements comprised our raw dataset. Two *C. briggsae* strains (NIC1052 and VX34) were not fully paralyzed and straightened at the time of imaging, which created some misclassification of animal measurements. Thus, the animal lengths for strains NIC1052 and VX34 measured by *CellProfiler* are shorter than the actual animal lengths. However, the difference in animal lengths does not affect the classification of a strain as resistant or sensitive to ABZ. Data cleaning and analysis steps were performed using a custom R package, *easyXpress* (Version 2.0) [77] and followed methods previously reported [37]. All analyses were performed using the R statistical environment (Version 4.2.1) unless stated otherwise.

### C. briggsae and C. tropicalis fecundity assays

To define the fitness costs associated with a loss of *ben-1* in *C. briggsae* and *C. tropicalis*, we performed fecundity assays. For *C. briggsae*, we used the two strains with independent edits of *ben-1* in the AF16 background (ECA3953 and ECA3954) and the AF16 reference strain. For *C. tropicalis*, we used the two strains with independent edits of *ben-1* in the NIC58 background (ECA4247 and ECA4248) and the NIC58 reference strain. To perform fecundity assays, we placed a single L4 larval stage hermaphrodite from each strain onto a 6 cm NGMA plate that was spotted with *E. coli* OP50. *C. briggsae* assay plates were maintained at 20°C, and *C. tropicalis* assay plates were maintained at 25°C. For each assay plate, the original hermaphrodite parent was transferred to a fresh 6 cm NGMA plate every 24 hours for 192 hours. Ten technical replicates were prepared for each strain. The Basic Imaging Platform from Tau Scientific was used to collect images for each of the assay plates (0, 24, 48, 72, 96, and 120-hours) at either 48 or 72 hours for *C. briggsae* and *C. tropicalis* animals, respectively, after the removal of the parent from each NGMA plate. The total offspring was counted from each image by visual inspection using the Multi-point tool in ImageJ (Version 1.54g). The original hermaphrodite parents were excluded from the counts. Replicates where the original hermaphrodite parent died were excluded from the analysis. Only biological replicates with data from at least six assay plates were used to calculate total fecundity (**S15 Fig., S6 Table, and S7 Table**).

### Tissue-specific beta-tubulin gene expression conservation in *C. elegans*, *C. briggsae*, and *C. tropicalis*

To assess beta-tubulin expression divergence across neuronal cell classes, we used whole-animal single-cell transcriptomes of *C. elegans*, *C. briggsae*, and *C. tropicalis* [43]. We quantified neuronal cell expression divergence across species using precomputed Jaccard distances calculated from the neuronal cell classes in at least one species and the number of neuron classes in which a gene is expressed in all three species [43]. We also downloaded gene expression summary data for each homologous cell class from *CaenoGen* and analyzed expression patterns in cell classes that used acetylcholine (ACh) as a neurotransmitter in *C. elegans [79]*. We identified cell classes in which *ben-1* orthologs showed species-specific presence or absence of expression.

We also analyzed embryonic single-cell transcriptomes from the *C. briggsae* and *C. elegans* reference strains to examine the conservation of beta-tubulin gene expression in homologous cell types [42]. We used pre-computed summary statistics to quantify two components of expression divergence among beta-tubulin genes: gene distance (reported as Jensen-Shannon Distances for each gene) and expression breadth (reported as Tau metrics for each species). Beta-tubulin gene distances and expression breadth metrics are available in the gene data summary table file (**S9 Table and S10 Table**). Estimates for three cell populations are distinguished by cell-type assignment: progenitor, terminal, and joint (combined progenitor and terminal cells).

### Protein structure and visualization of *ben-1* and *tbb-isotype-1* variants

For *C. elegans*, the BEN-1 amino acid sequence was obtained from WormBase (WS283) [60]. For *C. briggsae* and *C. tropicalis*, BEN-1 amino acid sequences were the best hits to the *C. elegans* BEN-1 query from the reciprocal BLASTp search used to identify beta-tubulin orthologs (*see Identification of beta-tubulin loci*) (**S16 Table**). The *H. contortus tbb-isotype-1* amino acid sequence was obtained from WormBase Parasite (Version: WBPS19) [41]. Protein structures were predicted using AlphaFold3 [80]. All BEN-1 variant data and associated benzimidazole-response phenotypes were compiled from this study and previously published data [14,37]. A custom Python script was used to generate a PyMOL script (Version 3.1.6.1) for structural visualization. Each predicted beta-tubulin structure was aligned to *Cel-*BEN-1 using the PyMOL ‘*align*’ command.

### Mutation of ben-1 orthologs in Pristionchus pacificus

To test the function of *ben-1* in *Pristionchus pacificus*, we used CRISPR-Cas9 genome editing to delete the two orthologs of *Cel-ben-1*. Designed crRNAs and tracrRNA were synthesized by Integrated DNA Technologies (IDT). For the creation of the guide RNAs, 3 μL of 100 μM tracrRNA (IDT) was combined with 3 μL of 100 μM crRNA (IDT) and incubated at 95°C for five minutes, followed by five minutes at room temperature for annealing. The ribonucleoprotein (RNP) complex was then created by combining 0.61 μL of the guide-RNA mixture with 0.25 μL of Cas9 protein (IDT), followed by incubation at 37°C for 10 minutes. The final microinjection mix was then created by combining the 0.86 μL RNP complex with 9.14 μL of a TE buffer mixture containing a (55 ng/μL) plasmid containing a *P. pacificus* codon-optimized *egl-20p::TurboRFP::rpl-23UTR* construct [81] used as a co-injection marker for visual identification of successful injections by the presence of fluorescent F1 individuals (**S13 Table)**. All F1 individuals displaying fluorescence were then isolated and allowed to self-fertilize. After successful F2 embryo hatching, F1 individuals were genotyped for mutations using a heteroduplex mobility assay (HMA). Individuals from F2 broods that were determined to contain heterozygous mutations in F1 mothers were then isolated, allowed to self-fertilize, and genotyped by HMA after F3 embryos had hatched to identify homozygous mutant lines, followed by Sanger sequencing to determine types of mutations induced. Additionally, for mutations that were homozygous lethal, the F3 from heterozygous F2 individuals were likewise genotyped by HMA to quantify survivability based on deviations from expected Mendelian ratios.

## DATA AVAILABILITY STATEMENT

All code and data used to replicate the data analysis and figures are available on GitHub at: https://github.com/AndersenLab/ce_cb_ct_betatubulin. S1 Table contains the list of *C. elegans* isotype reference strains, any beta-tubulin variants, sample collection locations, and substrate types. S2 Table contains the list of *C. briggsae* isotype reference strains, any beta-tubulin variants, sample collection locations, and substrate types.

S3 Table contains the list of *C. tropicalis* isotype reference strains, any beta-tubulin variants, sample collection locations, and substrate types. S4 Table contains all *C. briggsae* and *C. tropicalis* isotype reference strains with beta-tubulin variants and genetically related strains. S5 Table contains details about the CRISPR-Cas9 genome-edited strains, CRISPR-Cas9 reagents, and oligonucleotide sequences for the deletion of *ben-1* in *C. briggsae* and *C. tropicalis*. S6 Table contains the results from the fecundity assays of the *C. briggsae* reference strain (AF16) and the *Cbr-ben-1* deletion strains.Table S7 contains the results from the fecundity assays of the *C. tropicalis* strain NIC58 and the *Ctr-ben-1* deletion strains. S8 Table contains the BLOSUM62 and Granthum scores for amino acid changes in the beta-tubulin genes in the three *Caenorhabditis* species. S9 Table contains expression breadth (Tau) estimates for beta-tubulin genes calculated on *C. elegans* and *C. briggsae* embryonic cells by Large *et al.* 2024. S10 Table contains conservation estimates of embryonic expression patterns between *C. elegans* and *C. briggsae* with Jensen-Shannon gene distances estimated in Large *et al.* 2024 for beta-tubulin genes. S11 Table contains the neuronal cell-class expression divergence of beta-tubulin orthologs across *C. elegans*, *C. briggsae*, and *C. tropicalis*, reported as Jaccard distances obtained from Toker *et al.* 2025. S12 Table contains results from the substrate enrichment analysis. S13 Table contains details about the CRISPR-Cas9 genome edited strains, CRISPR-Cas9 reagents, and oligonucleotide sequences for the deletion of *ben-1* in *P. pacificus*. S14 Table contains all beta-tubulin transcript IDs in the three *Caenorhabditis* species. S15 Table contains the manual curation of SVs found in beta-tubulin genes. S16 Table contains the amino acid sequences for the BEN-1 proteins from the three *Caenorhabditis* species.

## FUNDING

Amanda O. Shaver was funded by the National Institutes of Health grant F32AI181342. Ryan McKeown was supported by Northwestern University’s Biotechnology Training Program (NIH T32 GM008449). This work was supported by the National Institutes of Health grant R01AI153088 to E.C.A.

## CRediT AUTHOR STATEMENT

**Conceptualization**: ECA

**Data curation:** AOS, RM

**Formal analysis:** AOS, RM

**Funding acquisition:** ECA

**Investigation:** AOS, JMRO, JBC, DWH, JSF, SD, EJR

**Methodology:** AOS, RM

**Project administration:** ECA

**Resources:** ECA

**Software:** AOS, RM

**Supervision:** ECA

**Validation:** AOS, RM

**Visualization:** AOS, RM

**Writing - original draft:** AOS

**Writing - reviewing & editing:** AOS, RM, ECA

## DECLARATION OF COMPETING INTERESTS

The authors have declared that no competing interests exist.

## ACKNOWLEDGEMENTS

We would like to thank members of the Andersen laboratory for their feedback and helpful comments on this manuscript. We thank members of the *C. elegans* community for collecting the diverse set *Caenorhabditis* strains included in this study and the *Caenorhabditis* Natural Diversity Resource (NSF Capacity grant 2224885) for providing the strains for this study. We thank WormBase for providing the amino acid sequences for all six *C. elegans* beta-tubulin proteins [60]. We thank WormBase Parasite for providing the amino acid sequences for TBB-ISOTYPE-1 beta-tubulin protein in *H. contortus* [41].

## Notes

https://github.com/AndersenLab/ce_cb_ct_betatubulin

### Competing Interest Statement

The authors have declared no competing interest.

### Summary of Updates

Much of the manuscript has been substantially revised in response to reviewer feedback.

