## SUPPORTING INFORMATION for "Evaluating beta-tubulin variants as predictors of benzimidazole resistance across *Caenorhabditis* nematodes"

### SUPPORTING TABLES

**S1 Table. *C. elegans* isotype variant table**
**S2 Table. *C. briggsae* isotype variant table**
**S3 Table. *C. tropicalis* isotype variant table**
**S4 Table. *C. briggsae* and *C. tropicalis* strain pairs**
**S5 Table. Table of CRISPR-Cas9 genome edited strains, CRISPR-Cas9 reagents, and oligonucleotide sequences**
**S6 Table. Results from *C. briggsae* fecundity assays**
**S7 Table. Results from *C. tropicalis* fecundity assays**
**S8 Table. BLOSUM and Grantham scores for amino acid changes in beta-tubulin genes in the three *Caenorhabditis* species**
**S9 Table. Conservation of embryonic expression patterns between *C. elegans* and *C. briggsae* with Jensen-Shannon gene distances estimated in *Large et al. 2024* for beta-tubulin genes** Jensen-Shannon gene distances ($JSD_{gene}$) quantify expression conservation across homologous embryonic cell types and range from zero (conserved) to one (diverged), where values below 0.45 indicate conserved expression patterns.
**S10 Table. Expression breadth estimates for beta-tubulin genes calculated on *C. elegans* and *C. briggsae* embryonic cells by Large *et al.* 2024** *Tau* estimates capture the cell-specific expression patterns of each gene for *C. elegans* and *C. briggsae*. Estimates range from zero (broad expression) to one (cell-type specific expression). The absolute difference between *C. elegans* and *C. briggsae Tau* is reported to reflect the divergence in cell-specificity across species for each beta-tubulin ortholog.
**S11 Table. Neuronal cell-class expression divergence of beta-tubulin genes across *C. elegans*, *C. briggsae*, and *C. tropicalis* obtained from *Toker et al.* 2025** Jaccard distances measure the proportion of neuronal classes where gene expression differs among species. Range from zero to one, where higher values (~1) indicate that a greater proportion of cell classes express the gene in only one or two species.
**S12 Table. Location and substrate where each isotype reference strain was collected**
**S13 Table. Genes targeted, CRISPR-Cas9 guide RNA, and detection-primer sequences used, and mutant alleles produced in *Pristionchus pacificus***
**S14 Table. Beta-tubulin transcript IDs**
**S15 Table. Manual curation of SVs**
**S16 Table. Amino acid sequences of the BEN-1 protein for the three Caenorhabditis species**

### SUPPORTING FIGURES

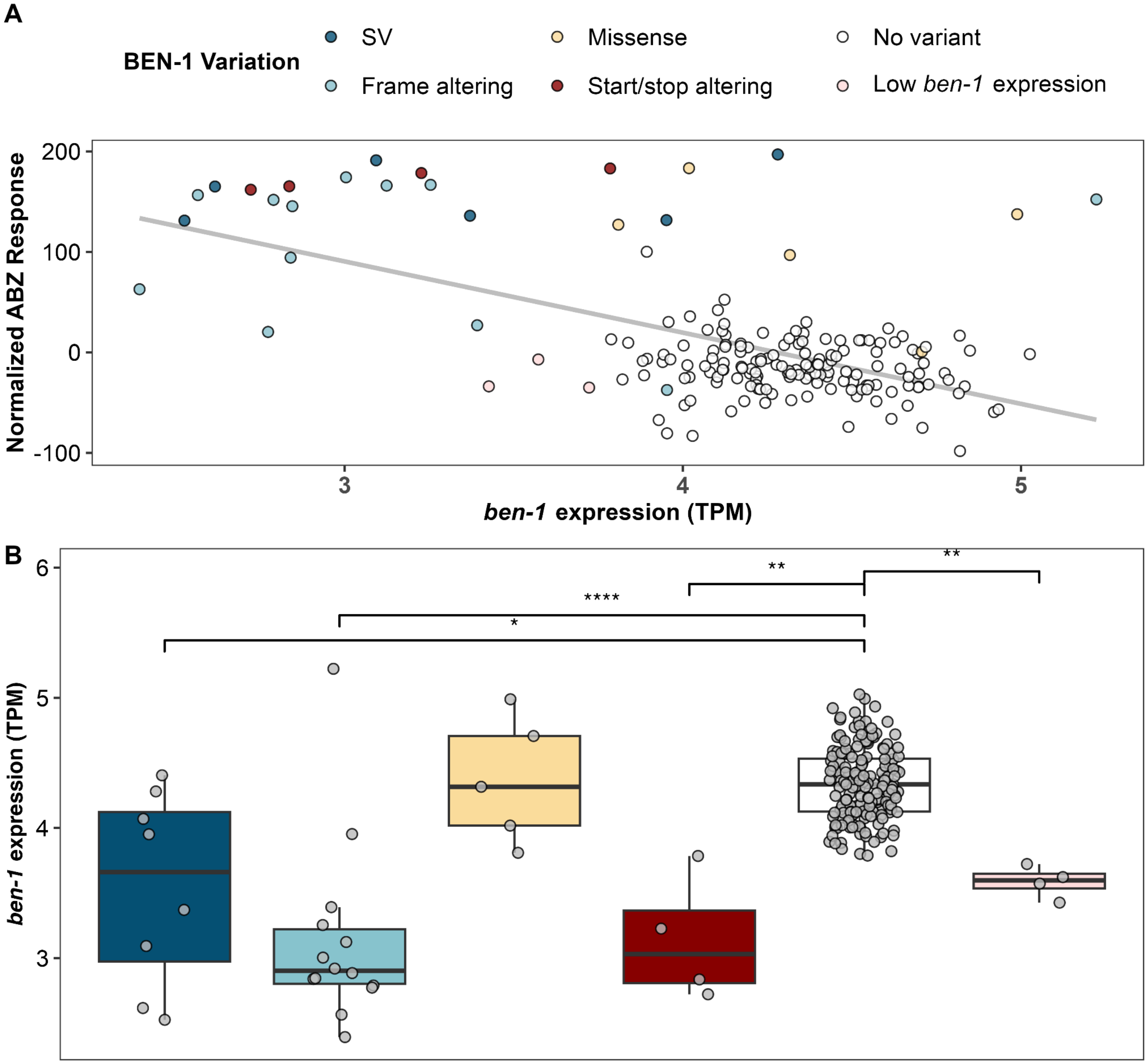


**S1 Fig. The relationship between *ben-1* expression levels and albendazole response in *C. elegans* strains.**

**(A)** Scatterplot of the relationship between *ben-1* expression levels and normalized albendazole (ABZ) response across *C. elegans* wild strains. Each point represents a strain phenotyped for ABZ response in previous publications (Hahnel *et al.*, 2018; Shaver *et al.*, 2024) with *ben-1* expression data (Zhang et al. 2022). The *ben-1* expression level measured in transcripts per million (TPM) is displayed on the x-axis. The normalized ABZ response values adjusted for assay-specific effects are displayed on the y-axis. The gray line represents the linear regression fit between *ben-1* expression and normalized response ($R^2$ = 0.34, *p*-value = 5.16e-18), with the linear model's coefficient of determination ($R^2$). Data points are colored based on the predicted functional consequence of the *ben-1* allele for each strain (*i.e.*, large structural variant (SV), frameshift, missense substitution, disrupted start/stop sequence, no high-impact variant, or low *ben-1* expression). **(B)** Boxplots of *ben-1* expression levels among strains grouped by the predicted functional consequences of their *ben-1* alleles. Each point represents the *ben-1* expression level of an individual within each group. We tested for statistically significant differences in the expression between each consequence type and wild strains without a high-impact *ben-1* allele with an unpaired Wilcoxon test. Significance levels are indicated by symbols: '*' ($p < 0.05$), '**' ($p < 0.01$), '***' ($p < 0.001$), '****' ($p < 0.0001$).

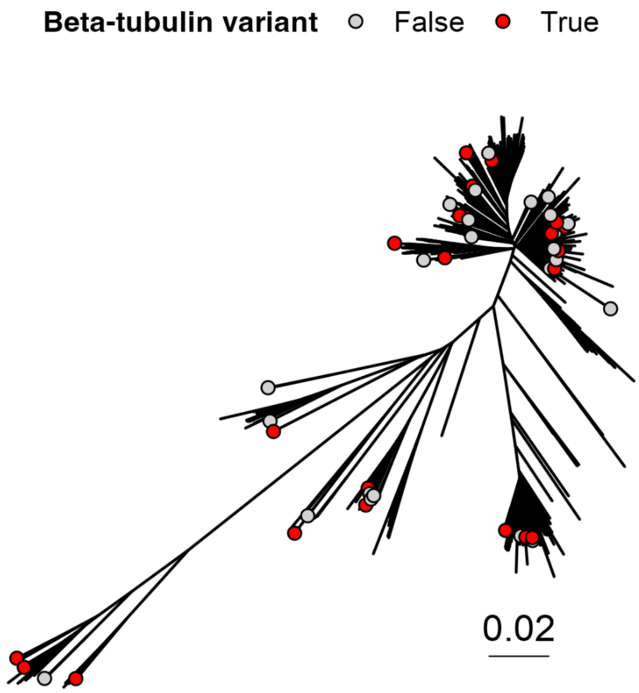


**S2 Fig. *C. briggsae* species tree highlighting isotype reference strains tested for ABZ resistance.**
*C. briggsae* strains included in high-throughput larval development assays (HTLDAs) are highlighted on the *C. briggsae* species tree. Strains with predicted high-impact variants in a beta-tubulin gene are denoted by red points. Strains with no predicted variants in any beta-tubulin gene are denoted by gray points.

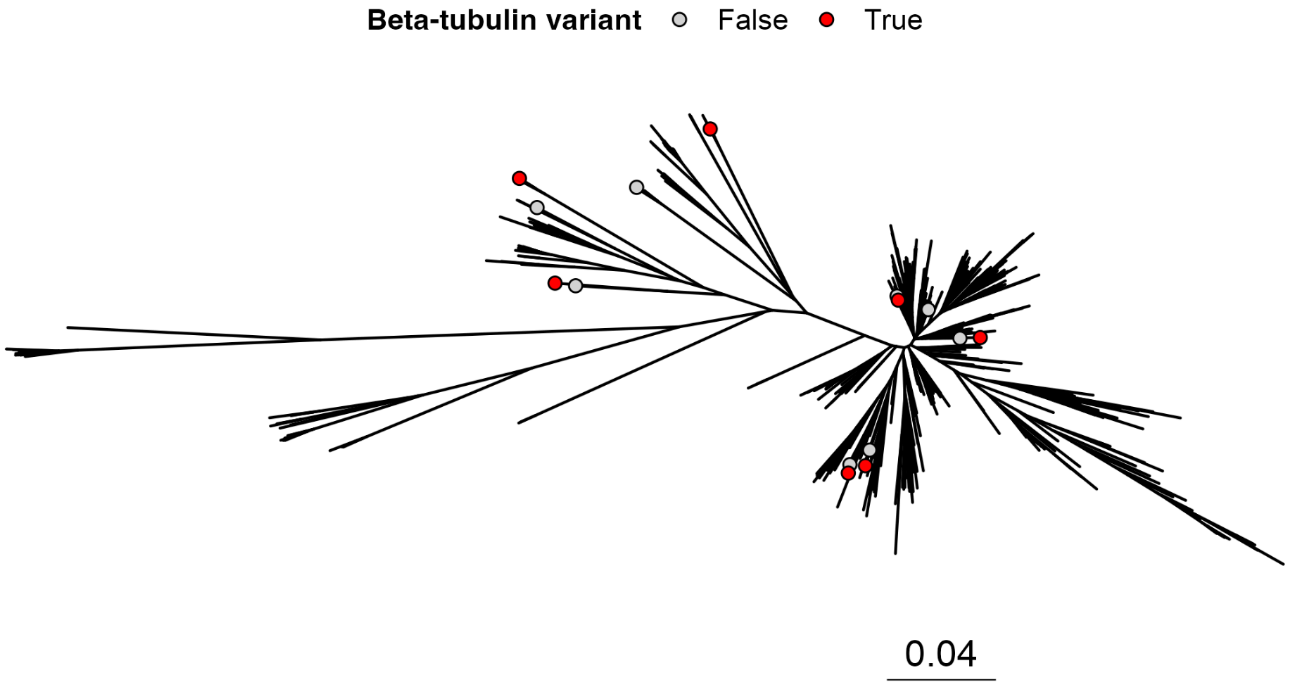


**S3 Fig. *C. tropicalis* species tree highlighting isotype reference strains tested for ABZ resistance.** *C. tropicalis* strains included in high-throughput larval development assays (HTLDAs) are highlighted on the *C. tropicalis* species tree. Strains with predicted high-impact variants in a beta-tubulin gene are denoted by red points. Strains with no predicted variants in any beta-tubulin gene are denoted by gray points.

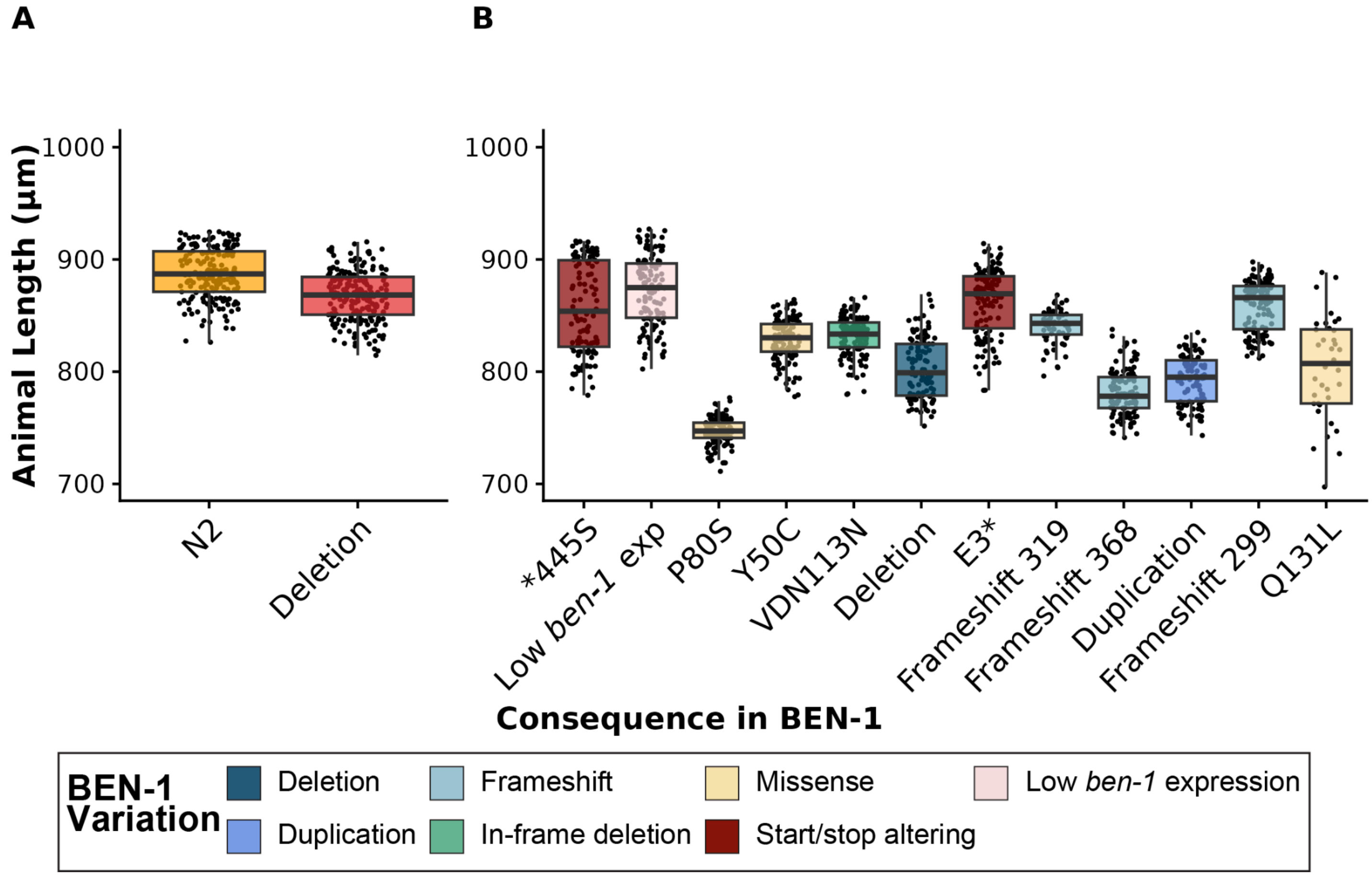


**S4 Fig. HTLDAs for *C. elegans* strains with high-impact BEN-1 variants in control conditions.** Median animal length values from populations of nematodes grown in DMSO are shown on the y-axis. Each point represents the median animal length from a well containing approximately 5-30 animals. Data are shown as Tukey box plots with the median as a solid horizontal line, the top and bottom of the box representing the 75th and 25th quartiles, respectively. The top whisker is extended to the maximum point that is within a 1.5 interquartile range from the 75th quartile. The bottom whisker is extended to the minimum point that is within the 1.5 interquartile range from the 25th quartile. Results for **(A)** the N2 reference strain (orange) and a strain with a *ben-1* deletion in the N2 background (red), and **(B)** all wild *C. elegans* strains with unique high-impact variants in *ben-1* are sorted by their relative resistance to ABZ based on median animal length. Wild *C. elegans* strains are colored by beta-tubulin variant status.

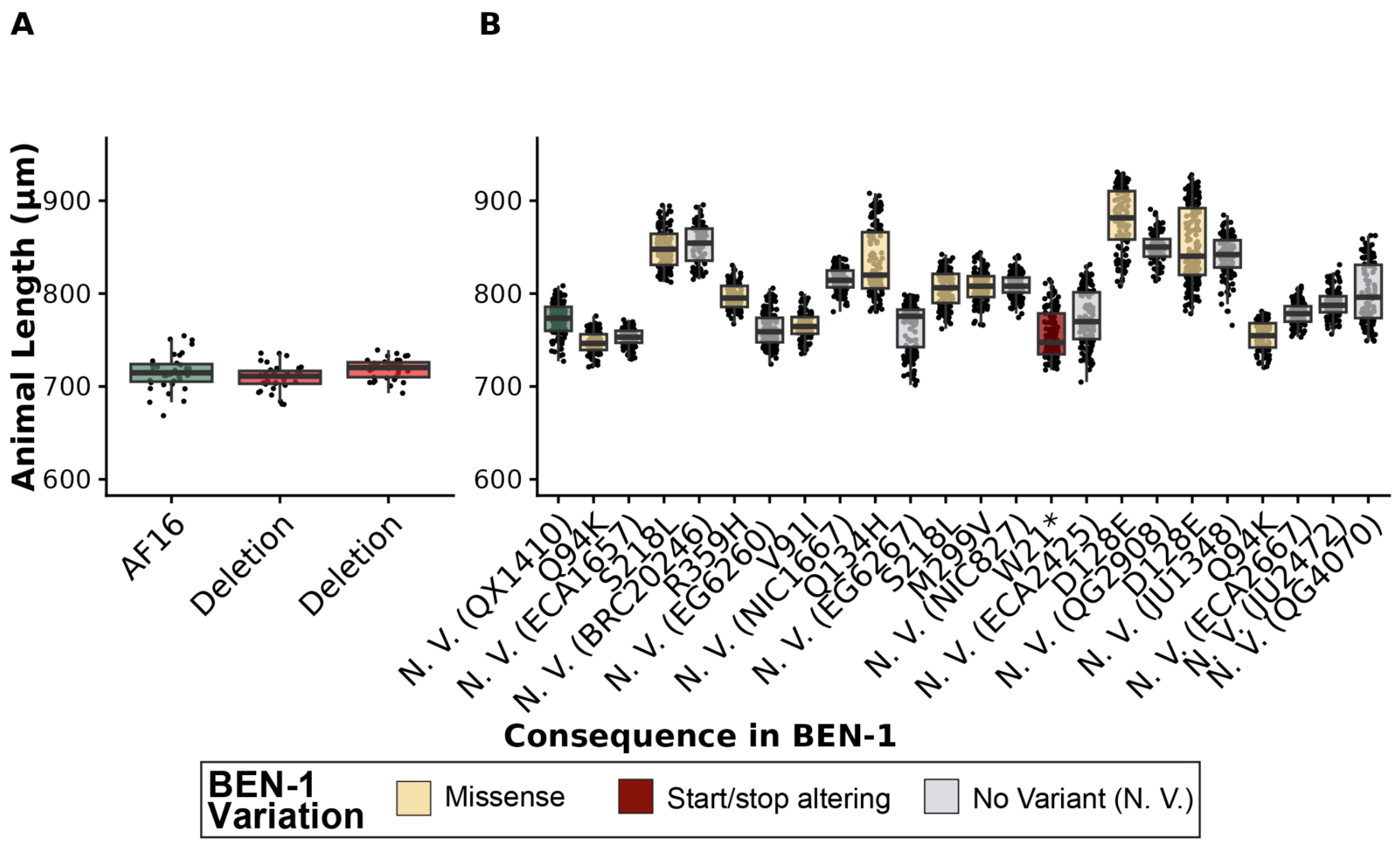


**S5 Fig. HTLDAs for *C. briggsae* strains with high-impact BEN-1 variants in control conditions.** Median animal length values from populations of nematodes grown in DMSO are shown on the y-axis. Each point represents the median animal length from a well containing approximately 5-30 animals. Data are shown as Tukey box plots with the median as a solid horizontal line, the top and bottom of the box representing the 75th and 25th quartiles, respectively. The top whisker is extended to the maximum point that is within a 1.5 interquartile range from the 75th quartile. The bottom whisker is extended to the minimum point that is within the 1.5 interquartile range from the 25th quartile. Results for **(A)** the AF16 reference strain (green) and two strains each with an independent *ben-1* deletion in the AF16 background (ECA3953 and ECA3954) (red), and **(B)** all wild *C. briggsae* strains with unique high-impact variants in *ben-1* are sorted by their relative resistance to ABZ based on median animal length. No variant (N. V.) strains (gray) paired with strains that have a high-impact variant in a beta-tubulin gene are shown alongside each corresponding strain with a high-impact variant in a beta-tubulin gene. Wild *C. briggsae* strains are colored by beta-tubulin variant status.

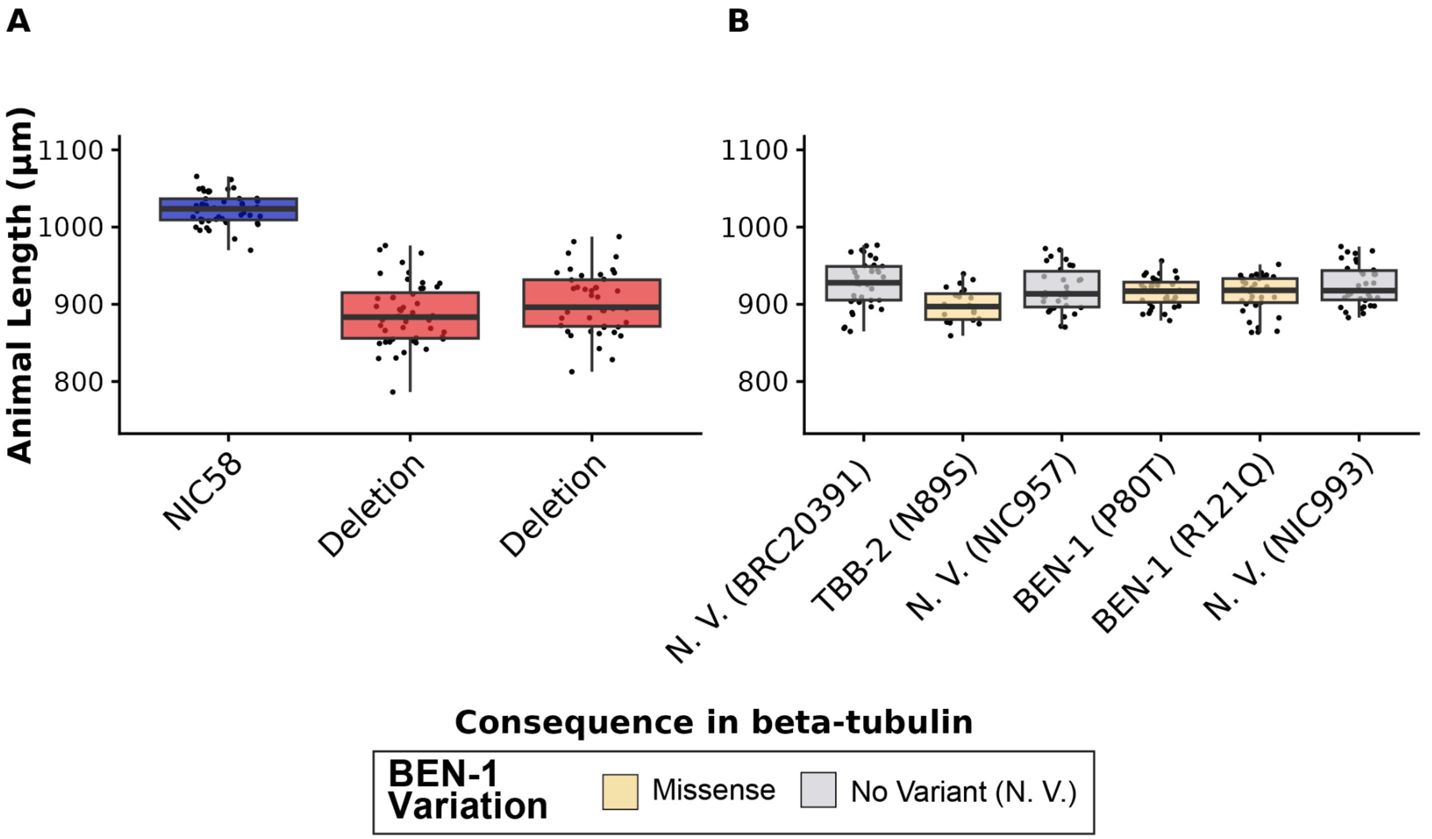


**S6 Fig. HTLDAs for *C. tropicalis* strains with high-impact TBB-2 or BEN-1 variants with paired predicted susceptible strains in control conditions.**

Median animal length values from populations of nematodes grown in DMSO are shown on the y-axis. Each point represents the median animal length from a well containing approximately 5-30 animals. Data are shown as Tukey box plots with the median as a solid horizontal line, the top and bottom of the box representing the 75th and 25th quartiles, respectively. The top whisker is extended to the maximum point that is within a 1.5 interquartile range from the 75th quartile. The bottom whisker is extended to the minimum point that is within the 1.5 interquartile range from the 25th quartile. Results for **(A)** the NIC58 reference strain (blue) and two strains each with an independent *ben-1* deletion in the NIC58 background (ECA4247 and ECA4248) (red), and **(B)** all wild *C. tropicalis* strains with unique high-impact variants in *ben-1* or *tbb-2* are sorted by their relative resistance to ABZ based on median animal length. No variant (N. V.) strains (gray) paired with strains that have a high-impact variant in a beta-tubulin gene are shown alongside each corresponding strain with a high-impact variant in a beta-tubulin gene. Wild *C. tropicalis* strains are colored by beta-tubulin variant status.

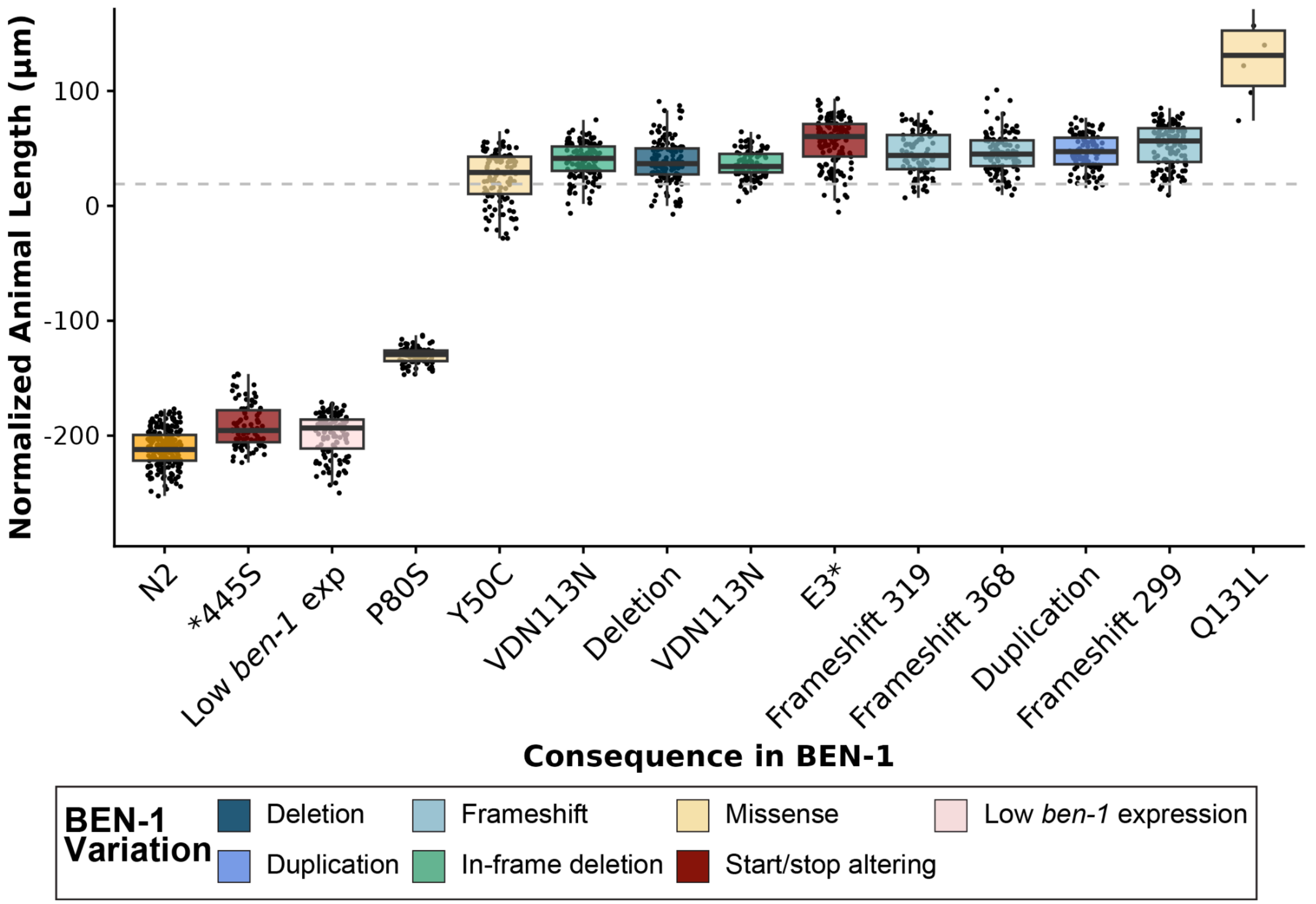


**S7 Fig. HTLDAs for each *C. elegans* strain with a high-impact variant in BEN-1 in albendazole**.
The regressed median animal length values for populations of nematodes grown in 30 µM albendazole (ABZ) are shown on the y-axis. Each point represents the normalized median animal length value of a well containing approximately 5-30 animals. Data are shown as Tukey box plots with the median as a solid horizontal line, and the top and bottom of the box representing the 75th and 25th quartiles, respectively. The top whisker is extended to the maximum point that is within the 1.5 interquartile range from the 75th quartile. The bottom whisker is extended to the minimum point that is within the 1.5 interquartile range from the 25th quartile. The gray dashed line marks the *C. elegans* resistance threshold, defined as two standard deviations below the mean of the *ben-1* deletion strain in the N2 reference strain background. Results for the N2 reference strain (orange) and all wild *C. elegans* strains with unique high-impact variants in *ben-1* are sorted by their relative resistance to ABZ based on median animal length. Wild *C. elegans* strains are colored by beta-tubulin variant status.

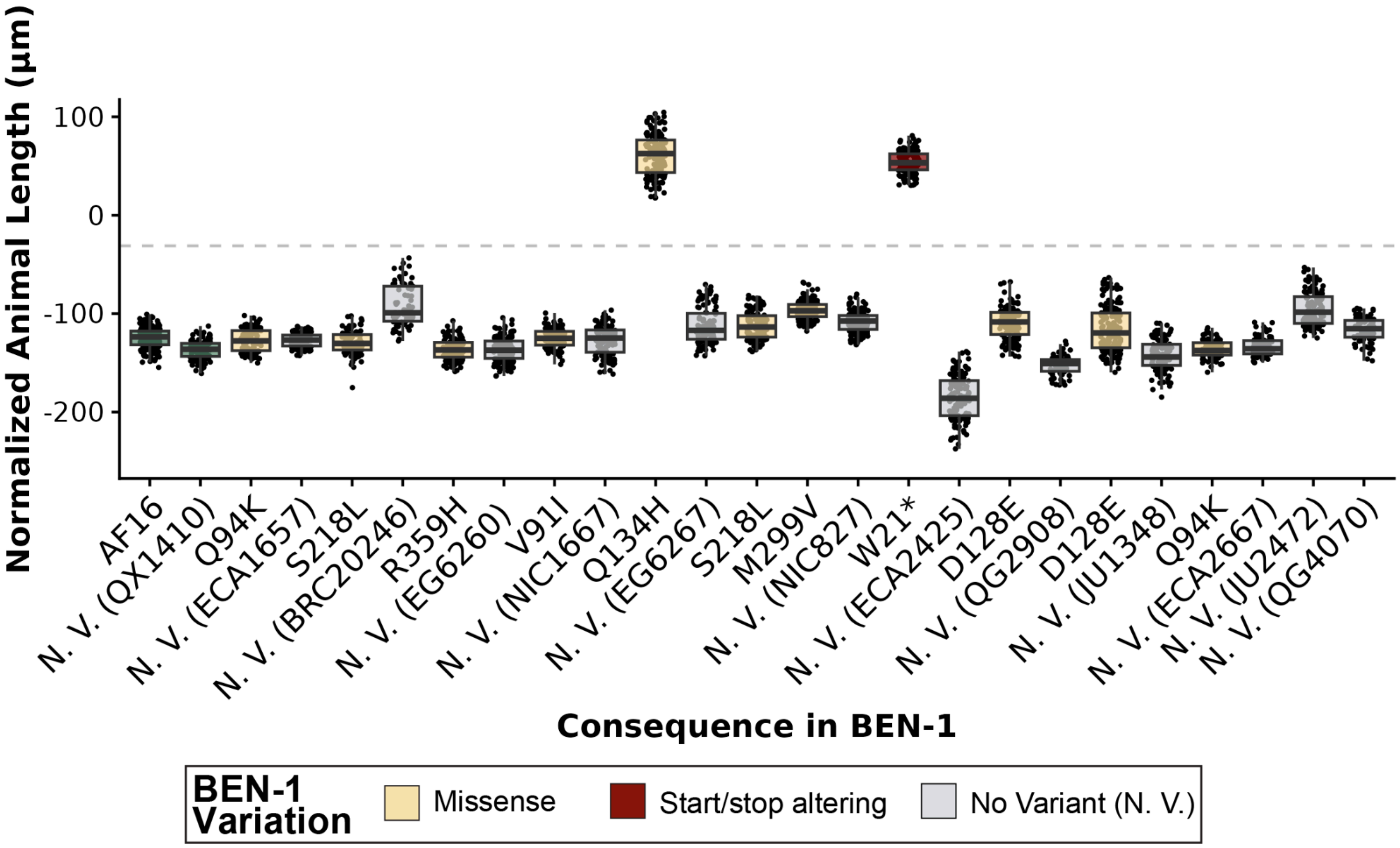


**S8 Fig. HTLDAs *C. briggsae* strains with a high-impact variant in BEN-1 with paired predicted susceptible strains in albendazole.**

The regressed median animal length values for populations of nematodes grown in 30 µM albendazole (ABZ) are shown on the y-axis. Each point represents the normalized median animal length value of a well containing approximately 5-30 animals. Strains are sorted by their relative resistance to ABZ based on median animal length. Data are shown as Tukey box plots with the median as a solid horizontal line, and the top and bottom of the box representing the 75th and 25th quartiles, respectively. The top whisker is extended to the maximum point that is within the 1.5 interquartile range from the 75th quartile. The bottom whisker is extended to the minimum point that is within the 1.5 interquartile range from the 25th quartile. The gray dashed line marks the *C. briggsae* resistance threshold, defined as two standard deviations below the mean of the *ben-1* deletion strain in the AF16 reference strain background. No variant (N. V.) strains (gray) paired with strains that have a high-impact variant in the *ben-1* gene are shown alongside each corresponding strain with a high-impact variant in *ben-1*. Wild *C. briggsae* strains are colored by beta-tubulin variant status.

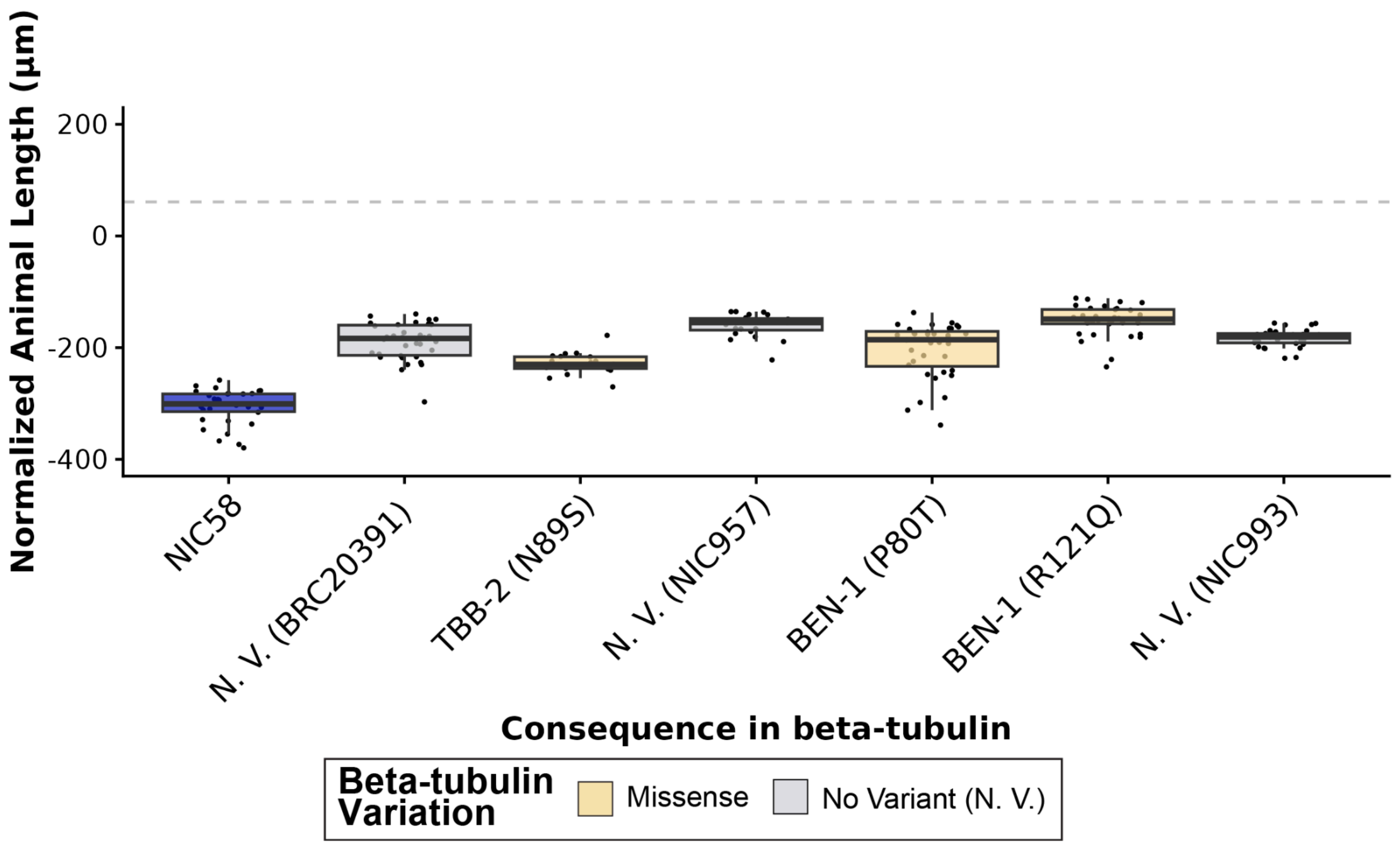


**S9 Fig. HTLDAs for *C. tropicalis* strains with high-impact variants in TBB-2 or BEN-1 with paired predicted susceptible strains in albendazole.**
The regressed median animal length values for populations of nematodes grown in 30 µM albendazole (ABZ) are shown on the y-axis. Each point represents the normalized median animal length value of a well containing approximately 5-30 animals. Data are shown as Tukey box plots with the median as a solid horizontal line, and the top and bottom of the box representing the 75th and 25th quartiles, respectively. The top whisker is extended to the maximum point that is within the 1.5 interquartile range from the 75th quartile. The bottom whisker is extended to the minimum point that is within the 1.5 interquartile range from the 25th quartile. The gray dashed line marks the *C. tropicalis* resistance threshold, defined as two standard deviations below the mean of the *ben-1* deletion strain in the NIC58 reference strain background. No variant (N. V.) strains (gray) paired with strains that have a high-impact variant in the *tbb-2* or *ben-1* genes are shown alongside each corresponding strain with a high-impact variant in *tbb-2* or *ben-1*. Wild *C. tropicalis* strains are colored by beta-tubulin variant status.

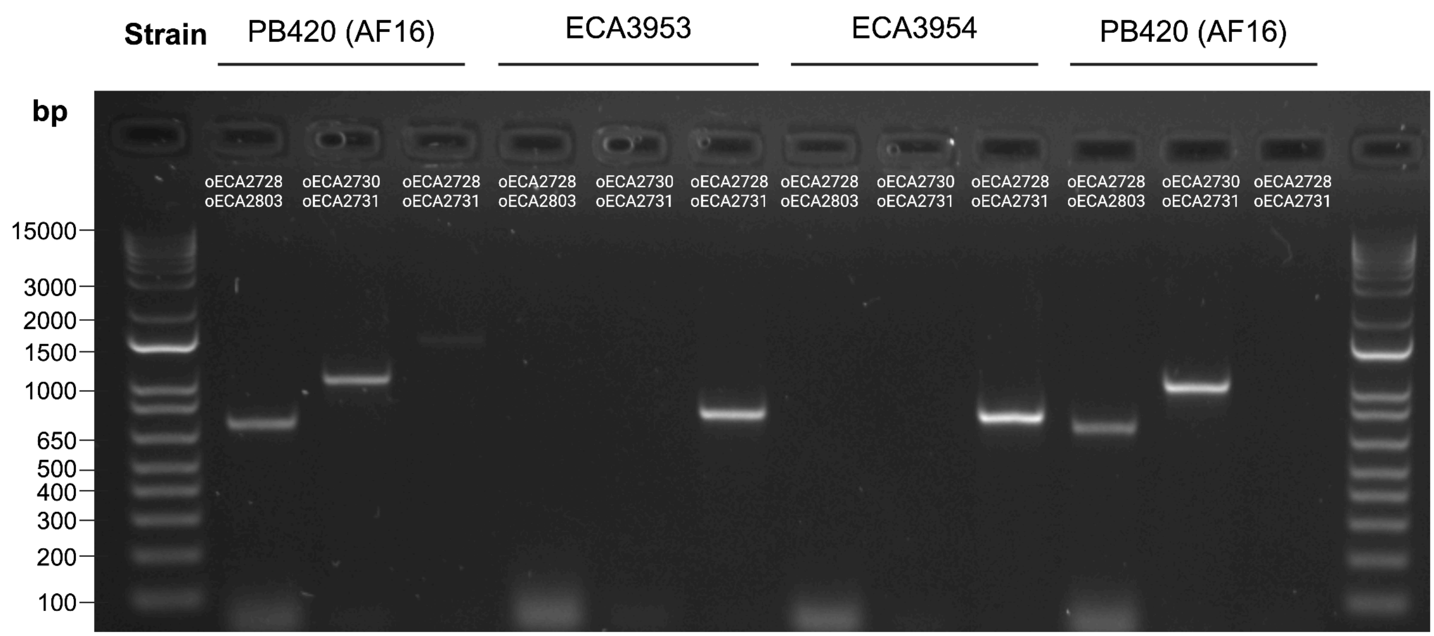


**S10 Fig. PCR confirmation of the *ben-1* deletion in the *C. briggsae* reference strain background, AF16**
Three primer pairs were used to confirm the deletion of *ben-1* in the *C. briggsae* reference strain, AF16. The oECA2728 (external) and oECA2803 (internal) primers flank either side of the guide region on the 5' end. The oECA2730 (internal) and oECA2731 (external) primers flank either side of the guide region on the 3' end. The oECA2728 and oECA2731 primers flank the outside of the *ben-1* region to be deleted. The wild-type (AF16) region spans 1383 base pairs (bp), while the *ben-1* deletion is reduced to 732 bp. The top of the gel is labeled by the three strains: PB420 (AF16) and the two independently edited *ben-1* deletion strains in the AF16 background (ECA3953 and ECA3954). Each well of the gel is labeled by the primer pair used. The Invitrogen 1 Kb Plus DNA Ladder is shown on each side of the gel.

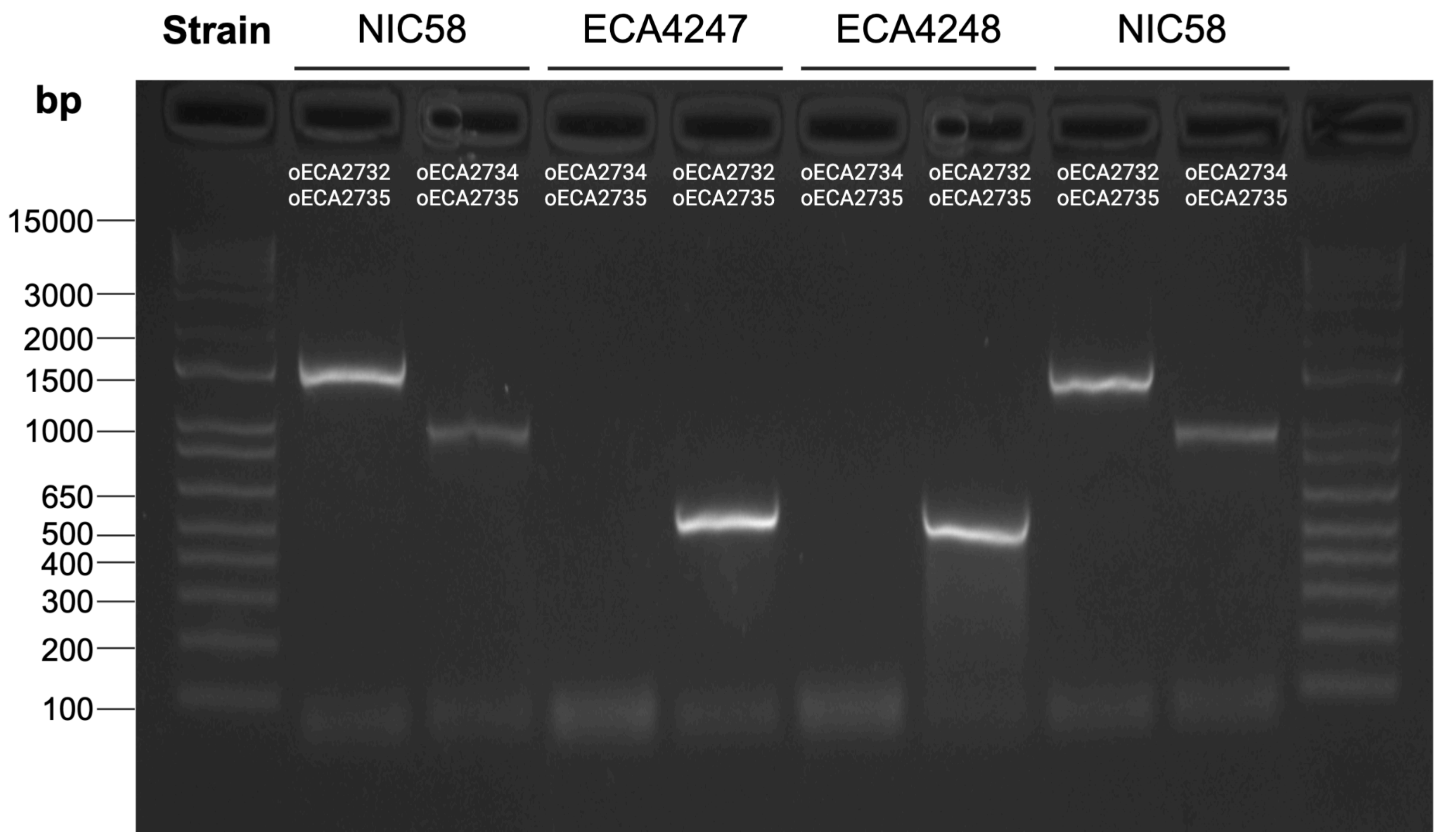


**S11 Fig. PCR confirmation of the *ben-1* deletion in the *C. tropicalis* reference strain background, NIC58**
Three primer pairs were used to confirm the deletion of *ben-1* in the *C. tropicalis* reference strain background, NIC58. The oECA2734 (internal) and oECA2735 (external) primers flank either side of the guide region on the 3' end. The oECA2732 and oECA2735 primers flank the *ben-1* region to be deleted. The wild-type (NIC58) region spans 1538 base pairs (bp), and the *ben-1* deletion reduces the region to 513 bp. The top of the gel is labeled by the three strains: NIC58 and the two independently edited *ben-1* deletion strains in the NIC58 background (ECA4247 and ECA4248). Each well of the gel is labeled by the primer pair used. The Invitrogen 1 Kb Plus DNA Ladder is shown on each side of the gel.

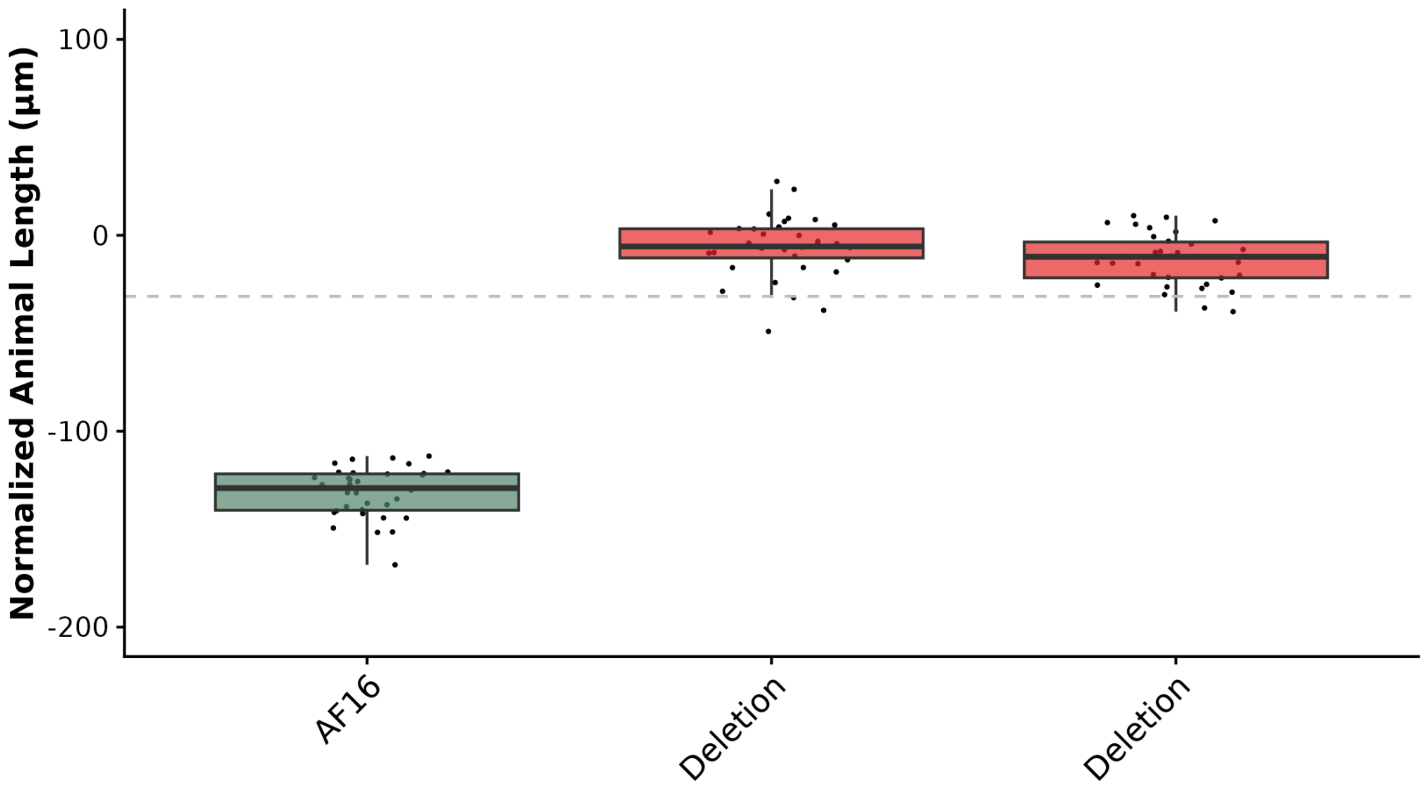


**S12 Fig. High-throughput larval development assays for two independently edited *C. briggsase* AF16 strains with a loss of *ben-1***

The regressed median animal length values for populations of nematodes grown in 30 µM albendazole (ABZ) are shown on the y-axis. Each point represents the normalized median animal length value of a well containing approximately 5-30 animals. Data are shown as Tukey box plots with the median as a solid horizontal line, and the top and bottom of the box representing the 75th and 25th quartiles, respectively. The top whisker is extended to the maximum point that is within the 1.5 interquartile range from the 75th quartile. The bottom whisker is extended to the minimum point that is within the 1.5 interquartile range from the 25th quartile. The gray dashed line marks the *C. briggsae* resistance threshold, defined as two standard deviations below the mean of the *ben-1* deletion strain (ECA3953) in the AF16 reference strain background. Results are shown for the AF16 reference strain (green) and two independently edited strains with a *ben-1* deletion in the AF16 background (ECA3953 and ECA3954) (red).

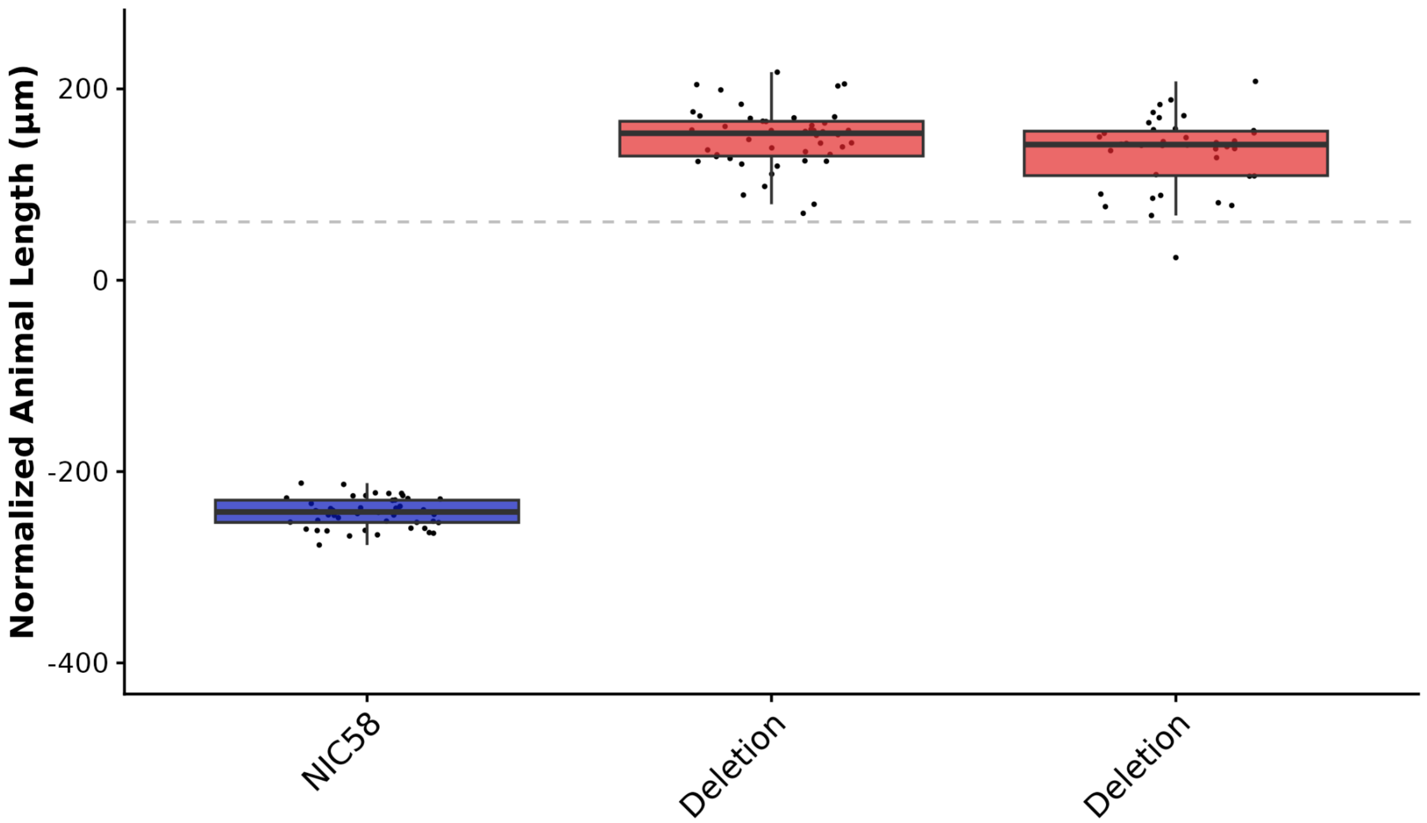


**S13 Fig. High-throughput larval development assays for two independently edited *C. tropicalis* NIC58 strains with a loss of *ben-1***

The regressed median animal length values for populations of nematodes grown in 30 µM albendazole (ABZ) are shown on the y-axis. Each point represents the normalized median animal length value of a well containing approximately 5-30 animals. Data are shown as Tukey box plots with the median as a solid horizontal line, and the top and bottom of the box representing the 75th and 25th quartiles, respectively. The top whisker is extended to the maximum point that is within the 1.5 interquartile range from the 75th quartile. The bottom whisker is extended to the minimum point that is within the 1.5 interquartile range from the 25th quartile. The gray dashed line marks the *C. tropicalis* resistance threshold, defined as two standard deviations below the mean of the *ben-1* deletion strain (ECA24248) in the NIC58 reference strain background. Results are shown for the NIC58 reference strain (blue) and two independently edited strains with a *ben-1* deletion in the NIC58 background (ECA4247 and ECA4248) (red).

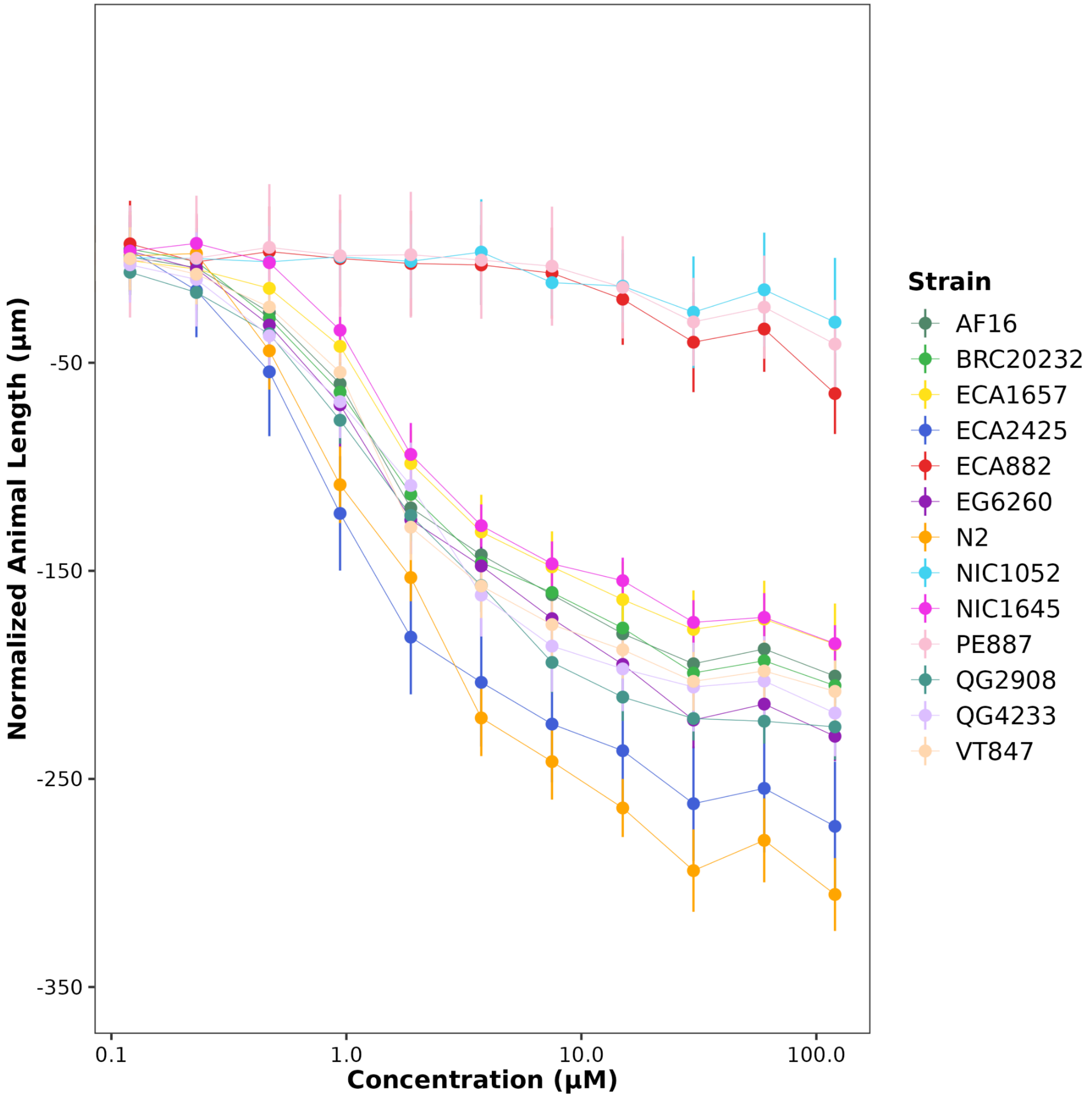


**S14 Fig. Dose-response curves for *C. briggsae* strains in albendazole**

Normalized animal lengths (y-axis) are plotted for each strain as a function of the dose of albendazole (ABZ) in the high-throughput larval development assay (x-axis). Strains are denoted by color. Lines extending vertically from points represent the standard deviation from the mean response. Statistical normalization of animal lengths is described in *Methods*. *C. elegans* strains N2 and ECA882 were added for ABZ-susceptible and ABZ-resistant controls, respectively.

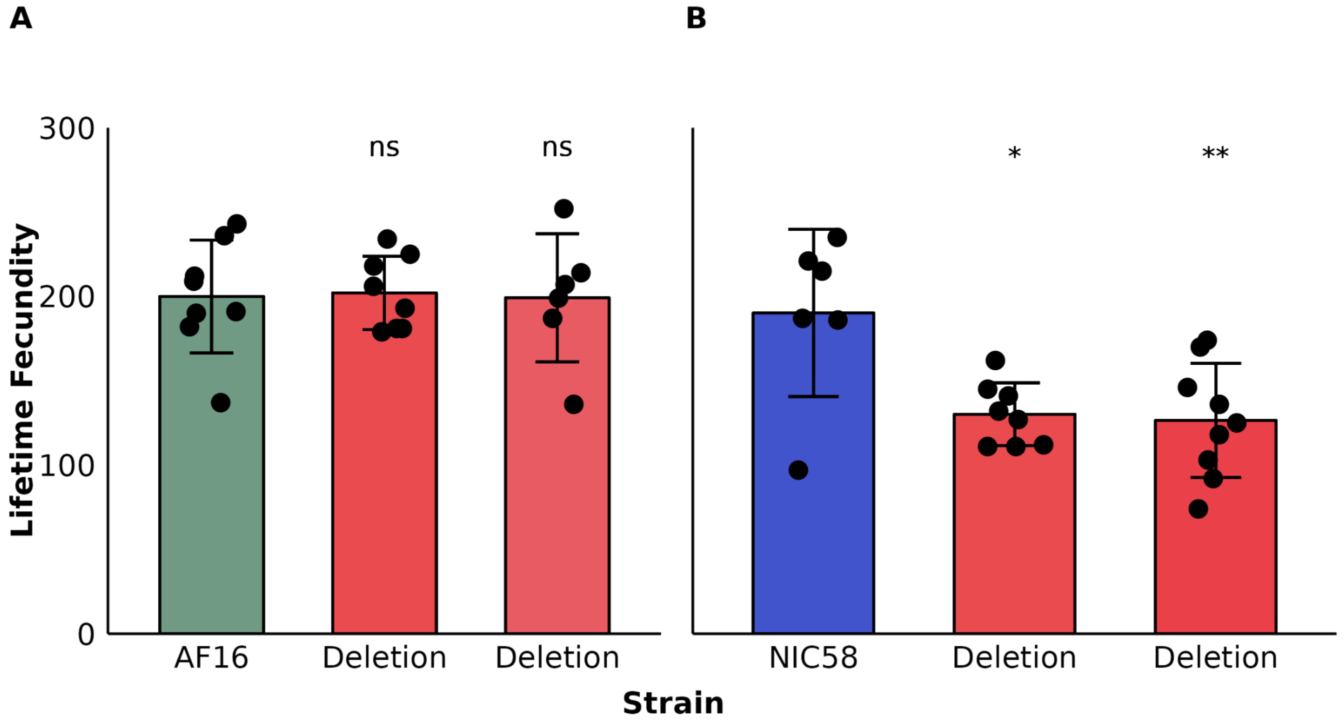


**S15 Fig. Variation in lifetime fecundity between the *C. briggsae* (AF16) and *C. tropicalis* (NIC58) reference strains and strains with a loss of *ben-1* in the AF16 and NIC58 reference strain backgrounds** Bar plots for the lifetime fecundity, y-axis, for each strain on the x-axis are shown. Error bars show the standard deviation of lifetime fecundity among 6 to 9 replicates. **(A)** A comparison of lifetime fecundities between the *C. briggsae* laboratory reference strain AF16 (green) and the two independently edited *C. briggsae* AF16 strains with a loss of *ben-1* (red). **(B)** A comparison between the *C. tropicalis* laboratory reference strain NIC58 (blue) and two independently edited *C. tropicalis* NIC58 strains with a loss of *ben-1* (red). Statistical significance was determined using Tukey HSD. Significance of each comparison is shown above each comparison pair ($p > 0.05$ = ns, $p < 0.05$ = *, $p < 0.01$ = ** Tukey HSD).

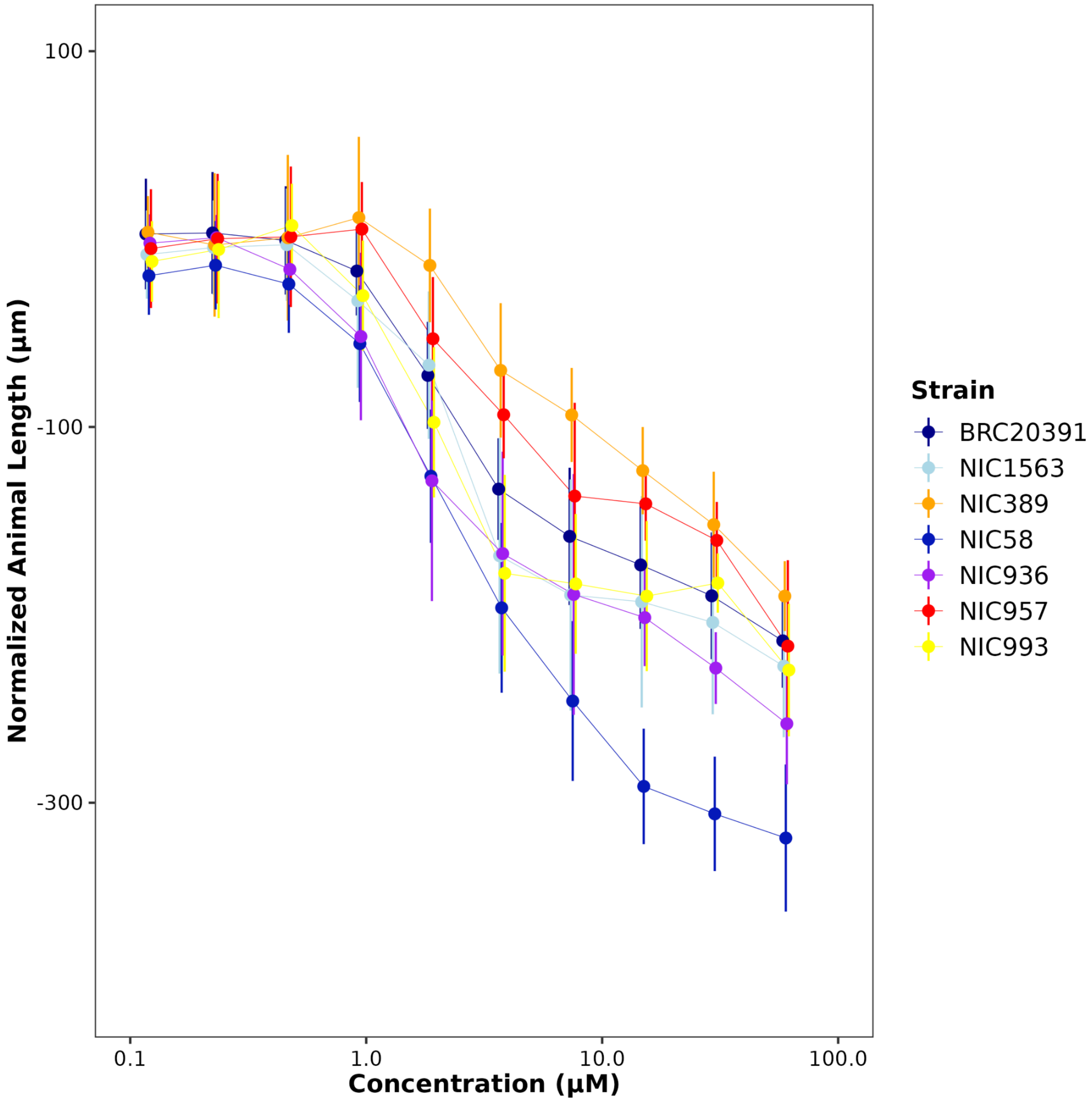


**S16 Fig. Dose-response curves for *C. tropicalis* strains in albendazole**

Normalized animal lengths (y-axis) are plotted for each strain as a function of the dose of albendazole (ABZ) in the high-throughput larval development assay (x-axis). Strains are denoted by color. Lines extending vertically from points represent the standard deviation from the mean response. Statistical normalization of animal lengths is described in *Materials and Methods*.

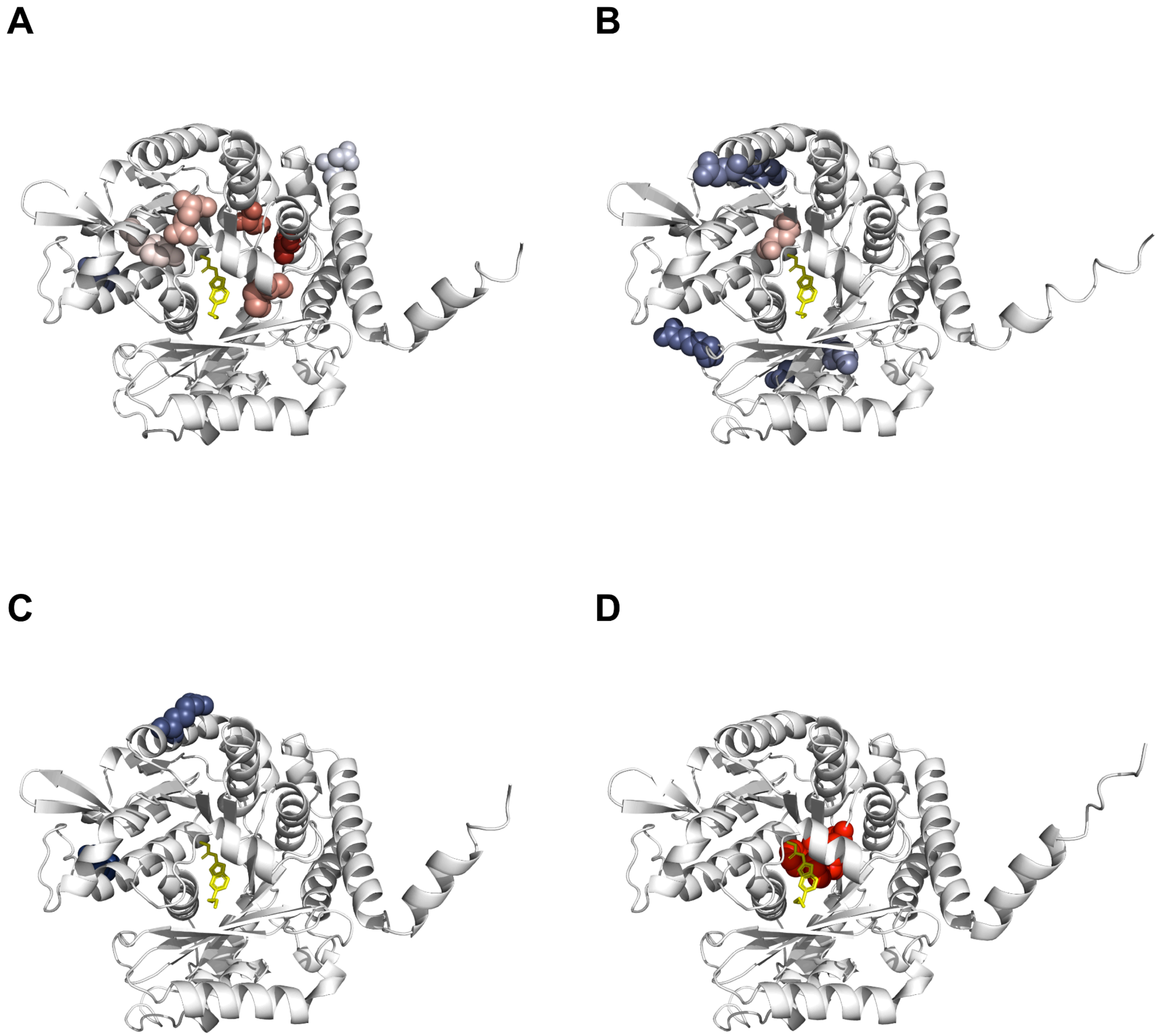


**S17 Fig**. **Structure of beta-tubulin orthologs bound to albendazole**

AlphaFold3 models of BEN-1 orthologs bound to albendazole (ABZ) are shown for **(A)** *Cel*-BEN-1, **(B)** *Cbr*-BEN-1, **(C)** *Ctr*-BEN-1, and **(D)** *Hcon*-TBB-ISO-1. ABZ is colored yellow. Residues impacted by amino acid substitutions are colored by resistance (red) or sensitivity (blue).

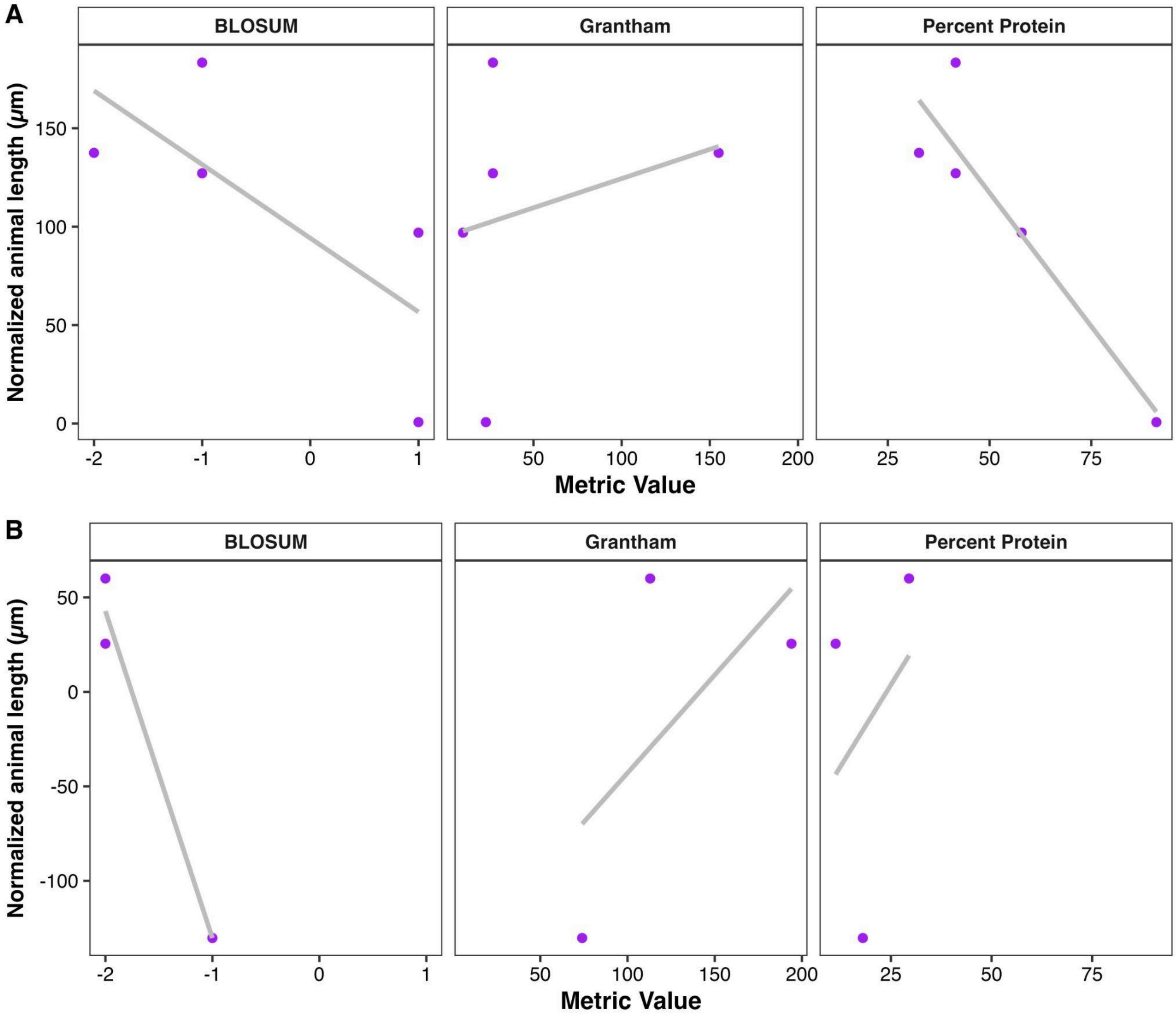


**S18 Fig. Relationship between missense substitutions in BEN-1 and albendazole response in *C. elegans* strains.** Scatterplots show the relationship between normalized median animal length (y-axis) and three amino acid substitution scoring metrics (x-axis): BLOSUM62 ($R^2$ = 0.97, *p*-value = 0.11), Grantham ($R^2$ = 0.39, *p*-value = 0.57), and percent protein ($R^2$ = 0.1, *p*-value = 0.8). Each point represents a *C. elegans* strain with a missense substitution in BEN-1. Gray lines indicate the linear regression fit for these models. **(A)** Strains phenotyped for ABZ response in previous assays are plotted (Hahnel et al. 2018; Shaver et al. 2024). **(B)** Strains phenotyped in the assays performed for this study are plotted.

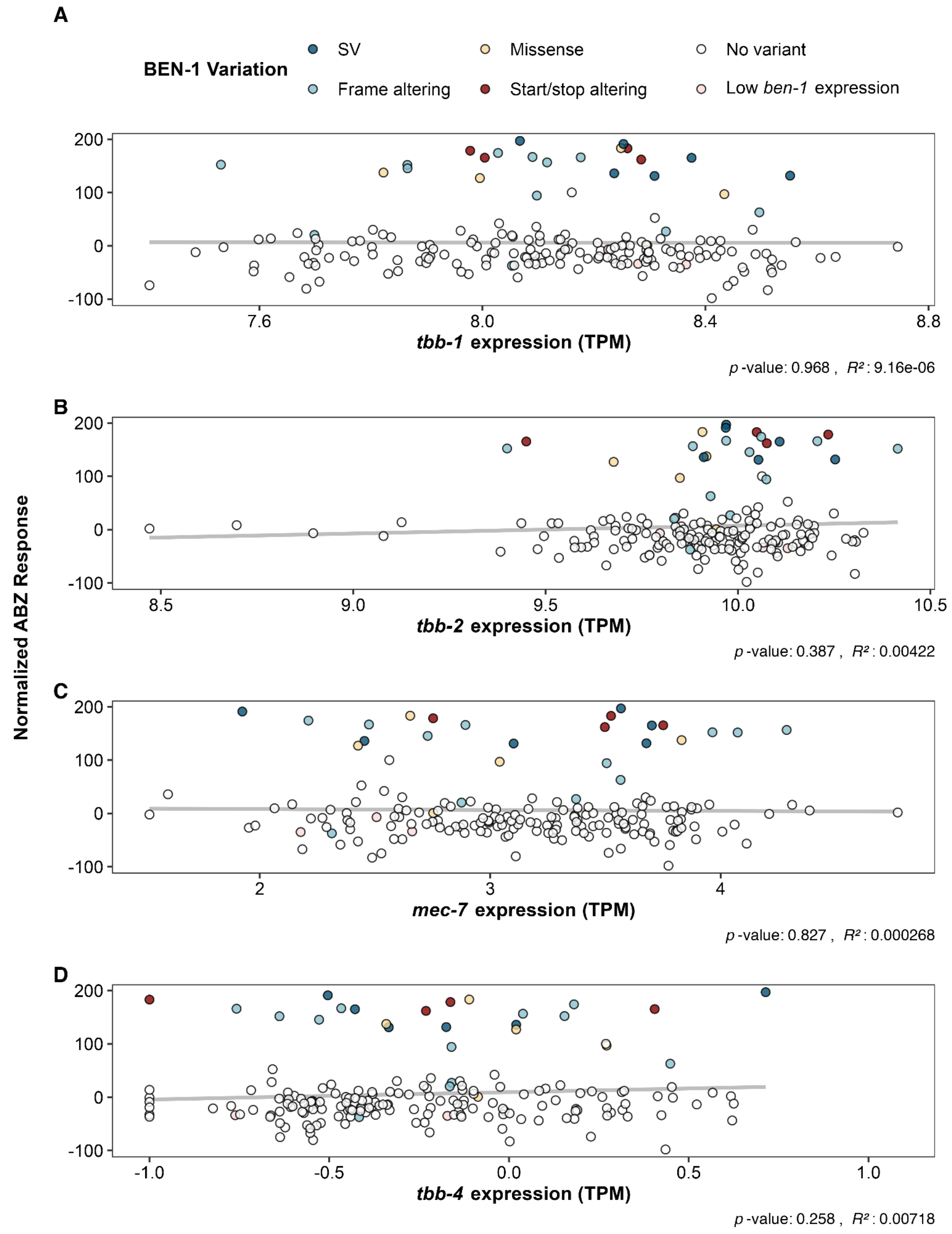


**S19 Fig. The relationship between *C. elegans* beta-tubulin expression and albendazole response**
Scatterplot of the relationship between **(A)** *tbb-1*, **(B)** *tbb-2*, **(C)** *mec-7*, and **(D)** *tbb-4* expression levels and normalized albendazole (ABZ) response across *C. elegans* wild strains. Each point represents a strain phenotyped for ABZ response in previous publications (Hahnel *et al.*, 2018; Shaver *et al.*, 2024) with *tbb-1, tbb-2*, *mec-7*, and *tbb-4* expression data (Zhang et al. 2022). The *tbb-1*, *tbb-2*, *mec-7*, and *tbb-4* expression levels measured in transcripts per million (TPM) are displayed on the x-axis. The normalized ABZ response values adjusted for assay-specific effects are displayed on the y-axis. The gray line represents the linear regression fit between beta-tubulin gene expression and normalized response, with the linear model's coefficient of determination ($R^2$). Data points are colored based on the predicted functional consequence of their *ben-1* alleles (*i.e.*, large structural variant (SV), frameshift, missense substitution, disrupted start/stop sequence, no high-impact variant, or low *ben-1* expression).

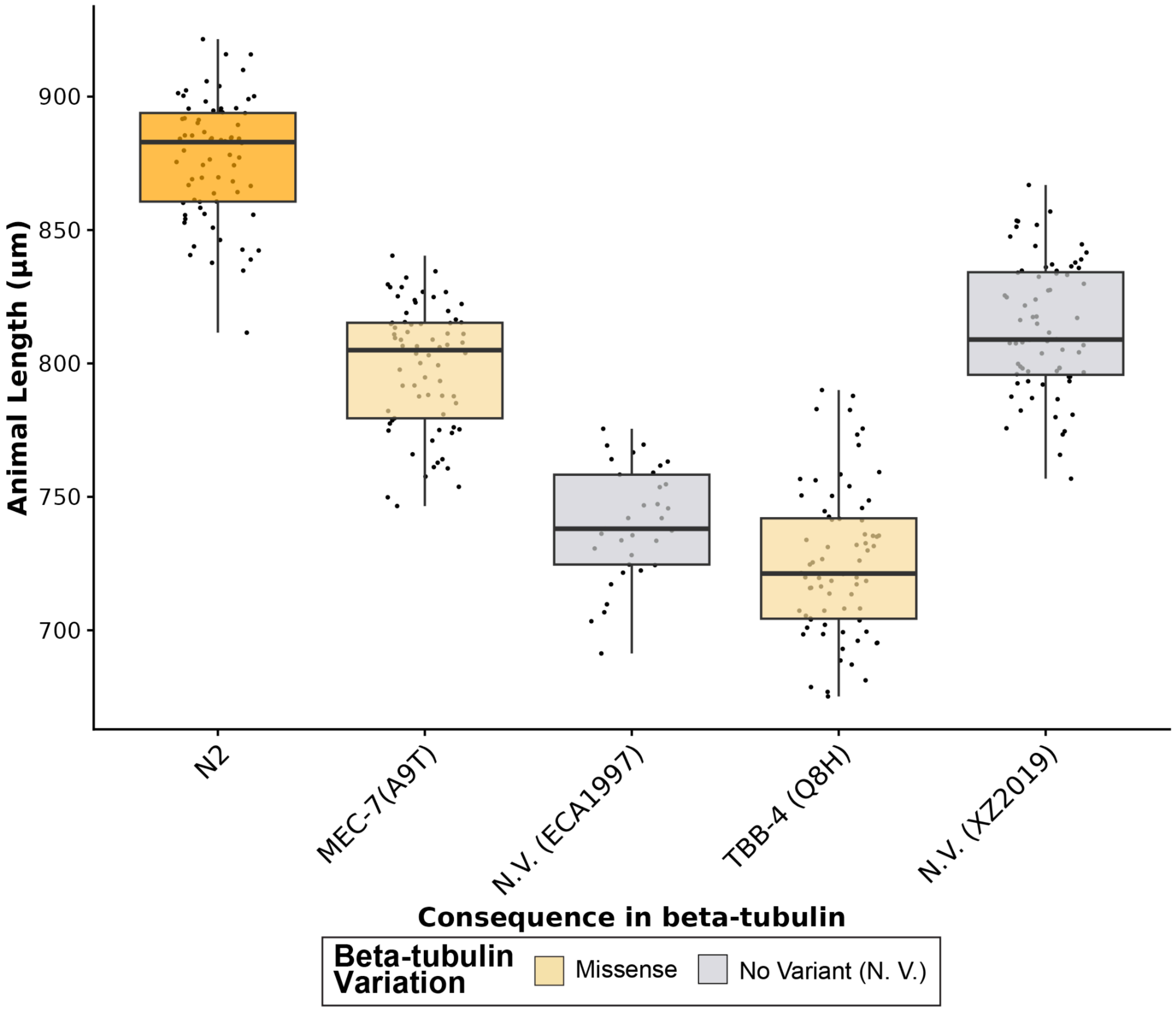


**S20 Fig. High-throughput larval development assays for each *C. elegans* strain with a high-impact variant in MEC-7 or TBB-4 with paired predicted susceptible strains in control conditions**

Median animal length values from populations of nematodes grown in DMSO are shown on the y-axis. Each point represents the median animal length from a well containing approximately 5-30 animals. Data are shown as Tukey box plots with the median as a solid horizontal line, the top and bottom of the box representing the 75th and 25th quartiles, respectively. The top whisker is extended to the maximum point that is within a 1.5 interquartile range from the 75th quartile. The bottom whisker is extended to the minimum point that is within the 1.5 interquartile range from the 25th quartile. No variant (N. V.) strains (gray) paired with strains that have a high-impact variant in a beta-tubulin gene are shown alongside each corresponding strain with a high-impact variant in a beta-tubulin gene. Wild *C. elegans* strains are colored by beta-tubulin variant status.

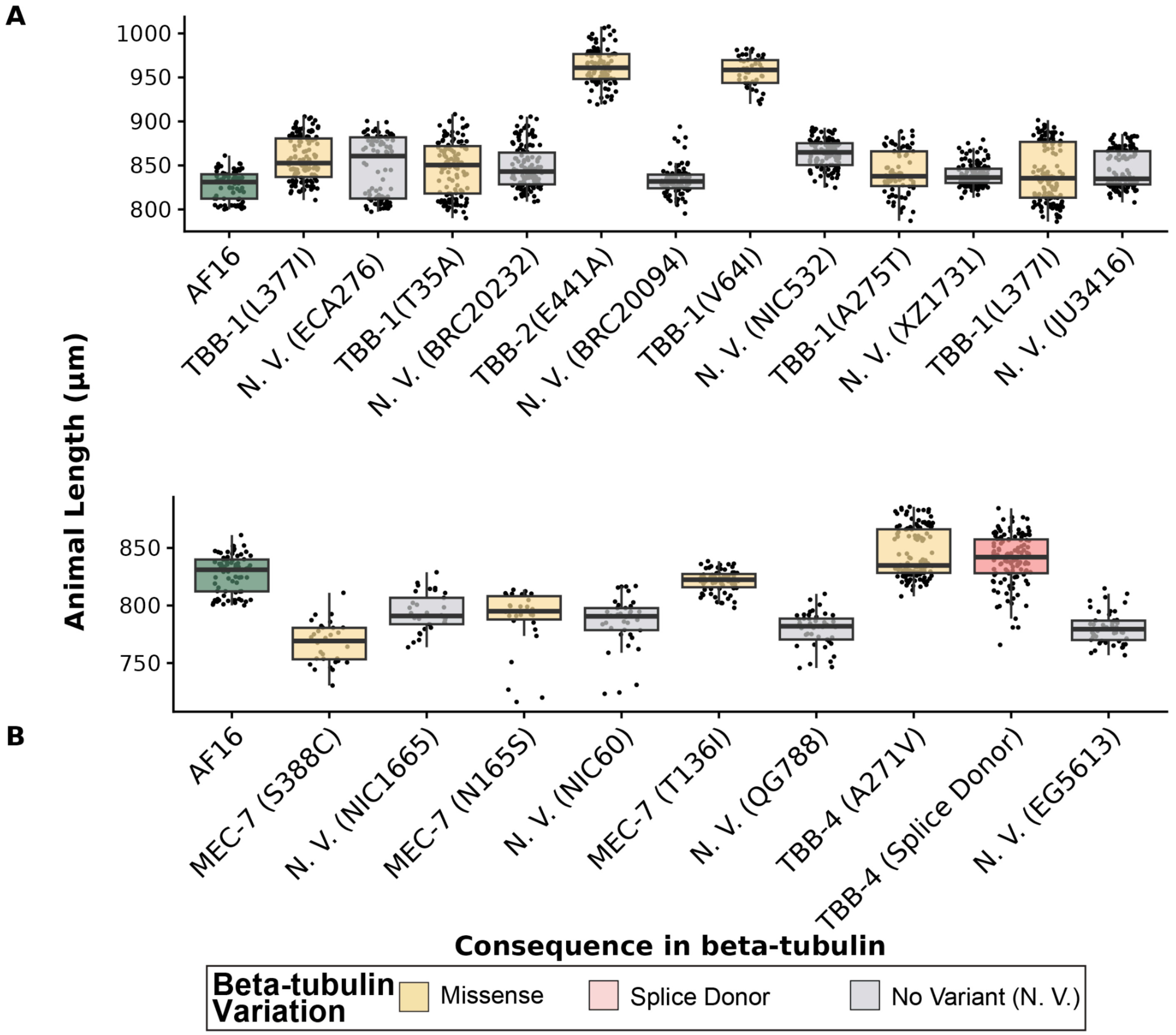


**S21 Fig. High-throughput larval development assays for each *C. briggsae* strain with a high-impact variant in TBB-1, TBB-2, MEC-7, or TBB-4 with paired predicted susceptible strains in control conditions**

Median animal length values from populations of nematodes grown in DMSO are shown on the y-axis. Each point represents the median animal length from a well containing approximately 5-30 animals. Data are shown as Tukey box plots with the median as a solid horizontal line, the top and bottom of the box representing the 75th and 25th quartiles, respectively. The top whisker is extended to the maximum point that is within a 1.5 interquartile range from the 75th quartile. The bottom whisker is extended to the minimum point that is within the 1.5 interquartile range from the 25th quartile. No variant (N. V.) strains (gray) paired with strains that have a high-impact variant in a beta-tubulin gene are shown alongside each corresponding strain with a high-impact variant in a beta-tubulin gene. Wild *C. briggsae* strains are colored by beta-tubulin variant status.

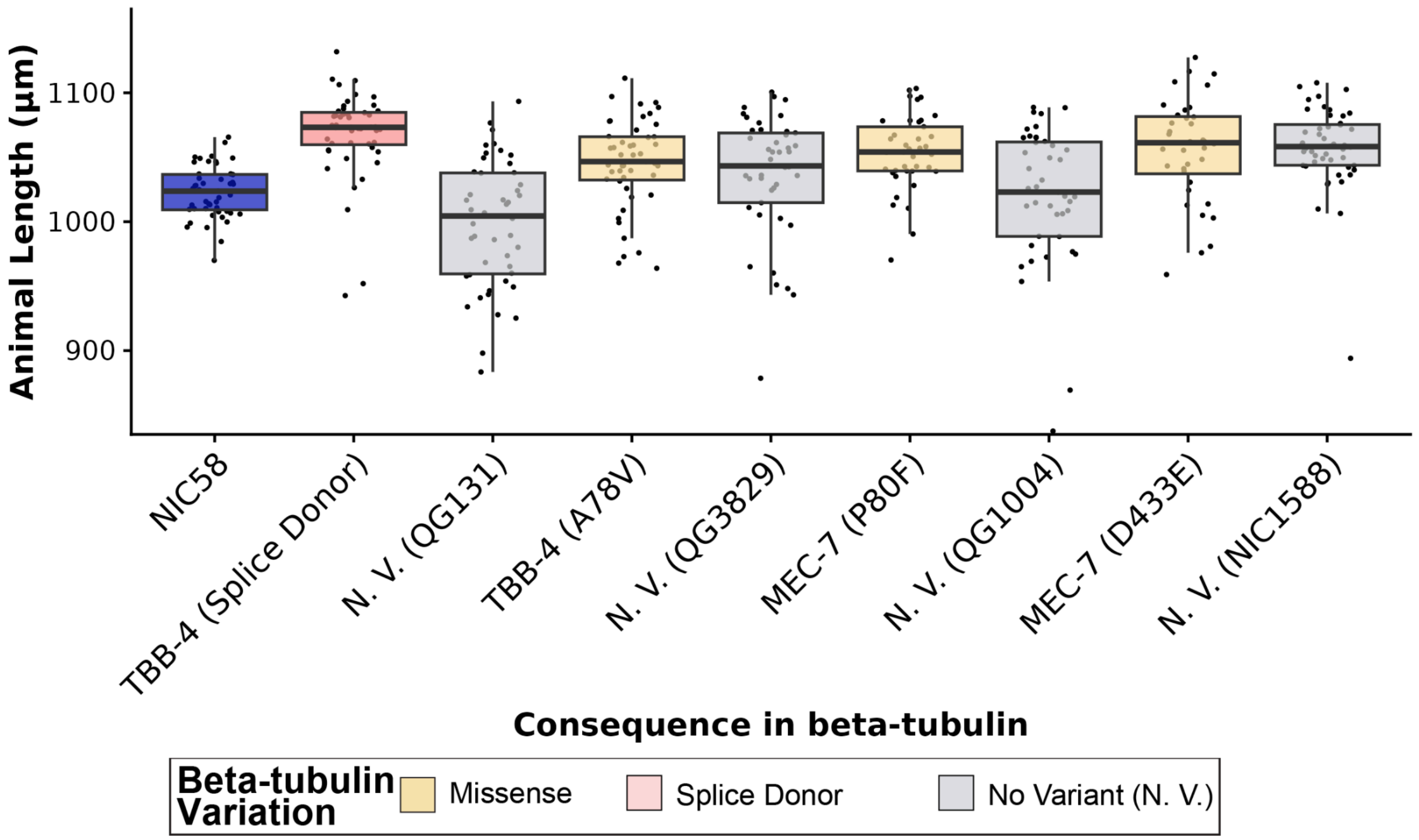


**S22 Fig. High-throughput larval development assays for each *C. tropicalis* strain with a high-impact variant in MEC-7 or TBB-4 with paired predicted susceptible strains in control conditions**

Median animal length values from populations of nematodes grown in DMSO are shown on the y-axis. Each point represents the median animal length from a well containing approximately 5-30 animals. Data are shown as Tukey box plots with the median as a solid horizontal line, the top and bottom of the box representing the 75th and 25th quartiles, respectively. The top whisker is extended to the maximum point that is within a 1.5 interquartile range from the 75th quartile. The bottom whisker is extended to the minimum point that is within the 1.5 interquartile range from the 25th quartile. No variant (N. V.) strains (gray) paired with strains that have a high-impact variant in a beta-tubulin gene are shown alongside each corresponding strain with a high-impact variant in a beta-tubulin gene. Wild *C. tropicalis* strains are colored by beta-tubulin variant status.

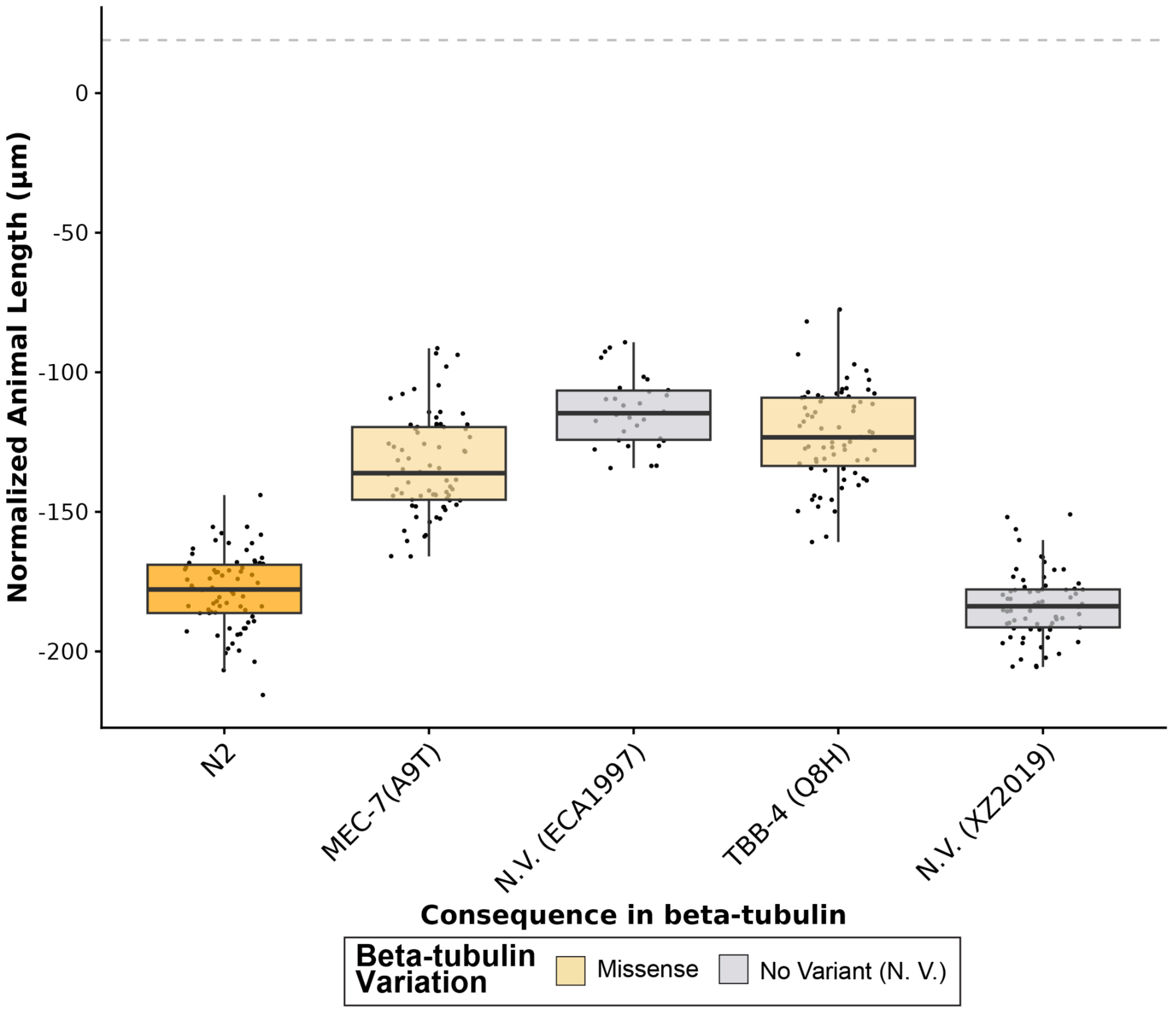


**S23 Fig. High-throughput larval development assays for each *C. elegans* strain with a high-impact variant in MEC-7 or TBB-4 in the presence of albendazole**

The regressed median animal length values for populations of nematodes grown in 30 µM albendazole (ABZ) are shown on the y-axis. Each point represents the normalized median animal length value of a well containing approximately 5-30 animals. Data are shown as Tukey box plots with the median as a solid horizontal line, and the top and bottom of the box representing the 75th and 25th quartiles, respectively. The top whisker is extended to the maximum point that is within the 1.5 interquartile range from the 75th quartile. The gray dashed line marks the *C. elegans* resistance threshold, defined as two standard deviations below the mean of the *ben-1* deletion strain in the N2 reference strain background. The bottom whisker is extended to the minimum point that is within the 1.5 interquartile range from the 25th quartile. No variant (N. V.) strains (gray) paired with strains that have a high-impact variant in a beta-tubulin gene are shown alongside each corresponding strain with a high-impact variant in a beta-tubulin gene. Wild *C. elegans* strains are colored by beta-tubulin variant status.

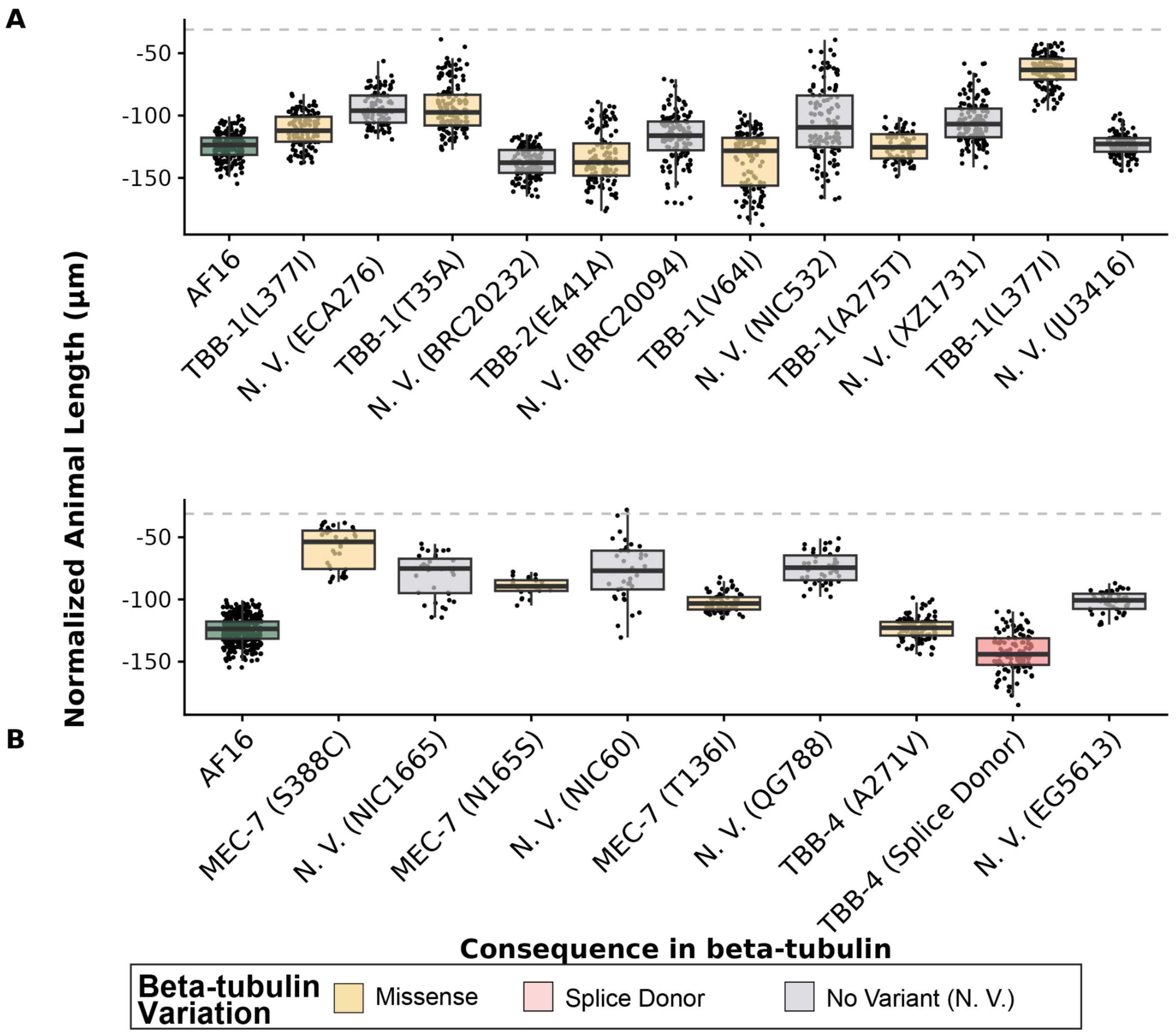


**S24 Fig. High-throughput larval development assays for each *C. briggsae* strain with a high-impact variant in TBB-2, MEC-7, or TBB-4 in the presence of albendazole**

The regressed median animal length values for populations of nematodes grown in 30 µM albendazole (ABZ) are shown on the y-axis. Each point represents the normalized median animal length value of a well containing approximately 5-30 animals. Data are shown as Tukey box plots with the median as a solid horizontal line, and the top and bottom of the box representing the 75th and 25th quartiles, respectively. The top whisker is extended to the maximum point that is within the 1.5 interquartile range from the 75th quartile. The bottom whisker is extended to the minimum point that is within the 1.5 interquartile range from the 25th quartile. The gray dashed line marks the *C. briggsae* resistance threshold, defined as two standard deviations below the mean of the *ben-1* deletion strain in the AF16 reference strain background. Results for the AF16 reference strain and all *C. briggsae* wild strains with unique high-impact variants in **(A)** TBB-1 and TBB-2 and **(B)** TBB-4 and MEC-7 are shown. No variant (N. V.) strains (gray) paired with strains that have a high-impact variant in a beta-tubulin gene are shown alongside each corresponding strain with a high-impact variant in a beta-tubulin gene. Wild *C. tropicalis* strains are colored by beta-tubulin variant status.

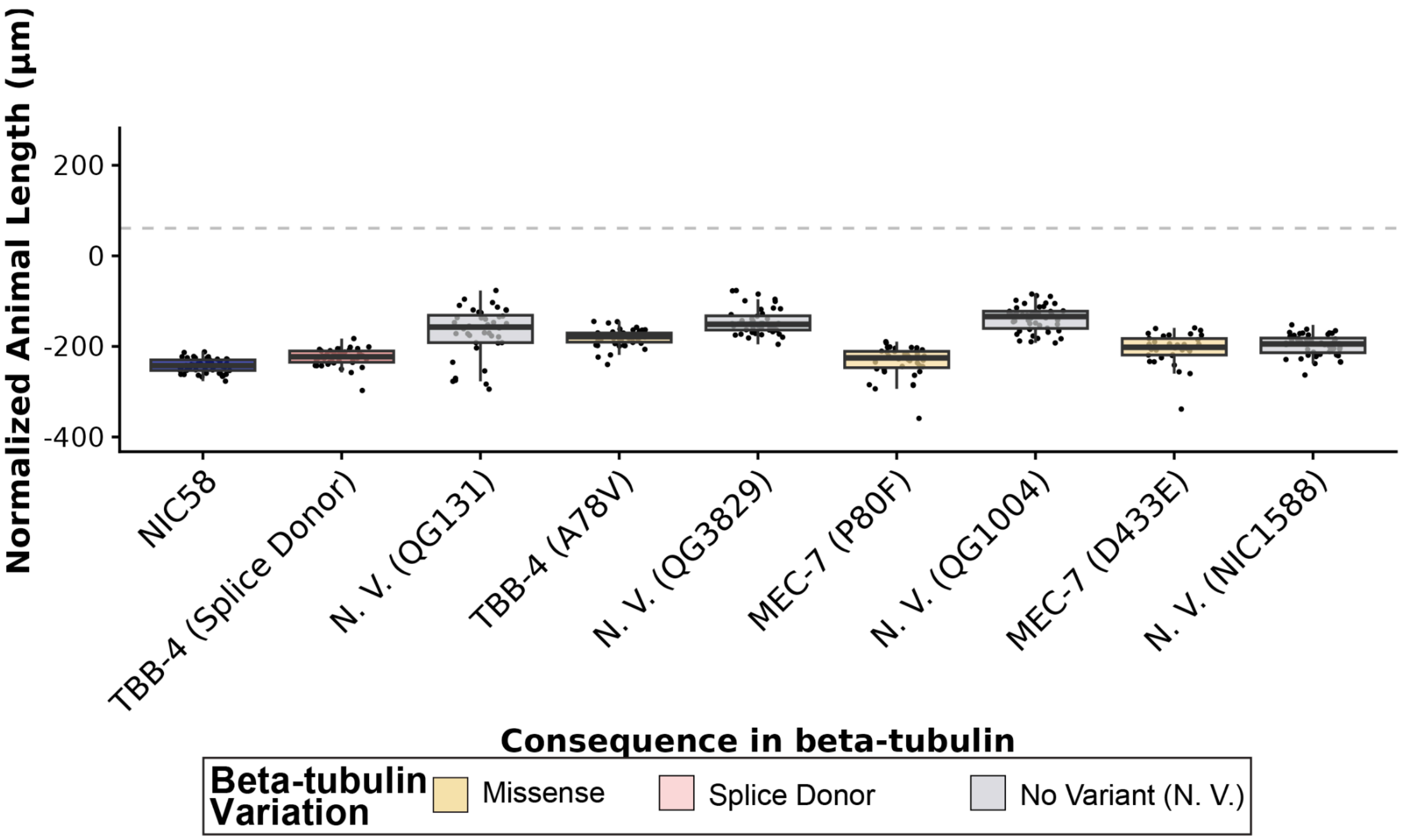


**S25 Fig. High-throughput larval development assays for each *C. tropicalis* strain with a high-impact variant in MEC-7 or TBB-4, and paired predicted susceptible strains in the presence of albendazole**
The regressed median animal length values for populations of nematodes grown in 30 µM albendazole (ABZ) are shown on the y-axis. Each point represents the normalized median animal length value of a well containing approximately 5-30 animals. Data are shown as Tukey box plots with the median as a solid horizontal line, and the top and bottom of the box representing the 75th and 25th quartiles, respectively. The top whisker is extended to the maximum point that is within the 1.5 interquartile range from the 75th quartile. The bottom whisker is extended to the minimum point that is within the 1.5 interquartile range from the 25th quartile. The gray dashed line marks the *C. tropicalis* resistance threshold, defined as two standard deviations below the mean of the *ben-1* deletion strain in the NIC58 reference strain background. No variant (N. V.) strains (gray) paired with strains that have a high-impact variant in a beta-tubulin gene are shown alongside each corresponding strain with a high-impact variant in a beta-tubulin gene. Wild *C. tropicalis* strains are colored by beta-tubulin variant status.

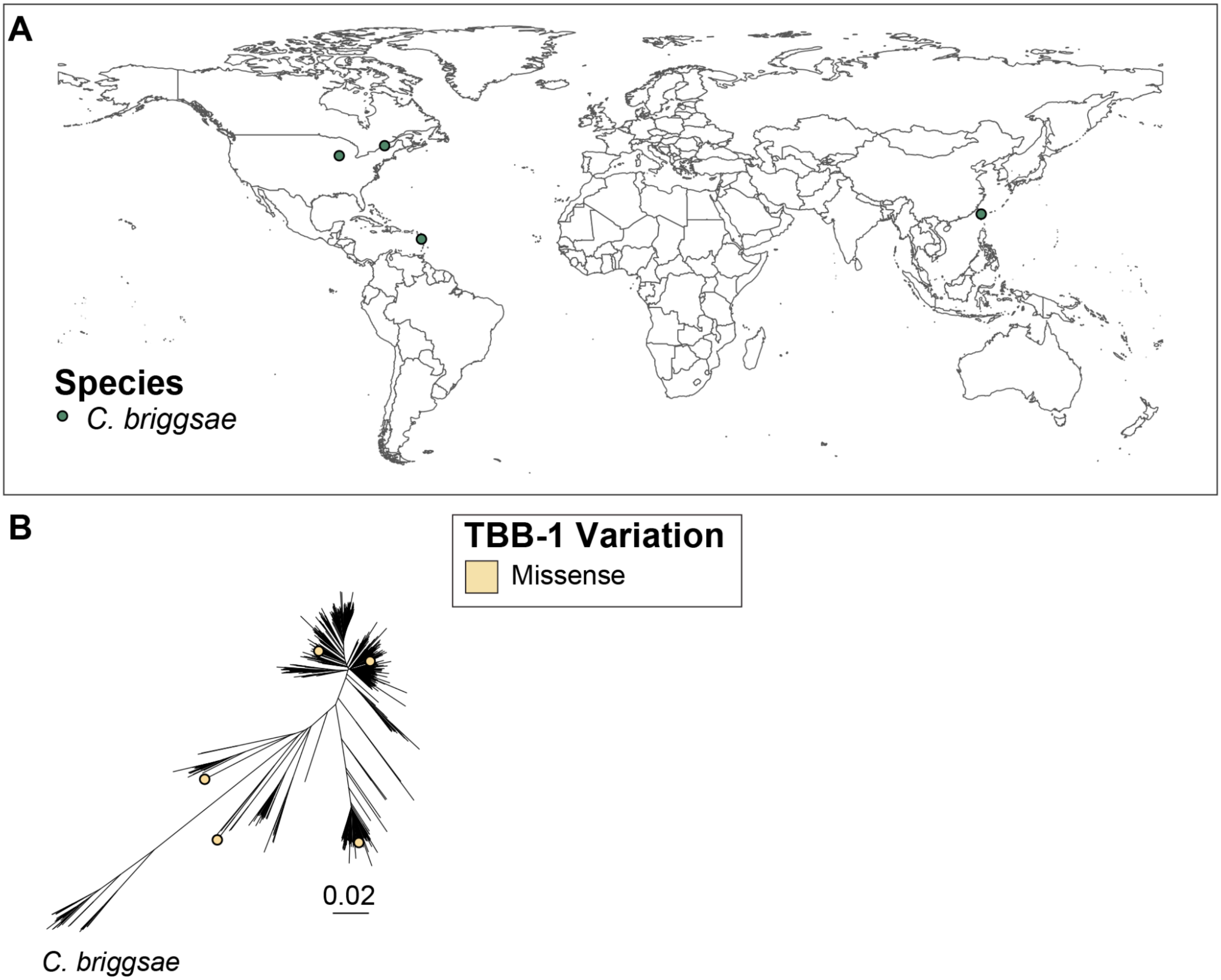


**S26 Fig. The global distribution of *Caenorhabditis* strains with predicted high-impact variation in *tbb-1***
Each point represents an isotype reference strain with a predicted high-impact variant in *tbb-1*. **(A)** Each point corresponds to the sampling location of the strain. **(B)** Each point corresponds to the location of the strain in a genome-wide phylogeny of 641 *C. briggsae* isotype reference strains. One isotype, XZ1213 has a high-impact *tbb-1* variant, but sampling coordinates were not recorded.

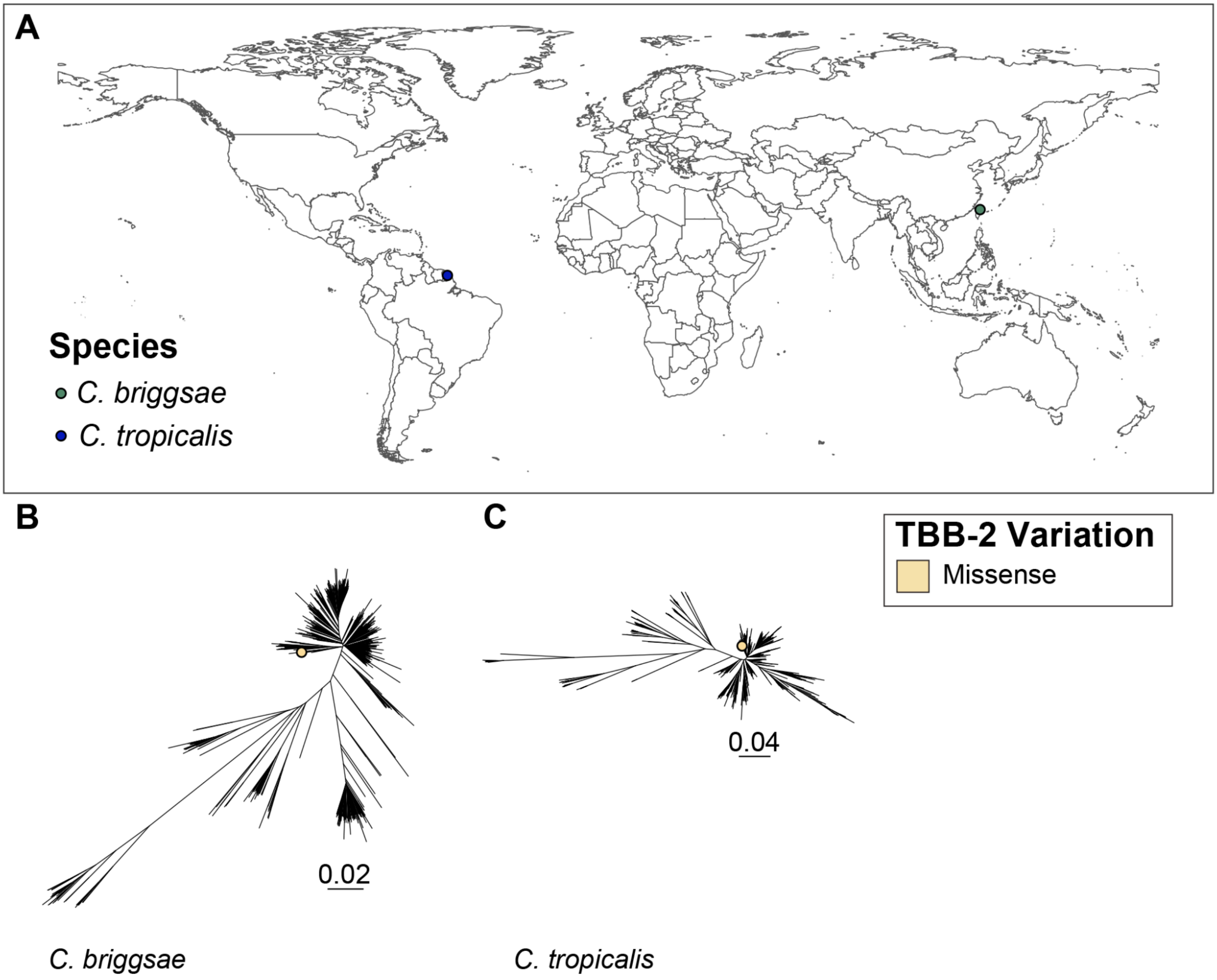


**S27 Fig. The global distribution of *Caenorhabditis* strains that contain predicted high-impact variation in *tbb-2***

Each point represents an isotype reference strain with a predicted high-impact variant in *tbb-2*. **(A)** Each point corresponds to the sampling location of the strain. Each point corresponds to the location of the strain in a genome-wide phylogeny of **(B)** 641 *C. briggsae* and **(C)** 518 *C. tropicalis* isotype reference strains.

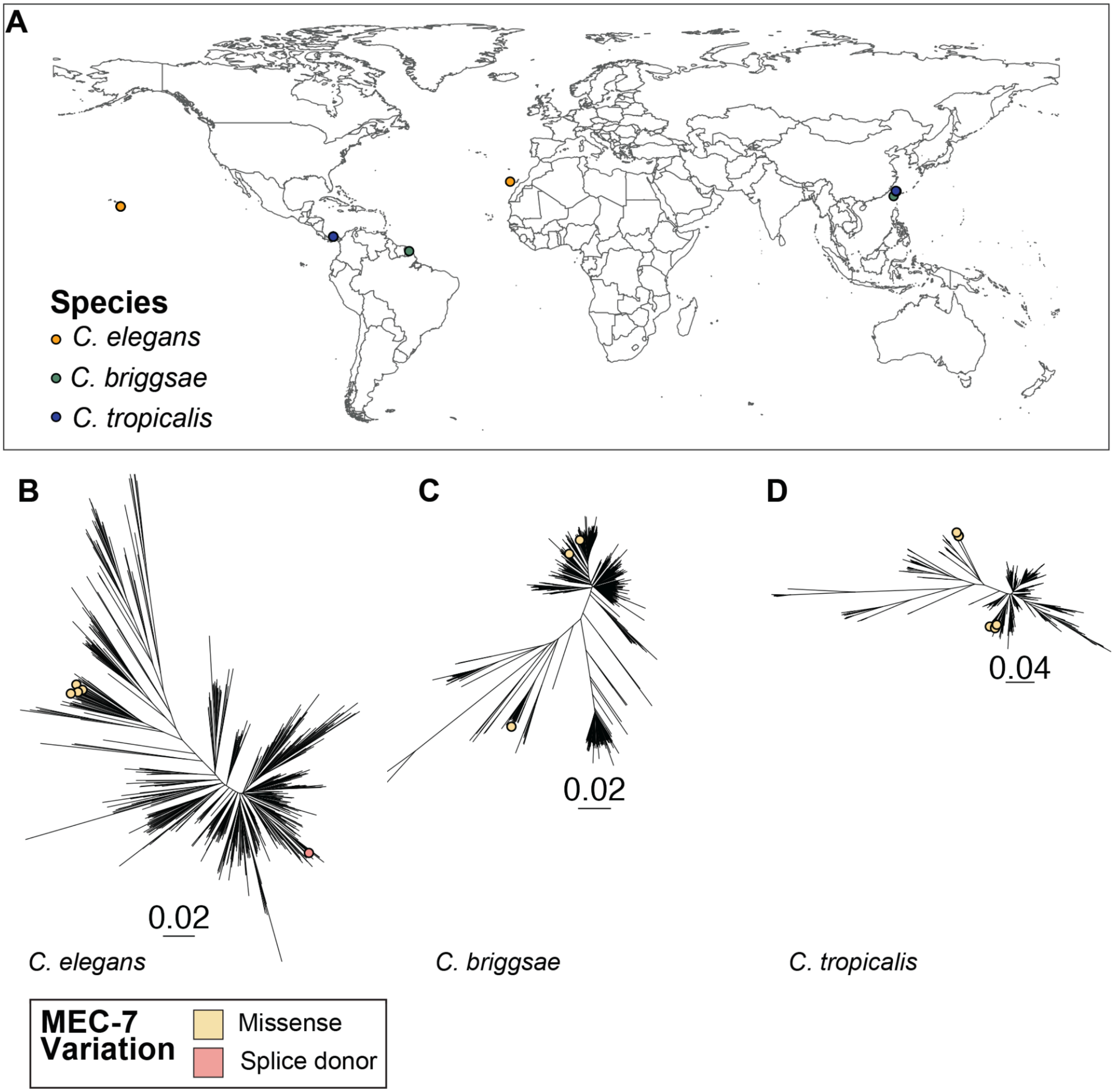


**S28 Fig. The global distribution of *Caenorhabditis* strains that contain predicted high-impact variation in *mec-7***

Each point represents an isotype reference strain with a predicted high-impact variant in *mec-7*. **(A)** Each point corresponds to the sampling location of an individual *C. elegans*, *C. briggsae*, or *C. tropicalis* strain. A genome-wide phylogeny of **(B)** 611 *C. elegans*, **(C)** 641 *C. briggsae*, and **(D)** 518 *C. tropicalis* isotype reference strains, where each point denotes an isotype reference strain with a predicted high-impact consequence in *mec-7*.

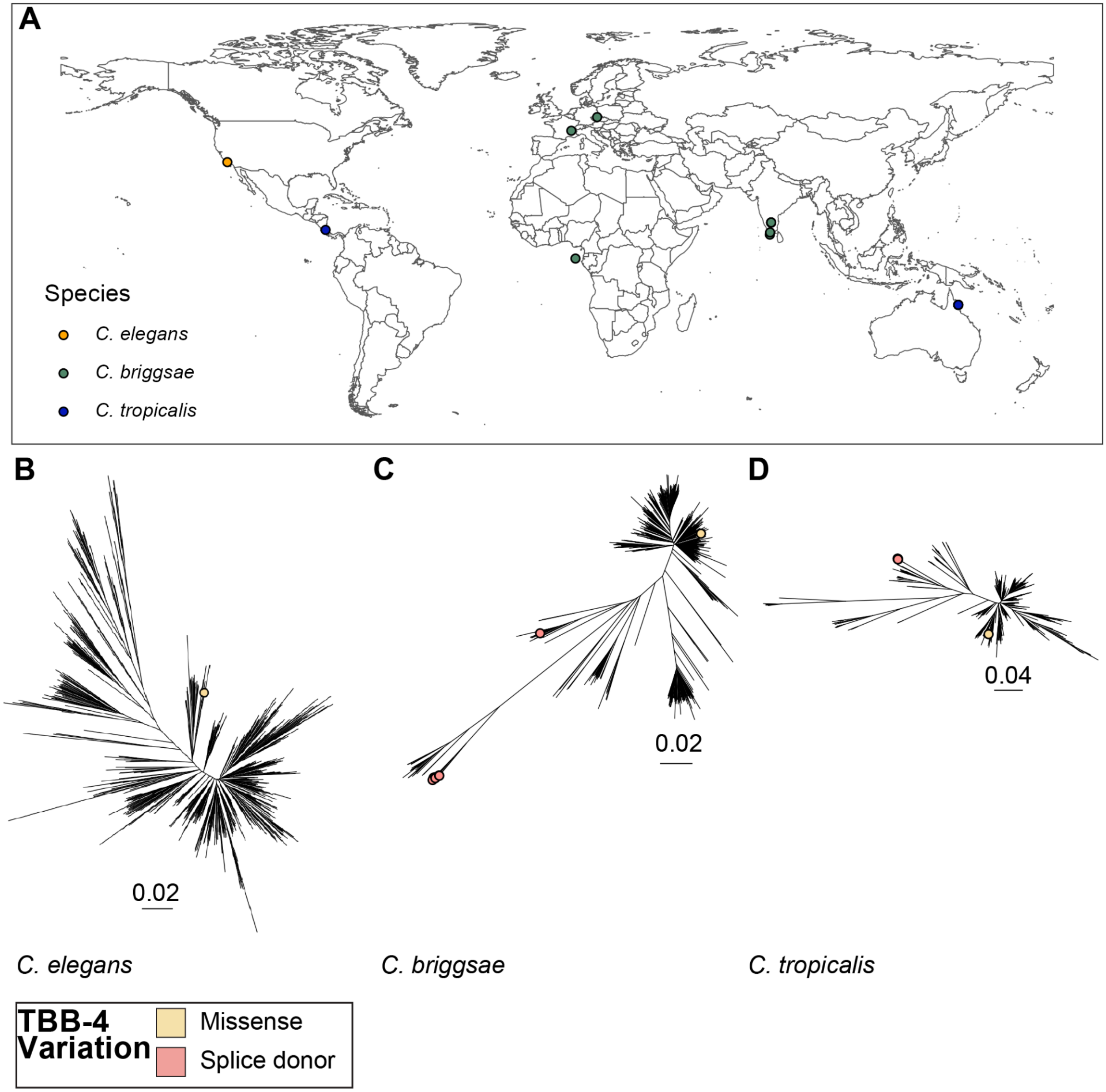


**S29 Fig. The global distribution of *Caenorhabditis* strains that contain predicted high-impact variation in *tbb-4*.**

Each point represents an isotype reference strain with a predicted high-impact variant in *tbb-4*. **(A)** Each point corresponds to the sampling location of an individual *C. elegans*, *C. briggsae,* or *C. tropicalis* strain with a predicted high-impact consequence in the gene *tbb-4*. Each point corresponds to the location of the strain in a genome-wide phylogeny of **(B)** 611 *C. elegans*, **(C)** 641 *C. briggsae*, and **(D)** 518 *C. tropicalis* isotype reference strains.

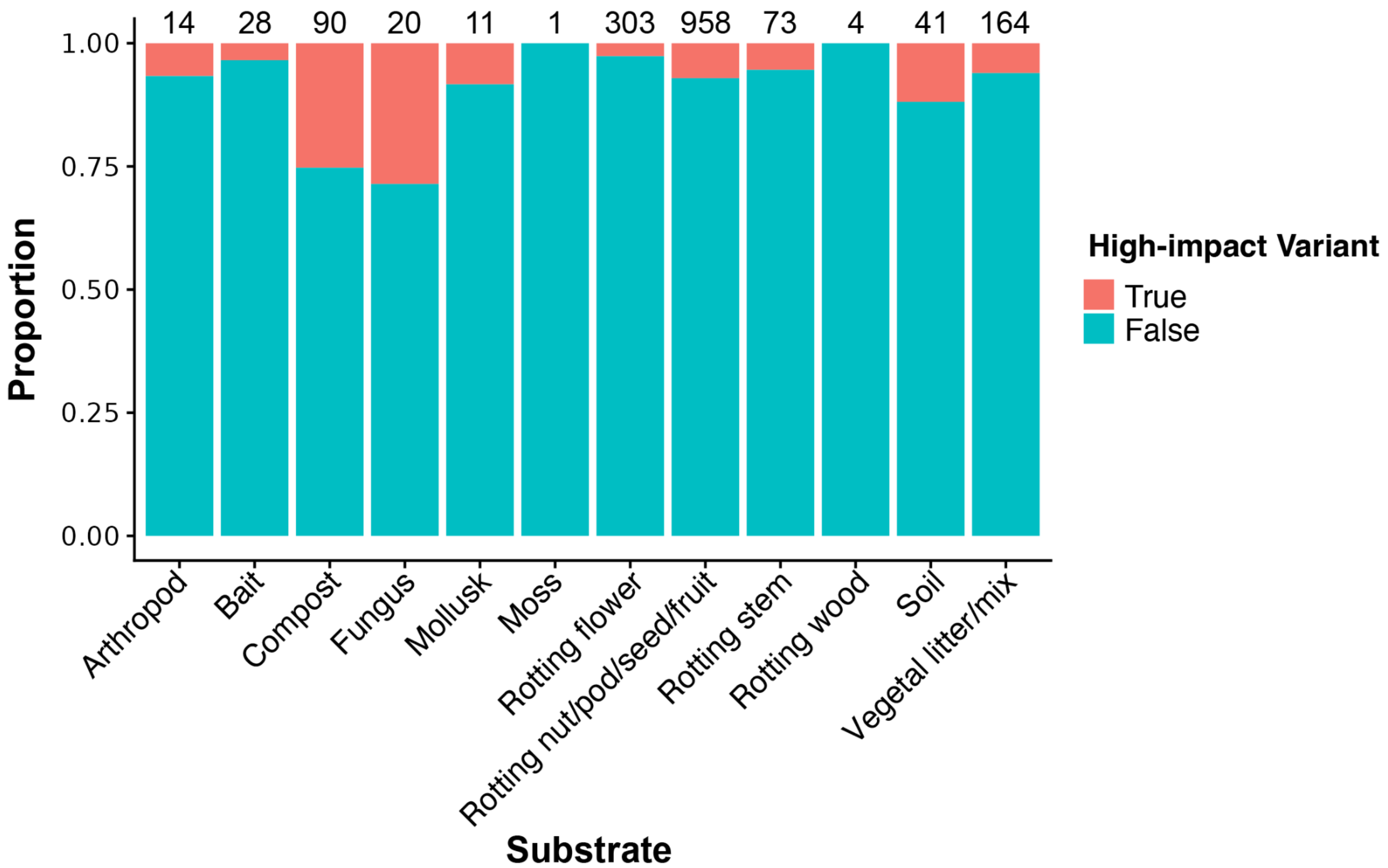


**S30 Fig. The proportion of strains with high-impact resistant variants in beta-tubulin genes and the substrates where those strains were found.**

The proportion of strains (y-axis) found in a given substrate (x-axis) are displayed. Strains with a high-impact variant in a beta-tubulin gene are colored salmon. Strains with no variants in a beta-tubulin gene are colored teal. The total number of strains isolated from a given substrate is displayed above each column. Moss and rotting wood were not included in the substrate enrichment analysis due to the small sample size. No significant relationship between beta-tubulin gene variant status and substrate were identified (Fisher's Exact Test, $p$=1).

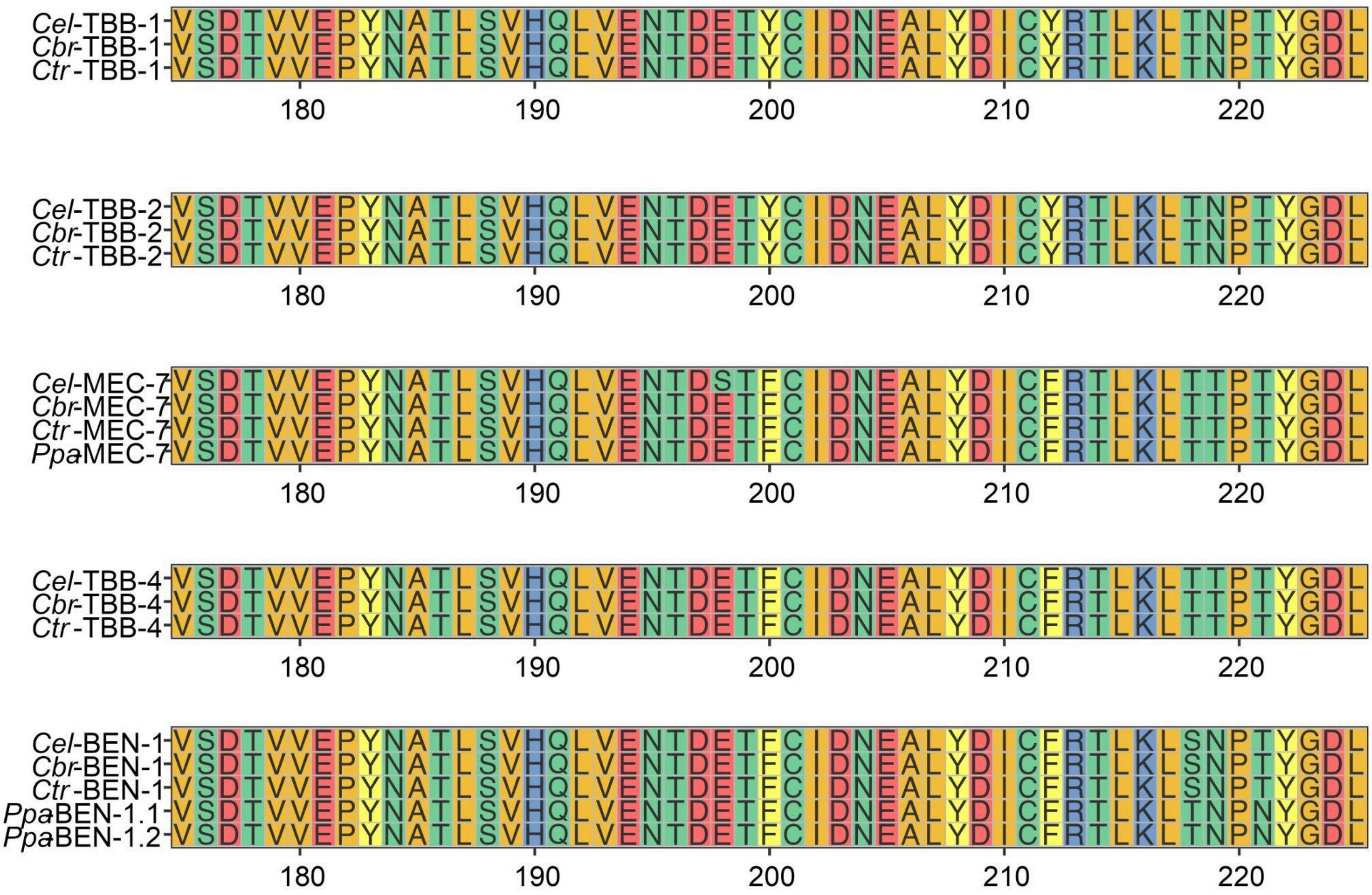


**S31 Fig. Multiple sequence alignment of TBB-1, TBB-2, MEC-7, TBB-4, and BEN-1 proteins from four free-living Clade V nematode species.**

Amino acid sequences of beta-tubulin isoforms TBB-1, TBB-2, MEC-7, TBB-4, and BEN-1 from *C. elegans* (*Cel-*), *C. briggsae (Cbr-)*, *C. tropicalis* (*Ctr-*), and *P. pacificus* (*Ppa-*) are aligned with MAFFT, and the alignment is displayed from amino acid residue 175 to residue 225. The region displayed is hypothesized to bind benzimidazoles. Residues are colored by side-chain chemical properties with the default *ggmsa* color scheme.
